# Parallels in leg and wing proximal-distal patterning in Diptera and Lepidoptera

**DOI:** 10.64898/2026.05.08.723717

**Authors:** Jeriel C. H. Lee, Tirtha Das Banerjee, Antónia Monteiro

## Abstract

Both insect wings and legs are appendages with a proximal–distal (PD) axis that is patterned during development, likely through a shared mechanism. Comparisons between the two types of appendages have been detailed in *Drosophila melanogaster*, a derived insect where both traits develop initially as imaginal discs. Here, we visualise and compare the expression of prominent PD developmental genes in the embryonic legs and larval wings of a lepidopteran species, *Bicyclus anynana*. We examine the domains of twelve appendage patterning genes that subdivide legs into distal, medial, or proximal domains (*homothorax, extradenticle, teashirt, dachshund, Distal-less, spineless, rotund, bric-à-brac, Bar, aristaless, Lim, and apterous)*, two genes that refine the PD axis after initial specification in both legs and wings (*fringe, brother of odd with entrails limited)*, and three morphogens (*wingless, decapentaplegic, and vein)*. Our results reveal high spatial congruence in the order of PD gene expression between the two appendages providing evidence for a partially co-opted leg-PD-patterning network during wing evolution. Notably, we observe a distinct loss of the medial domain in the *Bicyclus* wing compared to the leg, supported by the absence of broad *dachshund* expression, as also observed in flies. Further comparisons with *Drosophila* highlight additional conserved versus lineage-specific regulatory architectures. These findings suggest a deeply conserved PD patterning logic across Mecopterida, while pointing to divergent mechanisms that likely facilitated the morphological innovation of insect wings.

## INTRODUCTION

Novel complex traits often use core patterning mechanisms, or gene-regulatory networks (GRNs), that are also used to pattern other (more ancestral) traits in the same body. For example, petals in flowers use a modified leaf genetic program for their development (Sablowski, 2015), the novel turtle carapace uses a fused and restructured rib cage (Hirasawa *et al*., 2013), and new lobes in the male genitalia of *Drosophila* emerge from the same genetic program that produces breathing spiracles in larvae (Glassford *et al*., 2015). For insect wings, however, a trait with remarkable patterning complexity that evolved ∼400 million years ago, it remains unclear which ancestral GRN, if any, was reused to produce them. While wings are positioned dorsally in the body, they likely share patterning mechanisms with legs, as previous studies have shown that both appendages grow along a Proximal-Distal (PD) axis (Kojima, 2004; Ruiz-Losada *et al*., 2018). Key modifications to transform a tube-like leg into a sheet-like wing have also been proposed in *Drosophila*, which mostly involve wings recruiting *apterous* (*ap*), a distal leg PD patterning gene, into a novel dorsal compartment (Williams *et al*., 1994; Williams & Carroll, 1993).

The Proximal-Distal (PD) patterning of legs has been most extensively studied in *Drosophila* (Fig. 1), among other insects. In *Drosophila*, leg PD patterning starts with the juxtaposed expression of two ligands, Wnt family ligand Wingless (Wg/Wnt1) and TGF-β family ligand Decapentaplegic (Dpp), in ventral and dorsal domains of the imaginal disc, respectively (Lecuit & Cohen, 1997). These long-range morphogens induce the expression of a specific class of appendage patterning genes, called leg gap genes (LGGs): *Distal-less* (*Dll*), at the centre of the disc (distal end), and *dachshund* (*dac*), in a more medial domain. The expression of these genes represses another LGG, *homothorax* (*hth*), in the disc center, limiting *hth* expression to the disc periphery (proximally along the PD axis). Hth interacts with a Hox cofactor, Extradenticle (Exd), for nuclear localisation and the formation of an Hth-Exd heterodimer for cooperative DNA binding (Jaw *et al*., 2000). In turn, when *hth*, *dac*, or *Dll* expression is disrupted, a corresponding regional deletion or “gap” in the adult leg is seen (Rauskolb, 2001).

**Figure 1:**
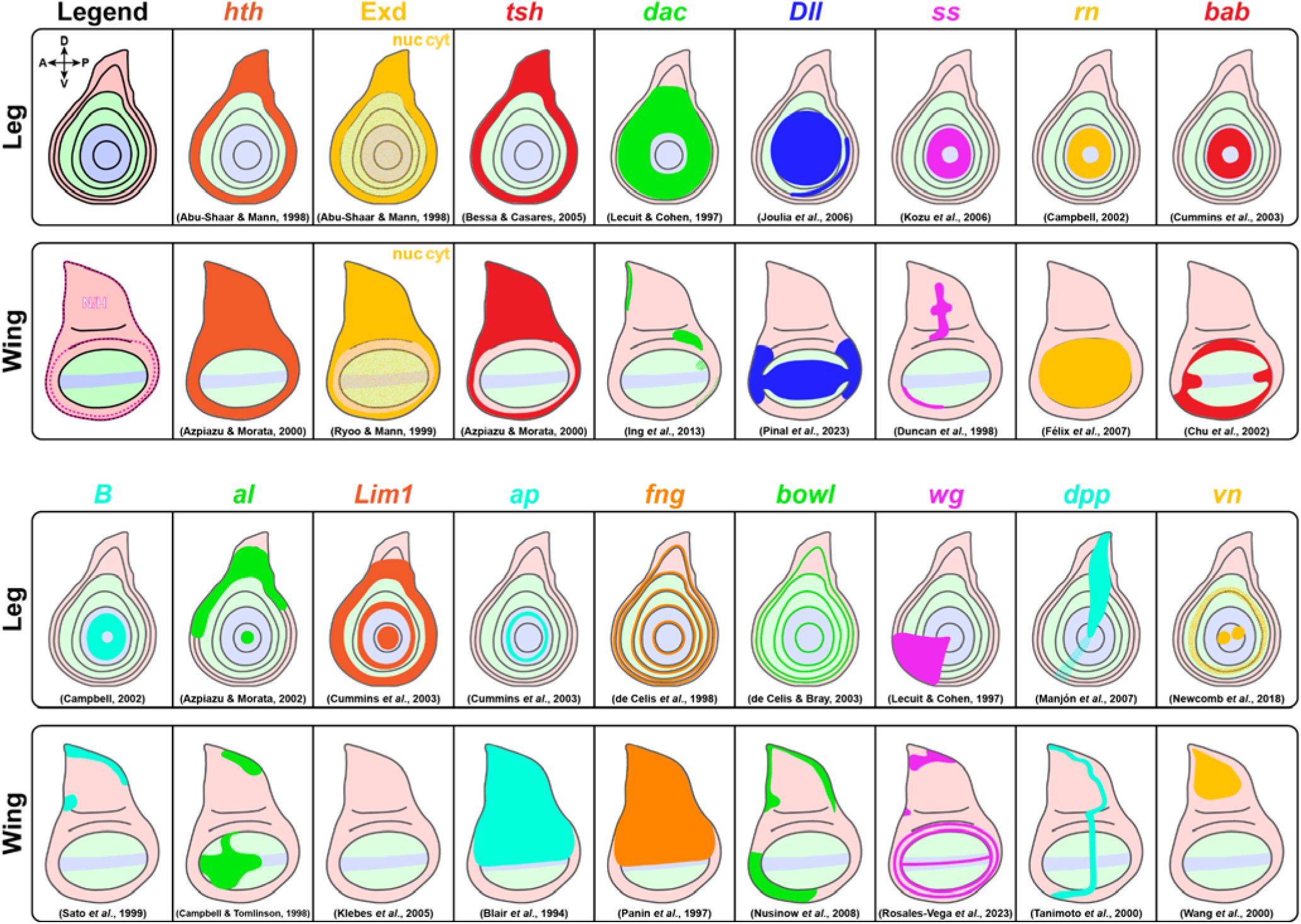
Review of APG expression domains in both *Drosophila* third-instar leg and wing imaginal discs. The PD domains in the leg and wing disc and the notum-hinge region (pink dashed outline; N/H) in the wing have been outlined in the first box (Red = Proximal; Green = Intermediate; Blue = Distal). The other axes of both appendages are specified at the top left of the leg legend (D = Dorsal; V = Ventral; A = Anter or; P = Posterior). Relative areas of expression are highlighted for each gene, with references below each illustration. For Exd, domains that have nuclear colocalisation (nuc) were represented with a solid colour, while cytoplasmic domains (cyt) were represented with stippling. Little to no *Lim1* expression was recorded in the *Drosophila* wing.

Outside *Drosophila*, Wg and Dpp gradients also establish the PD axis in insects such as *Tribolium castaneum* (Grossmann *et al*., 2009) and *Gryllus bimaculatus* (Niwa *et al*., 2000), in myriapods such as *Glomeris marginata* (Prpic, 2004), and in crustaceans such as *Triops longicaudatus* (Nulsen & Nagy, 1999). Although some lineage-specific modifications exist, the regionalization of the arthropod leg PD axis by these canonical LGGs is broadly conserved, with *hth*, *dac*, and *Dll* predominantly specifying proximal, intermediate, and distal territories, respectively (Abzhanov & Kaufman, 2000; Angelini & Kaufman, 2004; Inoue *et al*., 2002; Prpic *et al*., 2003; Prpic & Tautz, 2003; Sewell *et al*., 2008; Sharma *et al*., 2013). Studies on an arthropod outgroup, the onychophorans, suggest similar LGG patterning roles in unsegmented limbs (Janssen *et al*., 2010). Notably, while the hth/exd proximal co-expression domain is highly conserved throughout arthropods, the individual expression domains of these two genes have diverged (Angelini & Kaufman, 2005; Prpic & Telford, 2008).

In contrast to legs, where PD patterning has been extensively characterised in many insects, wing patterning studies lack comparable depth and diversity. In *Drosophila*, the activities of Wg and Dpp are still central to initiating and organising the distal development and growth of wings (Bosch *et al*., 2017; Swarup & Verheyen, 2012). Subsequently, several LGGs are expressed in wings in similar PD domains as in legs, such as *hth* (Azpiazu & Morata, 2000, 2002), *teashirt* (*tsh*) — another proximal LGG (Erkner *et al*., 1999; Wu & Cohen, 2002), and two distal gap genes, *rotund* (*rn*) (St Pierre *et al*., 2002), and *Dll* (Gorfinkiel *et al*., 1997), supporting the hypothesis that most animal appendages use a common shared PD patterning network (Panganiban *et al*., 1997). Despite these parallels, important differences between leg and wing patterning programs have also been documented (Fig. 1, Tripathi & Irvine, 2022; Williams *et al*., 1994), albeit never directly compared outside of *Drosophila*. Furthermore, comparative work on PD axis specification in wings outside of *Drosophila* remains scarce (Hanly *et al*., 2019; Hines *et al*., 2012).

Evidence that wings and legs share a fundamental appendage patterning program is supported by the ectopic expression of selector genes, which induce a change in appendage fate in *Drosophila*. *Dll* overexpression in the larval wing imaginal disc is sufficient to transform wings and halteres into leg-like appendages (Gorfinkiel *et al*., 1997), while ectopic *vestigial* (*vg*) expression in leg imaginal discs is sufficient to induce these to grow into flat, wing-like outgrowths (Kim *et al*., 1996). In addition, these ectopic outgrowths displayed proper PD axis specification, with fate-altered wings showing a femur-like proximal, tibia-like medial, and tarsi-like distal segments (Gorfinkiel *et al*., 1997), and fate- altered legs having a “wing” margin aligned with the distal tip of the modified leg (Kim *et al*., 1996). Hence, a common PD axis patterning program is likely underscoring these similarities.

Here, we test the hypothesis of a common PD leg-wing patterning program in insects by characterising the spatial-temporal expression of twelve prominent appendage patterning genes (APGs) in both embryonic thoracic legs and larval wings of the model butterfly *Bicyclus anynana* (*Bicyclus*). These APGs are further categorised as broad-domain APGs involved in proximal (*hth, exd*, and *tsh*), intermediate (*dac*), and distal patterning (*Dll*), and narrow-domain APGs involved mainly in the distal subdivision of the tarsus and pretarsus: *spineless* (*ss*), *rotund* (*rn*), *bric-à-brac* (*bab*), *Bar* (*B*), *apterous* (*ap*), *aristaless* (*al*), and *Lim1*. We also explored the expression domains of five other APGs, which include two genes involved in PD segment boundary refinement (joint formation) after initial PD axis specification by the broad-domain APGs, *fringe* (*fng*) (Rauskolb, 2001) and *brother of odd with entrails limited* (*bowl*) (de Celis Ibeas & Bray, 2003; Hao *et al*., 2003), as well as three crucial morphogens involved in establishing PD domain boundaries, *wg*, *dpp*, and *vein* (*vn*) (Estella *et al*., 2012).

## RESULTS

### Expression of broad-domain APGs in the embryonic leg

We first examined the expression of five broad-domain APGs: *hth*, *exd*, *tsh*, *dac*, and *Dll,* in 48 h embryonic thoracic legs (TLs). To characterise gene expression along the PD axis, we anatomically assigned the proximal leg region to include the coxa and trochanter, the intermediate region to comprise the femur and tibia, and the distal region to consist of the tarsus and pretarsus. All expression patterns were examined at the mRNA level using *in situ* hybridization (HCR).

These genes were expressed in slightly overlapping domains along the PD axis. For ***hth*,** we observed two distinct rings restricted to the proximal domain (Fig. 2A, A’). For ***exd***, baseline expression occurred in the entire leg, but it was slightly elevated in the proximal domain, presumably due to the modest upregulation of *exd* via Hth (Fig. 2B, B’, G) (Corsetti & Azpiazu, 2013). ***tio*** was broadly expressed in the entire proximal domain of the leg (Fig. 2C, C’, H). ***dac*** had a single, broad ring in the intermediate domain (Fig. 2D, D’, I). ***Dll*** had two separate rings: a broader one at the tip (distal domain) and a narrower one in the proximal/intermediate domain (Fig. 2E – I). The more distal ring of *hth* expression overlapped with the more proximal ring of *Dll* expression (Fig. 2F), and this region of overlap also expressed *tio* (Fig. 2H). *dac* was expressed as a broad ring within both *Dll* rings, with a small region of co-expression on both sides (Fig. 2I).

**Figure 2:**
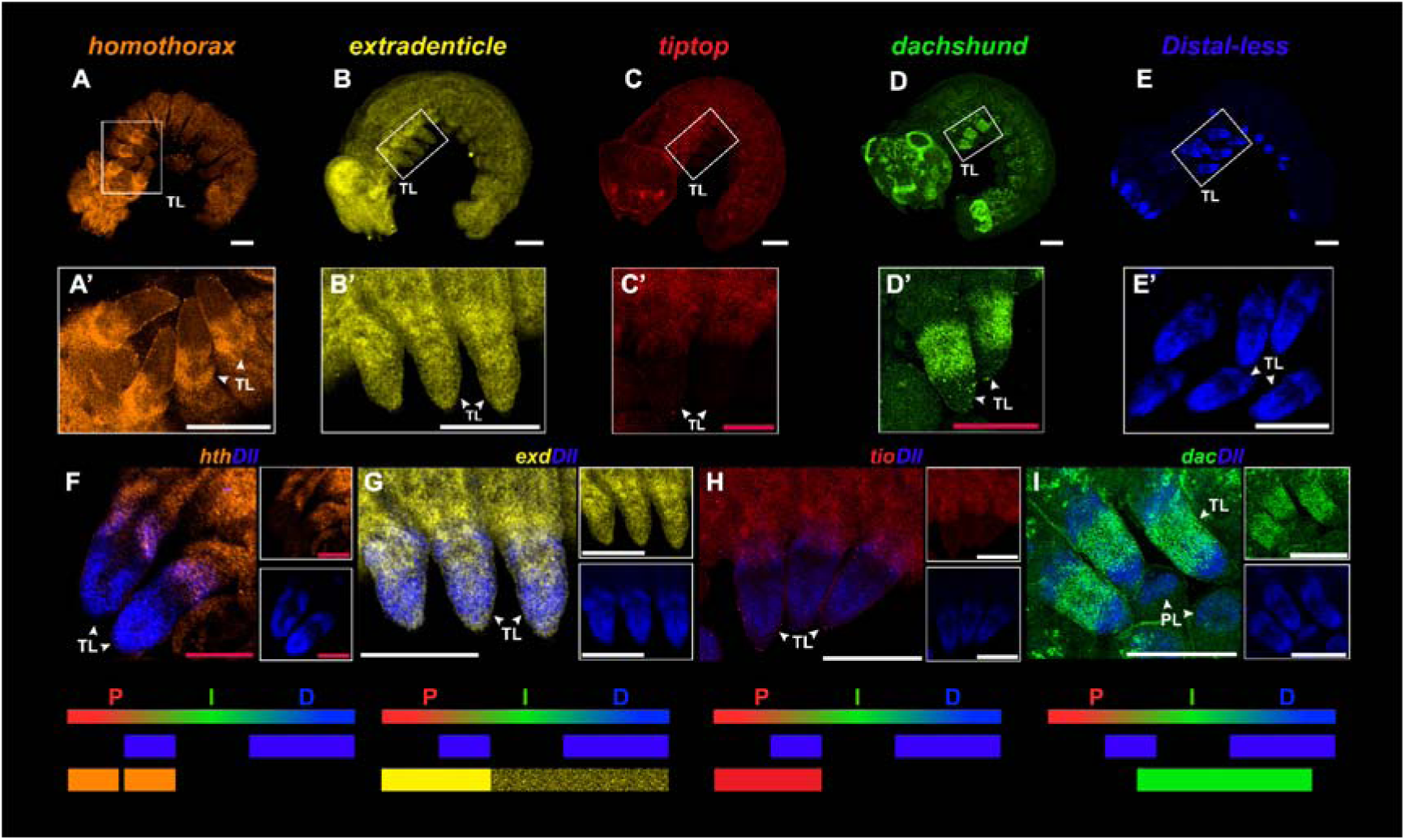
Expression of broad-domain APGs in thoracic legs (TLs) of *Bicyclus* 48 h embryos. (A’, B’, C’, D’, E’) are leg magnifications of **(A, B, C, D, E),** respectively (PL = prolegs). For each merged figure, a summary illustration is included at the bottom with an overview of the respective leg expression areas along the PD axis (Red domain = proximal, P; Green domain = intermediate, I; Blue domain = distal, D). **(A, A’)** *hth* is expressed as two rings in the leg proximal domain. **(B, B’)** *exd* staining shows uniform baseline expression throughout the leg, and a slightly higher expression level in the proximal region. **(C, C’)** *tiptop* (*tio*) is the single *tio/tsh* ortholog in lepidopterans (Zhang *et al*., 2020) and is expressed in the entire proximal region of the leg. **(D, D’)** *dac* expression is a single broad ring in the intermediate domain of the leg. **(E, E’)** *Dll* is expressed in two rings at the proximal-intermediate region and in the distal domain of the legs. *Dll* is also expressed strongly in the prolegs. (**F)** Single and merged channels of *hth* and *Dll* show overlap between the more distal ring of *hth* and the proximal ring of *Dll*. **(G)** Single and merged channels of *exd* and *Dll*. **(H)** Single and merged channels of *tio* and *Dll* showing overlap only at the proximal *Dll* ring. **(I)** Single and merged channels of *dac* and *Dll* show a small overlap between both ends of the *dac* ring and the two rings of *Dll*. All embryos are oriented from anterior to posterior (left to right). White scale bars: 100 µm; red scale bars: 50 µm.

### Expression of narrow-domain APGs in the embryonic leg

Next, we explored the expression of narrow-domain APGs, *ss*, *rn*, *bab*, *ap*, *BarH1* (*B-H1*), *al*, *Lim1*, in embryonic legs. *apterous-A* (*apA*) was used to represent *ap*, which is the only gene copy in *Drosophila* (Prakash & Monteiro, 2018). BLAST and cross-referencing with OrthoDB (v12.2) revealed *B-H1* to be the only genomic copy of *Bar* in *Bicyclus*. ***ss*** was expressed in a few cells in the anterior-proximal region on the leg epithelium (Fig. 3A, A’). ***rn*** was expressed along a broad domain that spanned the intermediate and distal regions of the leg (Fig. 3B, B’). ***bab*** was expressed sparsely in the distal domain of the leg (Fig. 3C, C’). ***B-H1*** was expressed strongly in a single broad distal ring (Fig. 3D, D’). ***al*** was expressed in a strong proximal ring, as well as in a much thinner ring just before the tip of the leg (Fig. 3E, E’). ***Lim1*** was expressed along four strong, thick rings throughout the PD domains (Fig. 3F, F’). Lastly, ***apA*** was not expressed in the leg epidermis (Fig. S2A – A’’). In addition, we also characterised the expression of the *apA* paralog, *apterous-B* (*apB*), and found no visible expression in the legs (Fig. S3A).

**Figure 3:**
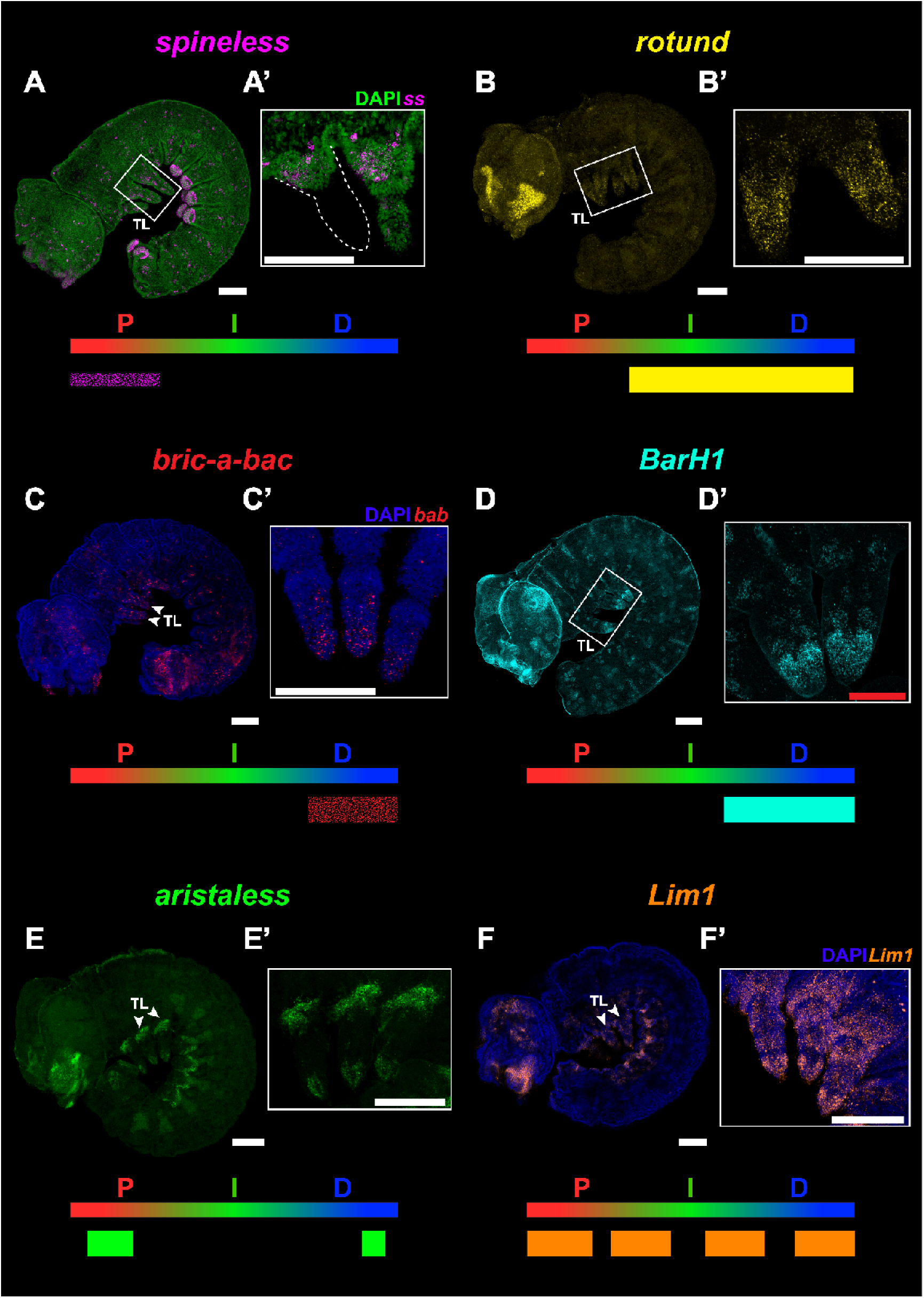
Expression of narrow-domain APGs in the thoracic legs (TLs) of *Bicyclus* embryos. (A’, B’, C’, D’, E’, F’) are leg magnifications of (A, B, C, D, E, F), respectively, with a summarised illustration for each gene. Leg magnifications (A’, C’) are maximum intensity projections that are restricted to the leg epithelium layer. Gene expressions that are not uniform rings around the leg epidermis are denoted by a thin bar, while sparse expression is represented by a speckled bar. (A, A’) *ss* is expressed at low levels in the anterior-proximal region of the leg epidermis. (B, B’) *rn* is expressed uniformly throughout the intermediate and distal regions of the leg. (C, C’) *bab* expression is sparse and only at the distal region of the leg epidermis. (D, D’) *B-H1* is expressed in a single broad distal domain. (E, E’) *al* is expressed in two rings at the proximal and distal regions. (F, F’) *Lim1* is expressed in four rings of similar thickness throughout the leg. All embryos are oriented from anterior to posterior (left to right). White scale bar: 100 µm; red scale bar: 50 µm.

### Expression of other APGs in the embryonic leg

In addition to characterising the expression of important PD patterning genes, we also explored *fng* and *bowl*, which refine the PD axis after broad-domain specification, and the morphogens *wg*, *dpp*, and *vn*, responsible for establishing the PD axis of the leg. Four rings of ***fng*** expression could be seen along all PD domains with different thicknesses (Fig. 4A, A’). ***bowl*** had a constitutive, low-level expression throughout the entire leg, except for two strong rings at the proximal-intermediate region (Fig. 4B, B’). ***wg*** had at least four short bands of strong expression throughout the ventral PD axis, interspersed by sparser expression (Fig. 4C, C’). A unique striped expression pattern of ***dpp*** was observed along the entire embryonic epidermis, including weak expression in the leg (Fig. 4D, D’). ***vn*** was present in the leg epidermal layer at the very tip (Fig. 4E – E’, F’’). These three morphogens only overlapped in epidermal expression at the very tip of the leg (Fig. 4F, F’).

**Figure 4:**
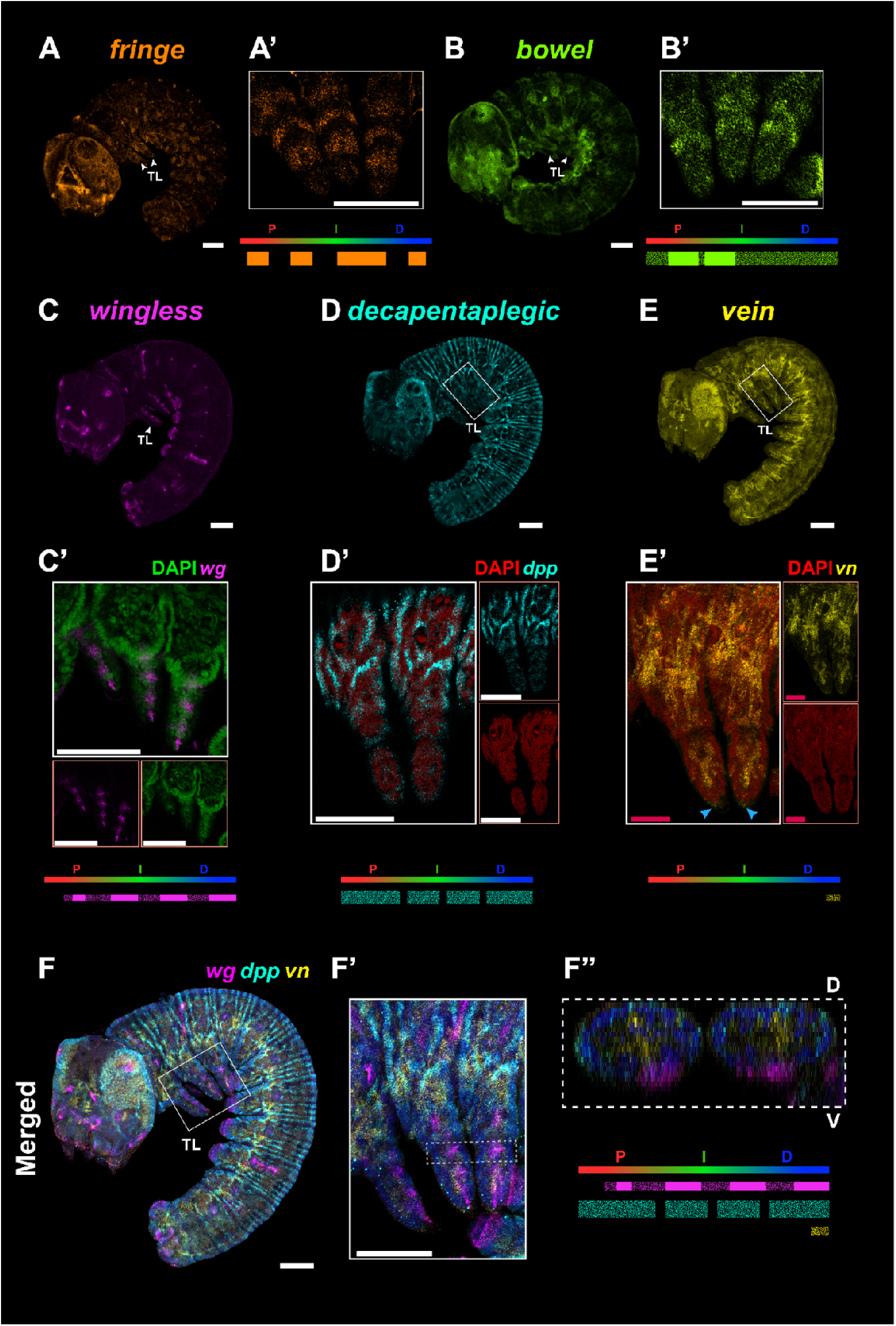
Expression of other APGs in the thoracic legs (TLs) of *Bicyclus* embryos. For each panel, a summarised illustration is included at the bottom. All embryo orientations, annotations, and summary illustration interpretations follow that of Fig. 2. **(A’, B’, C’, D’, E’)** are leg magnifications of **(A, B, C, D, E),** respectively. The leg magnifications **(C’, D’)** are maximum intensity projections restricted to the epithelium layer. **(A, A’)** *fng* is expressed in four rings of varying thickness across the three domains in the leg. **(B, B’)** *bowl* is expressed at a low level throughout the whole leg, with two stronger rings of expression in the proximal and intermediate regions of the leg. **(C, C’)** *wg* is expressed along the leg AP axis ventrally, with varying degrees of expression. **(D, D’)** *dpp* is expressed in a unique striped pattern sparsely in the leg. **(E, E’)** *vn* is expressed strongly in the mesodermal layer, and is only present weakly in the epidermis at the very tip of the leg (blue arrowheads). **(F, F’)** Merged image of *wg*, *dpp*, and *vn* morphogens. **(F’’)** Optical cross-section of merged leg outlined by a dashed box in (F’) showing *wg* present ventrally, *dpp* present sparsely throughout the leg epidermis, and *vn* only present in the middle mesodermal layer of the leg. All embryos are oriented from anterior to posterior (left to right). White scale bars: 100 µm; red scale bars: 50 µm.

### Expression of broad-domain APGs in the larval wing

We next investigated the expression of the APGs in fifth instar larval wings at three different developmental time points. To accurately track wing developmental stages, we used venation and tracheal growth markers, translated to a numerical system: early (0.00 – 0.75, EL), mid (1.00 – 1.75, ML), and late (2.00 – 2.75, LL) larval wing stages for temporal gene expression data (Banerjee & Monteiro, 2020b; Reed *et al*., 2007). All larval wings were imaged during the fifth instar.

Similar to the expression analysis in legs, we first explored the expression of broad-domain APGs in larval wings. ***hth*** was expressed in two strong bands in the proximal region, with a lower level of expression throughout the entire proximal domain, at all developmental stages (Fig. 5A – C). ***exd*** was expressed throughout the wing at all stages (Fig. 5D – F, Q). ***tio*** was expressed in a band that occupied most of the proximal domain at all stages (Fig. 5G – I). ***dac*** was expressed in two regions in the forewing and three in the hindwing (Fig. 5J – L). Both fore- and hindwings expressed *dac* at the proximal-most and anterior-proximal region. At the same time, *dac* was expressed in a cluster of intermediate cells at the intersection of the *Rs* and *M1* veins in the middle of hindwings alone (Fig. 5J – L, right wings). Expression of *dac* in all domains became strongest by mid-larval wing development. ***Dll*** was expressed in a band in the distal domain, along the wing margin but excluding the peripheral tissue, which will undergo apoptosis (Macdonald *et al*., 2010). *Dll* expression extended into the proximal domain at anterior and posterior regions, at all wing stages (Fig. 5M – O). *Dll* expression extended inward during LL (Fig. 5O) to form the centres of the future eyespots (Monteiro *et al*., 2013). There was overlap between the proximal expression of *Dll* and the more distal *hth* band (Fig. 5P, arrowheads), the expression domains of *tio* and *Dll* abutted each other (Fig. 5R), and *dac* expression overlapped with *Dll* at the anterior-proximal region (Fig. 5S, red arrowheads).

**Figure 5:**
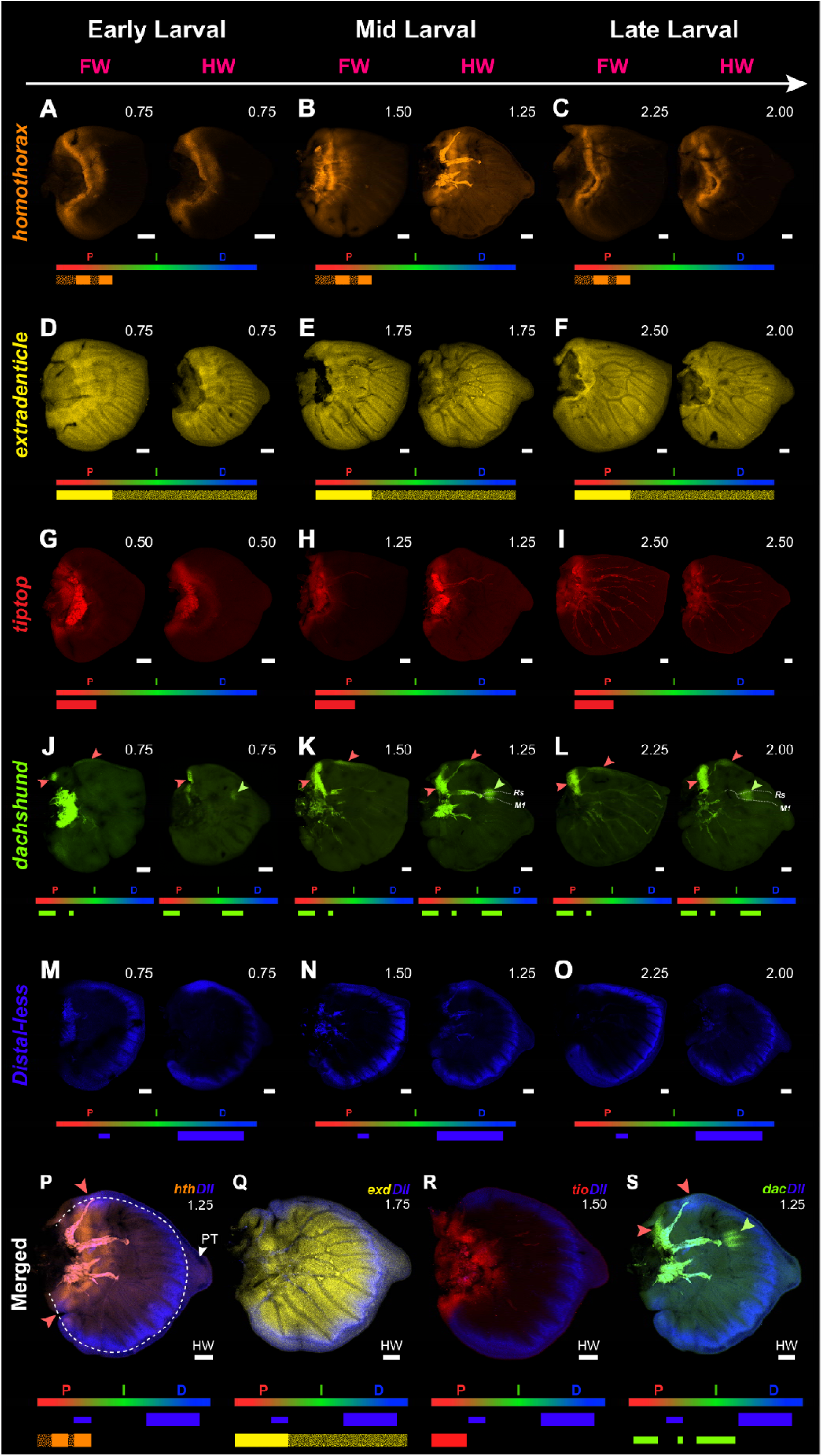
Spatial-temporal expression of broad-domain APGs in *Bicyclus* larval wings. Each panel consists of a forewing (FW; left) and a hindwing (HW; right). Wings are oriented along proximal-distal (left to right) and anterior-posterior (top to bottom) axes, and staged (top right). For each panel, a summarised illustration is included at the bottom (excluding tracheal/vein expression). Gene expressions that are not along a continuous AP band are denoted by a thin bar, while lower levels of expression are represented with a speckled bar. The wing margin is outlined with a white dashed line in subpanel P, while the peripheral tissue (PT) distal to the margin later undergoes apoptosis. **(A – C)** *hth* is expressed at the proximal domain for all wing stages. **(D – F)** *exd* expression is ubiquitous during wing development, with lower expression in the peripheral tissue. **(G – I)** *tio* is consistently expressed in the proximal region only. **(J – L)** *dac* varies across stages and wings. Red and green arrowheads represent expression in the proximal and intermediate domains, respectively. **(M – O)** *Dll* is consistently expressed along the wing margin at the distal region, and at both anterior and posterior ends of the proximal region throughout wing development. **(P – S)** Merged images of each gene with *Dll* in mid-larval hindwings. **(P)** *hth* and *Dll* colocalise at the anterior-most and posterior-most proximal region (red arrowheads). **(Q)** *exd* and *Dll* colocalise at the wing margin. **(R)** *Dll* expression stops just before *tio* expression in the proximal region. **(S)** Colocalization of *dac* and *Dll* in the anterior-proximal region (red arrowheads). All proximal tracheal and vein signals have been excluded from the summary illustration. Scale bar: 100 µm.

### Expression of narrow-domain APGs in the larval wing

Next, we characterised the expression of the narrow-domain APGs in fifth instar larval wings. Two small, distinct clusters of ***ss*** were observed at the proximal-most region across all larval wing stages (Fig. 6A – C). *ss* also has cell-type specific expression in discrete clusters at the wing margin, which are more visible in the mid and late larval wing (Fig. 6C). ***rn*** was consistently expressed in a broad domain spanning the intermediate and distal regions of the wing at all stages, with expression in the intermediate region being strongest (Fig. 6D – F). Multiple clusters of ***bab*** were seen throughout the three PD domains, with its onset in the early larval stage marked by strong localisation in a central cluster and at the wing tip. Peak *bab* expression was observed by the mid-larval stage, characterised by a central-proximal region and a triad of clusters in the intermediate domain. Comparison between the wing pairs revealed that the anterior-intermediate cluster was consistently more prominent in the forewing than in the hindwing (Fig. 6G – I). ***apA*** was expressed at a consistent baseline level throughout the dorsal side of the wing at all stages, as well as in a strong proximal band in all wings and at the anterior-distal region of all forewings (Fig. 6J – L). In contrast, while *apB* had the same expression domains as *apA*, the expression of this paralog in both fore and hindwings was more muted compared to *apA* (Fig. S3B; Prakash & Monteiro, 2018). Two short, anterior bands of ***al*** were seen in the intermediate and distal domains of the wing at all stages. The distal band in forewings crossed the AP boundary, while the same band in hindwings did not. The intermediate band in hindwings also seemed slightly shorter than that of the forewing at each developmental stage (Fig. 6M – O). ***B-H1*** and ***Lim1*** were not expressed in the wing throughout its development (Fig. S2B, C).

**Figure 6:**
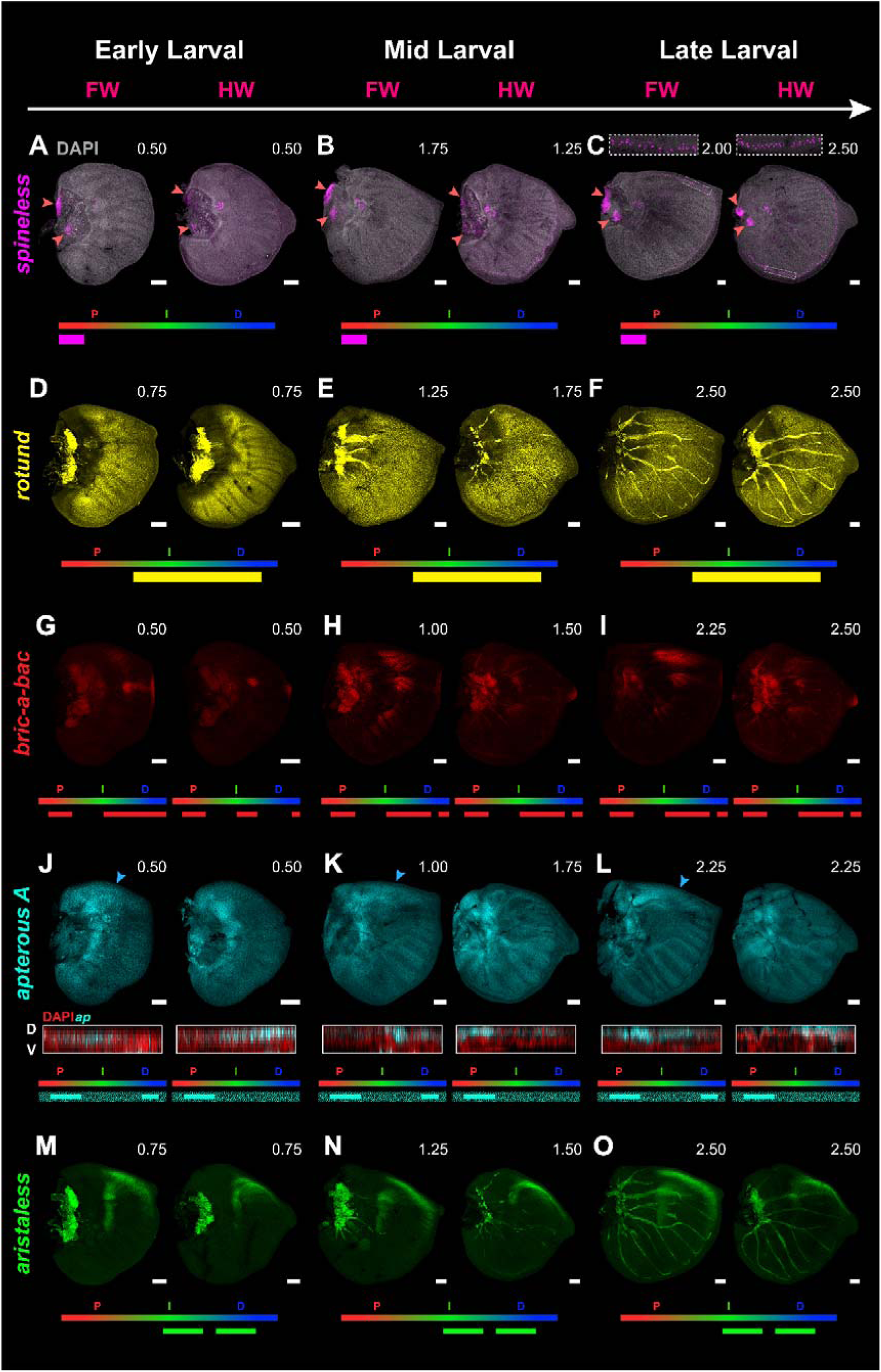
Spatial-temporal expression of narrow-domain APGs in *Bicyclus* larval wings. All wing orientations, annotations, and summary illustration interpretations follow that of Fig. 5. **(A – C)** *ss* is expressed in two relatively small cell clusters located at the proximal-most region (red arrowheads). A closeup of *ss* expression at the wing margin is shown for the late larval wings (white dashed boxes). This distal cell-type-specific *ss* expression is excluded from the summary illustration. **(D – F)** *rn* is expressed broadly in the entire intermediate and distal region except for the peripheral tissue **(G – I)** *bab* is expressed in one cluster of cells in the proximal region, up to three clusters in the intermediate region as the wing reaches mid-larval development, and at the distal tip in the peripheral tissue. **(J – L)** *apA* is expressed only in the dorsal cells of the wing, as shown in the DV cross-section below each wing. In forewings, strong *apA* expression is also seen in the anterior-dorsal region (blue arrowheads). **(M – O)** *al* is expressed consistently throughout larval wing development as a short band at the intermediate region and at the anterior-distal region. All proximal tracheal and vein signals have been excluded from the summary illustration. Scale bar: 100 µm.

### Expression of other APGs in the larval wing

We then looked at the expression of five other APGs in larval wings. ***fng*** was expressed in a single, thin band along the distal region separating the wing margin from the peripheral tissue, in the wing hinge, and along the veins across all stages. *fng* expression along the veins also became stronger in older wings (Fig. 7A – C). Constitutive ***bowl*** expression was observed at low levels throughout the entire wing at all stages, with a stronger proximal band (Fig. 7D – F, red arrowheads) and expression at the intersection between the *M3* and *Cu1* veins. In addition, forewings alone had an anterior-distal region of strong expression by mid- larval development (Fig. 7D – F, blue arrowheads). Consistent and strong ***wg*** expression was present throughout the peripheral tissue, in two thin proximal bands (one in the dorsal, the other in the ventral surface), in a cross-vein in the intermediate region of both fore- and hindwings, and in the anterior-proximal region of forewings (Fig. 7G – I). ***dpp*** was expressed in a strong band along the AP boundary, which splits into two at the distal region of forewings, in a more posterior band at earlier stages, in a short transversal band in the central-intermediate region of both wings, and flanking the posterior veins during the early larval stage before extending to the anterior veins during the late larval stage (Fig. 7J – L; Banerjee & Monteiro, 2020). Lastly, ***vn*** was strongly expressed in all intervein regions throughout all PD domains (Fig. 7M – O) and was expressed in a largely complementary pattern to that of *dpp* throughout wing development (Fig. 7P, Q). In older wings, *vn* was refined to the eyespot centres.

**Figure 7:**
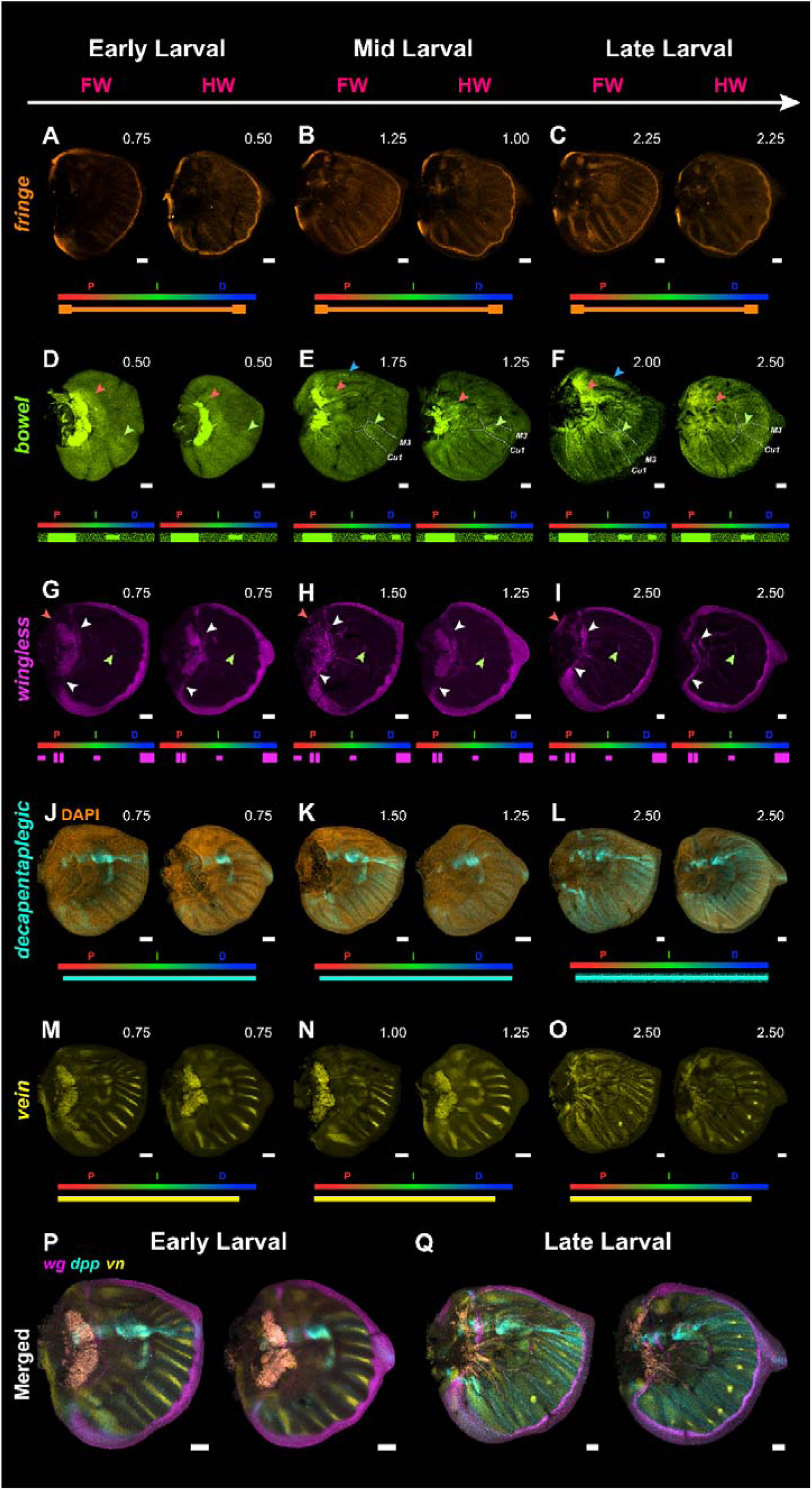
Spatial-temporal expression of other APGs in *B. anynana* larval wings. All wing orientations, annotations, and summary illustration interpretations follow that of Fig. 5. Red, green, and blue arrowheads indicate expression in the proximal, intermediate, and distal domains, respectively. **(A – C)** *fng* is expressed at the proximal-most region and along the wing margin, and in the veins. **(D – F)** *bowl* is sparsely expressed throughout the wing, with strong expression areas at all three domains of the larval wing. **(G – I)** *wg* expression is present at all three domains throughout larval wing development: in the most peripheral tissue along the margin of the disc, in a thin proximal band that is present in both the dorsal and ventral layers (white arrowheads), and on top of a future cross-vein (green arrowhead). **(J – L)** *dpp* is expressed strongly along the AP boundary from proximal to distal throughout all larval stages, along a more posterior domain, and in an orthogonal short stripe in the center of the wing at earlier stages. *dpp* flanks the veins at later stages. **(M, N, O)** *vn* is expressed in the central intervein regions throughout the entire wing and also at the proximal-most region during all larval stages. Distal expression becomes restricted to the eyespots during late wing development. **(P, Q)** Merged images of *wg*, *dpp* and *vn* in early (G, J, M) and late (I, L, O) wings. All proximal tracheal and vein signals have been excluded from the summary illustration. Scale bars: 100 µm.

## DISCUSSION

In this study, we tested whether genes that pattern the proximal-distal (PD) axis of legs are deployed similarly in wings. We characterised the spatial expression of twelve appendage patterning genes (APGs) and five additional APGs in 48 hr embryonic thoracic legs and across three developmental stages of fifth instar larval wings. Figure 8 summarises relative expression domains in *Bicyclus* legs and wings and compares them to *Drosophila*.

**Figure 8:**
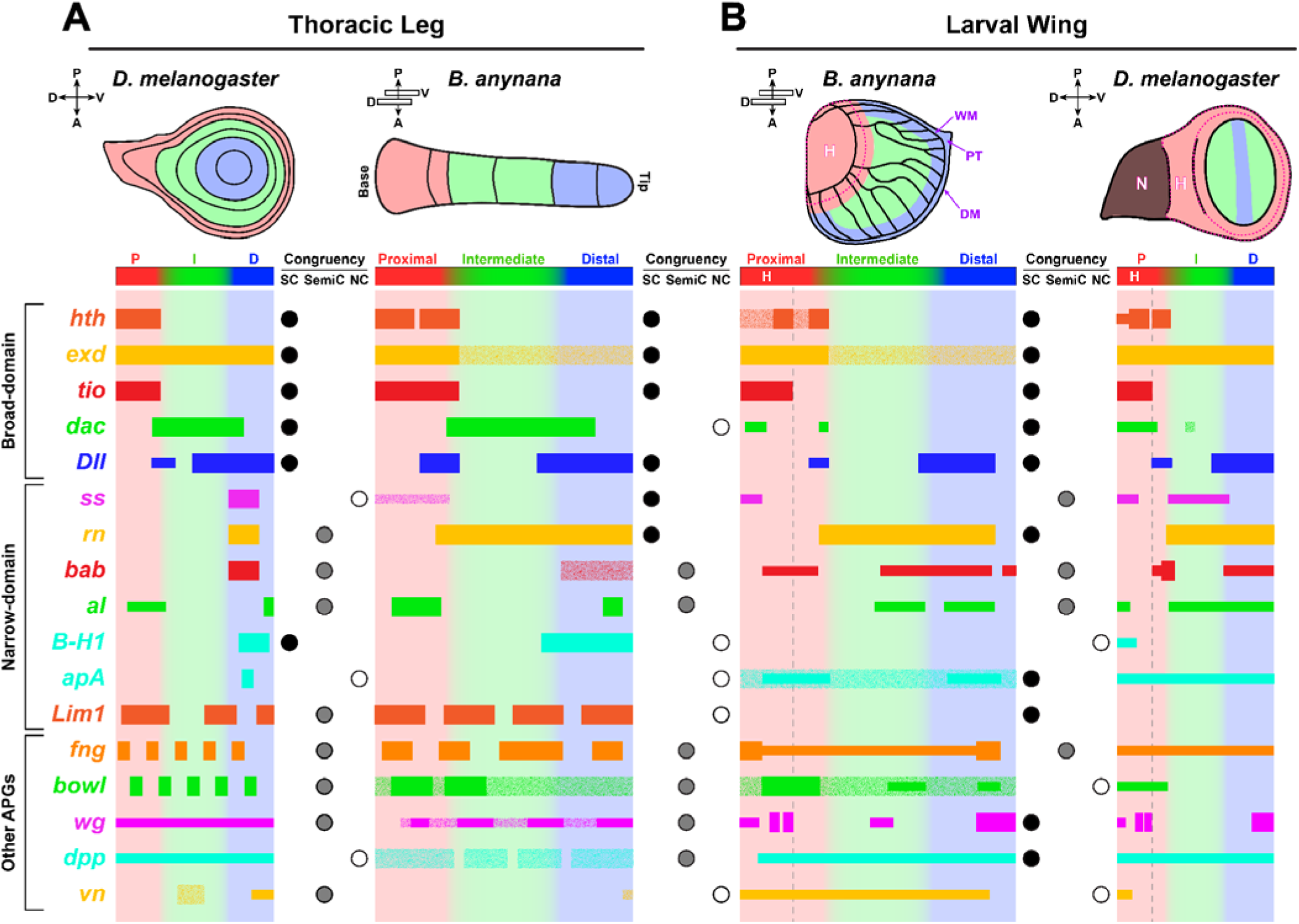
Overall summary of APG expressions in the 48hr embryonic thoracic leg and larval wing of *Bicyclus* and 3^rd^ instar larval imaginal discs of *Drosophila*. For all appendage illustrations and summary bars, red, green, and blue regions correspond to the proximal, intermediate, and distal domains, respectively. The length and co-expression domains of the summarized gene expressions are approximate. The other axes of each appendage are specified at the top left (D = Dorsal; V = Ventral; A = Anterior; P = Posterior). Genes that are not expressed as a ring around the leg epidermis or along the AP axis in larval wings are denoted with a thin bar, while lower expression levels in both appendages are represented by a speckled bar. For *Drosophila* comparisons, the presence of gene expression in the proximal (red), intermediate (green), and distal (blue) domains is denoted with P, I, and D, respectively. The concentric ring-format gene expression data of Figure 1 have been mapped onto a linear axis: distal domains represent the central domains, proximal domains represent the peripheral domains. (A) Relative areas of gene expression in the *Bicyclus* 48h leg (right) and late third instar *Drosophila* leg imaginal disc (left). (B) Relative areas of gene expression in the *Bicyclus* mid-larval forewing (left) and the *Drosophila* late third instar wing imaginal disc (right). The presumptive *Bicyclus* hinge region (H) and *Drosophila* hinge are outlined with pink dashed lines, while elements of the peripheral region were annotated with purple arrows (WM = Wing margin; PT = Peripheral tissue; DM = Disc margin). The *Drosophila* notum (N), represented by a dark brown area, is not included in the summary. The spatial congruency of all the genes between *Drosophila* and *Bicyclus* legs and larval wings is shown in the coloured circles between the summaries (black: Strongly congruent, SC; grey: Semi-congruent, SemiC; white: Non-congruent, NC).

### Defining the PD domains in *Bicyclus* embryonic leg and larval wing

We defined PD domains using conserved arthropod markers (Fig. 8B). We used *hth* to mark the proximal domain (Coulcher *et al*., 2015; Dong *et al*., 2001; Ruiz-Losada *et al*., 2018; Schaeper *et al*., 2013; Singh *et al*., 2019) and *Dll* to mark the distal domain (Jockusch *et al*., 2004; Tripathi & Irvine, 2022). In legs, the region between hth and Dll corresponds to the intermediate domain.

In wings, defining PD boundaries is less straightforward because of wing morphology and limited comparative data. We again used *hth* and *Dll* to define the proximal and distal domains, and *wg,* together with *hth,* and *tio/tsh* to identify the hinge (Bessa *et al*., 2009; Rosales-Vega *et al*., 2023). The two proximal *wg* bands of *Bicyclus* likely correspond to the inner and outer *wg* rings that define the *Drosophila* hinge (Fig. 7G – I; Rosales-Vega *et al*., 2023). With these markers, the remaining wing tissue was designated intermediate. This mapping places proximal and distal domains adjacent at the anterior and posterior margins (Fig. 8B), while the intermediate domain occupies a more central region that does not extend to the margins.

Despite broad correspondence between *Drosophila* and *Bicyclus* PD markers, we observed substantial heterochrony, particularly in vein development. In *Bicyclus* larval wings, *dpp* and *vn* patterns resemble later (pupal) *Drosophila* stages when these genes are expressed in veins (de Celis, 1997; Simcox *et al*., 1996; Sotillos & Celis, 2005; Sotillos & de Celis, 2006; Wessells *et al*., 1999). Nonetheless, a comparative study with the *Drosophila* larval appendages remains the most informative basis for cross-species comparison of most APG PD patterning functions.

### The majority of the APGs are expressed in similar domains across the PD axis in legs and larval wings of *Bicyclus*

We grouped APGs by spatial congruency between legs and wings (Fig. 8, central two columns): strongly congruent, semi-congruent, and non-congruent, based on relative position and approximate signal strength along the PD axis. Six APGs (*hth*, *exd*, *tio*, *Dll*, *ss*, and *rn*) are strongly congruent between Bicyclus embryonic legs and wings; this group includes four out of five broad-domain APGs. Two APGs (*bab* and *al*) are semi-congruent. *bab* is distal in both appendages but has additional proximal and intermediate wing expression. *al* is expressed at the distal tip in both appendages, but also shows proximal expression in legs and intermediate expression in wings. Four APGs, *dac*, *apA*, *Lim1,* and *B-H1*, are non-congruent. *dac* is strongly expressed in the intermediate leg domain, but is absent from the forewing intermediate domain and reduced to a small intermediate cluster in hindwings. *apA* is not expressed in the leg epidermis, while *Lim1* and *B-H1* are not expressed in wings (Fig. S2B, C). These divergences are consistent with a partial recruitment of a leg-PD-patterning GRN during wing evolution.

### Only proximal and distal PD patterning genes in the embryonic legs are likely reused to pattern the larval wing

Among strongly congruent genes, most are associated with proximal or distal patterning in *Drosophila,* with *exd* and *rn,* as exceptions: both span intermediate regions in legs and wings, but *exd* function is restricted to proximal identity (Rauskolb *et al*., 1995) and *rn* to distal patterning (St Pierre *et al*., 2002).

*tio*/*tsh* provides a clear example of conserved proximal expression across legs and wings in both *Drosophila* and *Bicyclus*. However, *tio*/*tsh* domains and functions vary widely across arthropods, including distal or broad appendage roles (Herke *et al*., 2005; March *et al*., 2018; Medina-Jiménez *et al*., 2021), suggesting that proximal-exclusive deployment arose within Holometabola. Within Lepidoptera, *tio* also contributes to both proximal and distal appendage development in *Bombyx mori* (Zhang *et al*., 2020), reinforcing this variability.

Leg-wing divergence in both *Drosophila* and *Bicyclus* is most evident in *wg* and *dpp* deployment in the wing, and in the exclusion of *dac* from most of the wing intermediate region. *dac* is the only broad-domain APG that lacks a comparable broad wing band. In *Drosophila*, *dac* is expressed in the notum/body wall, and in a relatively small proximal-intermediate domain in the wing proper compared to its leg domain. Interestingly, *dac* repression promotes leg-to-wing transdetermination and activation of wing selectors (Ing *et al*., 2013). In that context, ectopic dorsal *wg* expression in the leg pouch enables broad *dpp* expression, represses *dac*, and permits *vg* activation. Thus, loss of a continuous *dac* domain may be a prerequisite for wing specification.

Although most strongly congruent genes are conserved between *Drosophila* and *Bicyclus*, *ss* and *rn* differ between the species. In *Bicyclus ss* expression in wings and legs resembles *ss* expression in *Drosophila* antennae, rather than legs (Setton *et al*., 2017): *ss* is proximal in *Bicyclus* legs and wings but distal in *Drosophila* legs. In *Drosophila*, *ss* is activated by Dll (D. M. Duncan *et al*., 1998) while being proximally repressed by *Sex combs reduced* in the T1 (Tsubota *et al*., 2008), and Antennapedia in the T2/T3 (D. Duncan *et al*., 2010). Such repression may be absent in *Bicyclus* because sparse *ss* overlaps *hth* expression. Since *hth* activates *ss* in *Drosophila* antennae, this interaction may be conserved in *Bicyclus* legs and wings. *ss* expression in sensory organs, however, appears conserved (Fig. 5A – C; Emmons *et al*., 2007).

*rn* is more congruent between *Bicyclus* legs and wings than in *Drosophila*: both species express *rn* in intermediate and distal wing domains, but *rn* is restricted to the distal domain in *Drosophila* legs. Despite these differences, PD patterning genes generally show similar PD ordering within each organism.

### Conservation and novelty of PD patterning in legs and wing via changes in the deployment of *wg*, *dpp,* and *vn* morphogens

In *Drosophila* leg and wing discs, PD patterning relies heavily on Wg and Dpp gradients. In legs, Wg and Dpp are expressed in opposing ventral and dorsal stripes (Fig. 1) and jointly activate *Dll* at the disc centre, while repressing *dac* and *hth* (Lecuit & Cohen, 1997). At lower Wg/Dpp levels, *dac* is permitted in an intermediate domain, and Dll directly promotes *dac* in part of its distal domain (Estella *et al*., 2012; Giorgianni & Mann, 2011), while *hth* is restricted proximally (Abu-Shaar & Mann, 1998; Wu & Cohen, 1999). In wings, *wg* is high along the DV margin and periphery, and *dpp* is strong along the AP boundary; together they promote distal programs and repress proximal programs, although wing-specific factors (e.g., *vg* and *scalloped*) have stronger roles in distal domain specification (Tripathi & Irvine, 2022). Proximal and distal APG order in legs is conserved between *Drosophila* and *Bicyclus* despite strong differences in *dpp* expression (Fig. 8). The interrupted *dpp* stripe in *Bicyclus* more closely resembles patterns reported in other insects such as *Tribolium* and *Gryllus* (Niwa *et al*., 2000; Pechmann & Prpic, 2022). This variability may not affect APG order because (i) *dpp* may contribute less than *wg* to axis formation (Jockusch *et al*., 2000; Niwa *et al*., 2000) and/or (ii) leg patterning may follow a “cascade model”, in which other transcription factors shape *dpp* domains to preserve PD patterning (Pechmann & Prpic, 2022). Similarly, changes in *dpp* deployment in three-dimensional *Cupiennius* legs may not substantially alter the effective overlap of Wg and Dpp gradients relative to *Drosophila* (Prpic *et al*., 2003).

Vn is an additional short-range morphogen with distinct leg versus wing roles in *Drosophila.* In legs, *vn* is activated at the distal centre by Wg and Dpp (Galindo *et al*., 2002) and is required for tarsal differentiation (Newcomb *et al*., 2018). High Vn promotes APGs including *al* and *Lim1* (Campbell, 2005), supports *rn* via *dac* together with Dll (St Pierre *et al*., 2002), and refines *B-H1* (Kojima *et al*., 2000) and *ap* domains (Pueyo *et al*., 2000). *vn* also contributes to medial fate regulation (Newcomb et al., 2018). In contrast, *vn* in *Drosophila* wings is expressed dorsally and promotes notum and dorsal compartment fate (Wang *et al*., 2000).

In *Bicyclus*, *vn* is deployed differently from *Drosophila* in both appendages: it is expressed at the distal tip in legs, but broadly across interveins in a pattern largely complementary to *dpp*, consistent with roles in vein development and later eyespot formation; *vn* is also expressed proximally in the hinge, similar to *Drosophila* wings. Differences in *vn* deployment may therefore contribute to leg-wing non-congruence of some APGs. Notably, non-congruent genes are largely narrow-domain APGs that respond to EGFR ligands such as Vn above concentration thresholds. Genes such as *B-H1*, activated by intermediate Vn concentrations in *Drosophila* legs (Campbell, 2005) and possibly also in *Bicyclus* leg (Fig. 8), may fail to activate in wings because *vn* is strong and broad.

### Evidence for a conserved Mecopteridan appendage-specific PD patterning network

Comparisons between *Drosophila* and *Bicyclus* indicate that most APGs are deployed in an appendage-specific order. In legs, most genes show spatial congruence (except *ss* and *apA*), consistent with broader comparative work on ventral appendage PD patterning. In the wing, *B-H1* is the only incongruent APG across species among the broad- and narrow-domain APGs. Although *al* is largely conserved in wings, *Drosophila* shows and additional notum/hinge *al* domain that remains poorly studied (Campbell & Tomlinson, 1998). Overall, these data support a conserved Mecopteridan intra-appendage PD patterning network.

Broader comparisons across arthropods also support partial conservation of ventral appendage-PD-patterning. Similar broad-domain APG deployment (*hth*/*exd*, *dac*, *Dll*) occurs in legs of chelicerates, myriapods, crustaceans, and hemimetabolous insects (Angelini & Kaufman, 2005; Bruce & Patel, 2020; Chen *et al*., 2016; Prpic & Telford, 2008). Broad-domain APGs (*hth*, *dac*, *dll*) also occur in similar order in the crustacean labrum (Bruce & Patel, 2025), and proximal-distal ordering has been reported outside arthropods (onychophorans, tardigrades, cephalopod tentacles) (Game & Smith, 2020; Janssen *et al*., 2010; Tarazona *et al*., 2019).

In contrast, narrow-domain APGs are less consistently characterised across panarthropods. *al* shows the strongest conservation of distal identity among the narrow-domain APGs across insects, chelicerates, onychophorans, and tardigrades (Fig. 8) (Beermann & Schröder, 2004; Klementz *et al*., 2026; Mapalo *et al*., 2024; Miyawaki *et al*., 2002; Oliveira *et al*., 2014). *ss*, *rn*, *B-H1*, and *Lim1* show greater variability across legs (Angelini *et al*., 2012; Mapalo *et al*., 2024; Oliveira *et al*., 2014; Setton *et al*., 2017; Toegel *et al*., 2009; Treffkorn & Mayer, 2019), and comparative data for *bab* and *apA* remain limited (Mapalo *et al*., 2024).

Overall, the strong spatial congruency of APGs between legs and wings supports a deeply conserved PD patterning scaffold shared between ventral and dorsal appendages. Recruitment and specialisation of wing selectors (*vg*, *scalloped*, *nubbin*, and *Zn finger homeodomain 2*) likely enabled the transition of a leg-like program to a wing program (Tripathi & Irvine, 2022). Functional studies are now needed to test whether these spatial parallels reflect shared upstream morphogenetic inputs or regulatory convergence.

## Conclusion

Our characterisation of seventeen APGs in *Bicyclus* embryonic legs and larval wings supports a deeply conserved PD patterning scaffold. Broad-domain APGs show high leg-wing congruence, indicating substantial reuse of proximal and distal positional coordinates, whereas divergence in the intermediate domain – especially the absence of broad *dac* wing expression, and differences in morphogen deployment (*dpp* and *vn*) – point to regulatory changes associated with wing-specific traits such as venation. These results expand PD patterning data in butterflies, highlight key differences from *Drosophila*, and motivate additional comparative and functional studies across Panarthropoda.

## MATERIALS AND METHODS

### Rearing *Bicyclus anynana* and embryo collection

*B. anynana* butterflies were reared in the lab at 27°C, 60% humidity and a 12-12 h day-night cycle. Larvae and adults were fed corn leaves and mashed bananas, respectively. Corn leaves were placed inside adult *Bicyclus* cages when they were most active to promote oviposition. All embryos were incubated at 28°C for 48 h prior to dissection.

### Hybridisation chain reaction (HCR) v3.0

The top reciprocal BLAST hit was used to identify the *Bicyclus* gene orthologs. For three of these genes, *ap*, *al*, and *bowl* (odd-skipped gene family), paralogs were retrieved with OrthoDB (v12.2) and OrthoFinder v3 (Table S1). The probe sequences for each gene are available in Appendix C, all DNA oligos were procured from Integrated DNA Technologies.

An overview of HCR v3.0 was adapted from Choi *et al*. (2018) and can be found in the supplementary folder (Fig. S4). All DNA primary probes were manually designed from the unique CDS region of the gene of interest and ordered from Integrated DNA Technologies. Ultra-thin glass needles were first prepared via a needle puller (Narishige, PC-10). The 48 h embryos’ outer membrane was then punctured with the needle before being fixed in 1×phosphate-buffered saline (PBS) with 0.1% Tween® 20 (PBST) supplemented with 4% formaldehyde for 30 minutes. Following fixation, the outer membrane was removed using a Dumont #5 fine forceps (Dumont, 11254-20), and embryos were transferred to 1×PBST. Dissection of 5^th^ instar larval wings followed a previously described protocol (Banerjee & Monteiro, 2020a) in 1×PBS under a Zeiss Stemi 305 microscope (ZEISS) before being transferred into 1×PBST with 4% formaldehyde for 30 minutes. After washing with 1×PBST, the HCR procedure for embryos and larval wings followed an adapted, formerly detailed protocol (Bruce *et al*., 2021; Choi *et al*., 2018), following all buffer recipes from the original protocol. After incubation with the detergent solution at 37°C for 30 mins, three 3-minute 1×PBST washes were performed before another two 3-minute washes with 5×saline-sodium citrate with 0.1% Tween® 20 (SSCT), all at room temperature. The preparation of 1 mL probe hybridisation buffer (PHB) was optimally supplemented with 0.02 µM of DNA stock solution for each target gene. From the PHB incubation step onwards until mounting, the sample remains at 4°C. After overnight incubation with PHB, samples were washed five times with probe wash buffer, followed by twice with 5×SSCT before a second incubation with amplification buffer supplemented with 6 nM of each fluorescent hairpin probe (Molecular Instruments Inc.) for 4 h. After four 20-minute 5×SSCT washes were performed, the samples were mounted in 60% glycerol (in 1×PBS). For embryos, an additional two glass slides were used as a spacer to flank the samples, preserving their three-dimensional integrity. Embryo and larval wings were imaged using the FLUOVIEW FV3000 Confocal Microscope (Olympus Corporation). All mRNA expression patterns shown are maximum intensity projections unless otherwise stated. At least three biological replicates were performed for each sample type.

## Supporting information

Lee et al. 2026_Supplementary Information

## Acknowledgements

We would like to thank Roy Quah for help with gathering the gene paralogs, Tan Lu Wee for help with the lab management, and Tong Yan from the DBS-CBIS confocal facility for access and training for the Olympus FV3000 confocal microscope. We would also like to thank Ralf Janssen, and the two anonymous reviewers, for taking time to help us improve the contents of this manuscript.

## Ethics approval

Not applicable.

## Consent for publication

Not applicable.

## Availability of data and materials

Not applicable.

## Authors’ contributions

J.L performed all experiments, curated and illustrated all data, prepared all figures, and wrote the initial manuscript draft (main text + supplementary). J.L, T.D.B, and A.M participated in conceptualisation. J.L and T.D.B contributed to the methodology. J.L, T.D.B, and A.M were responsible for manuscript refinement and subsequent revisions. A.M is responsible for supervision and funding acquisition. All authors have reviewed and approved the submission of this manuscript.

## SUPPLEMENTARY MATERIALS

### Appendix A: List of APG paralogs and gene families

**Table S1:**
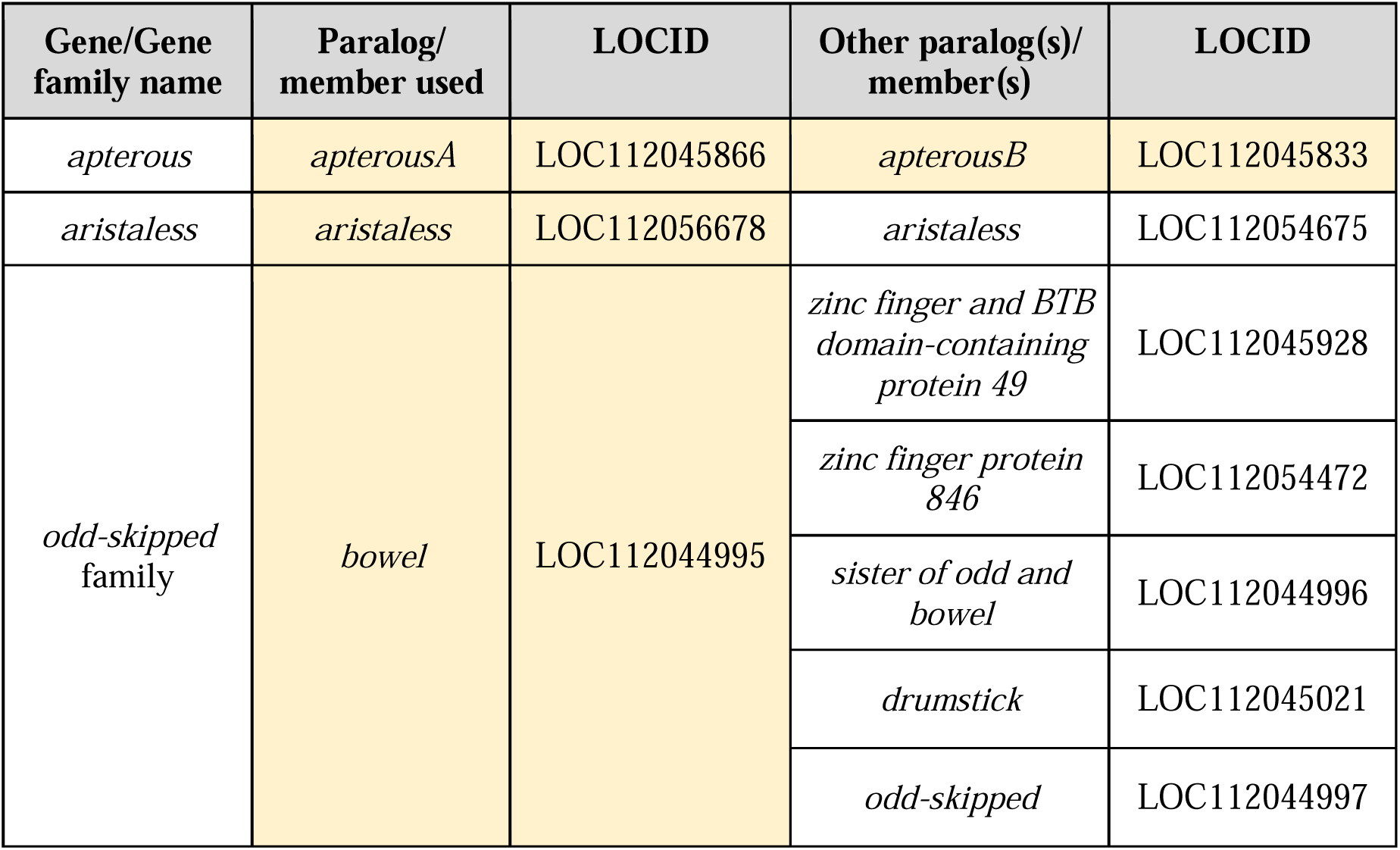
List of APGs in *Bicyclus anynana* with relevant paralogs or gene family members. All paralogs were identified through OrthoDB (v12.2) (https://www.orthodb.org/) and/or OrthoFinder v3 (https://github.com/OrthoFinder/OrthoFinder). The genes highlighted in yellow were tested in this manuscript.

### Appendix B: Supplementary expression of APGs in legs and wings

**Figure S2:**
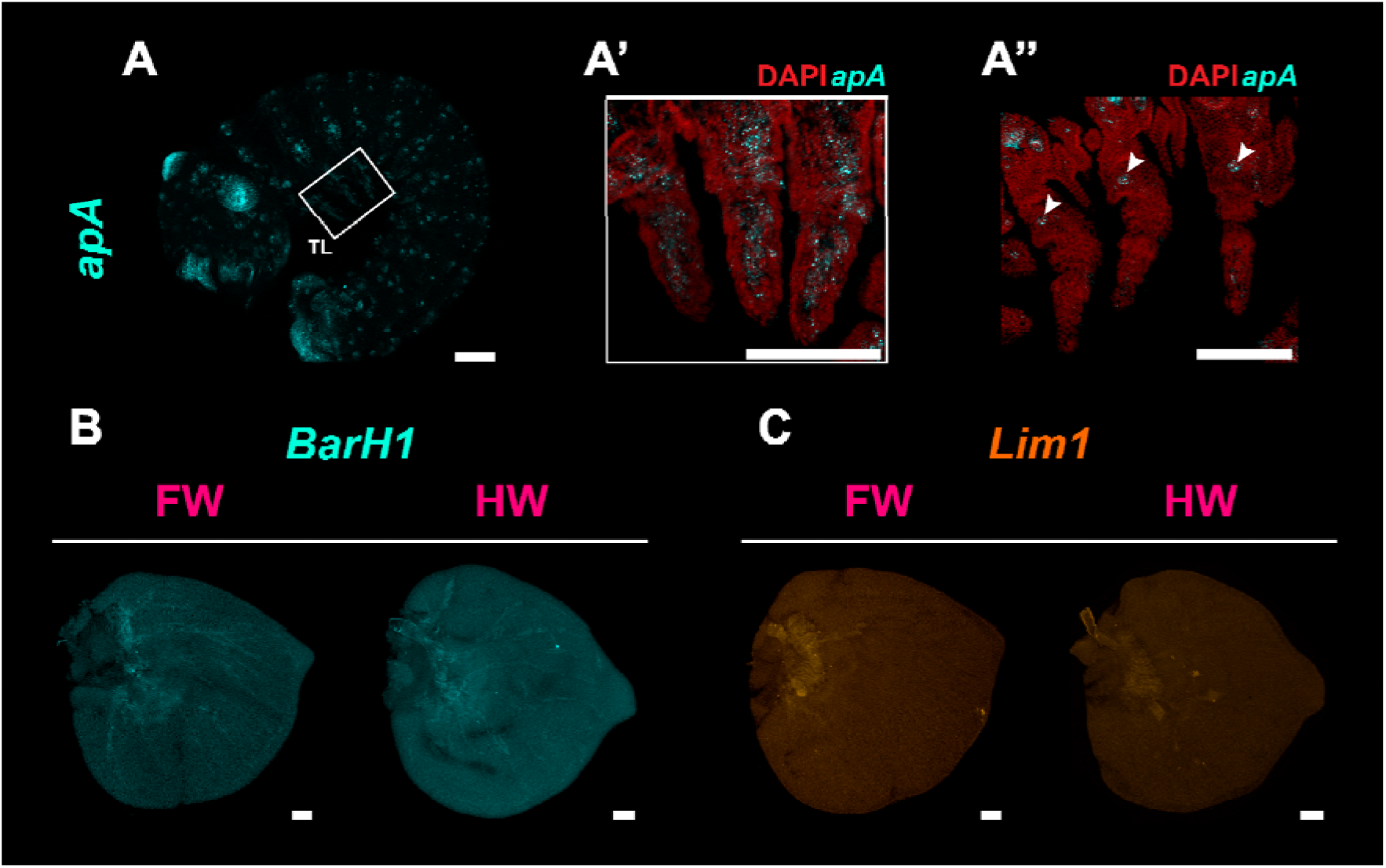
APGs with no expression in legs and wings. All wings are mid to late larval, and each subfigure consists of a forewing (FW; left) and hindwing (HW; right). **(A, A’)** *apA* is strongly expressed in mesodermal cell types throughout the centre of the leg. **(A’’)** Epidermal slice of an embryonic leg, where *apA* is not expressed. A small cluster of *apA-* expressing cells is observed at each thoracic segment, located in the ventral body wall and not at the proximal domain of the leg (white arrowheads). **(B)** *B-H1* and **(C)** *Lim1* are not expressed in the larval wings. Scale bars: 100 µm.

**Figure S3:**
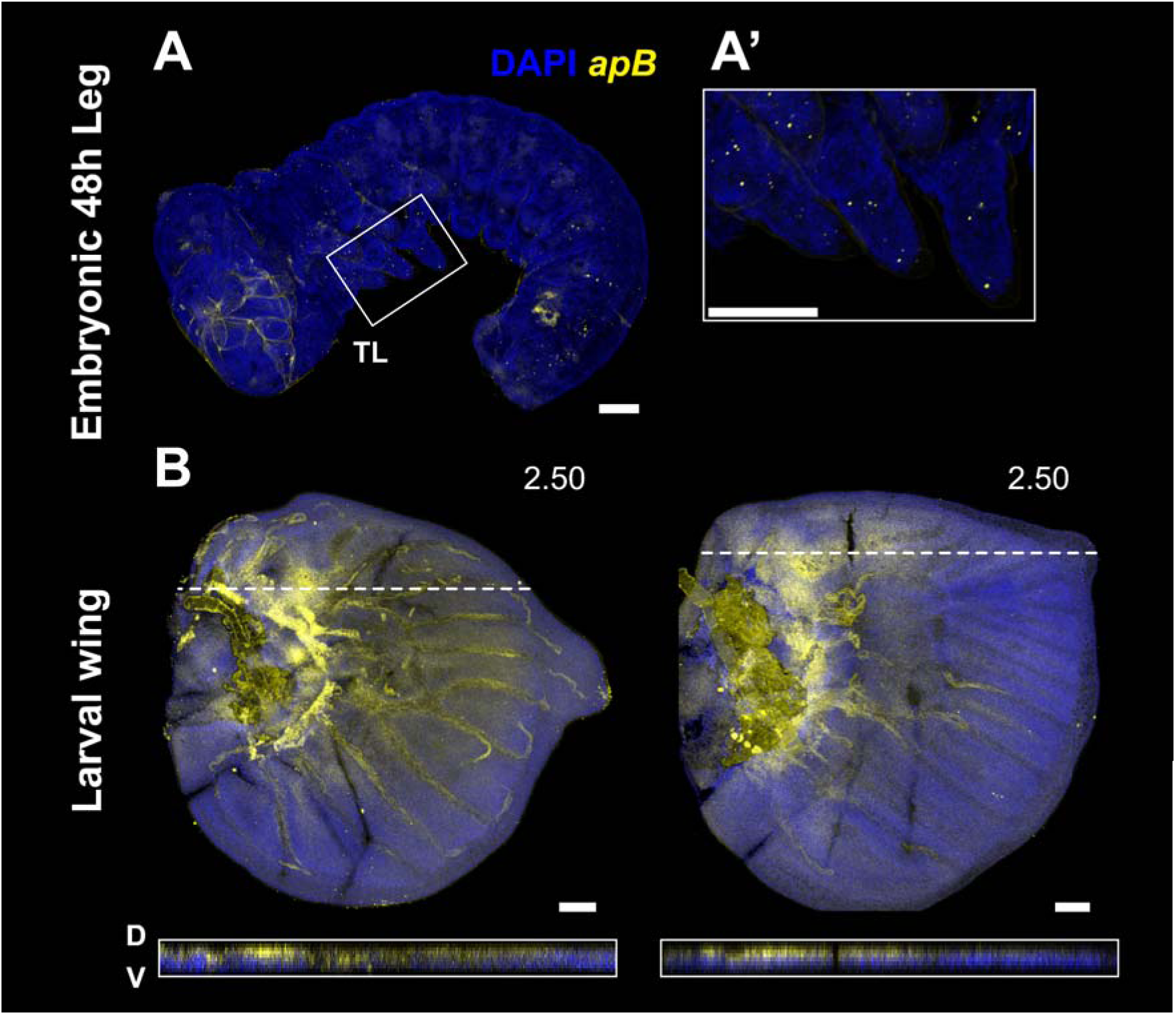
Expression of *apterous* paralog *apB* in legs and wings. **(A)** *apB* was not expressed in the TL, (A’) is a closeup of the legs from (A). **(B)** *apB* expression in the larval fore (left) and hind (right) wing. A DV cross-section of the white dashed line is displayed below each wing. While *apB* spatial expression domain is similar to that of *apA*, its expression is comparably more muted. Scale bar: 100 µm.

### Appendix C: HCR Probes

For each gene coding sequence, binding sites for split-initiator hairpin probes P1 and P2 are highlighted in green and yellow, respectively. Both P1 and P2, each carrying one-half of the HCR initiator I1 binding site, are required for subsequent binding of HCR amplifiers. Full P1 and P2 probe sequences are included below each respective sequence. Spacer bases are highlighted in blue.

### homothorax (hth)

>homothorax_LOC112054591_B1

https://www.ncbi.nlm.nih.gov/nuccore/XM_052888255.1

ATGGCTCAGCCTAGGTACGACGAGAGCCTCCACGGCGGGGGCTACATGGAGGGCGGCGCCATGTACCACGAGCACCGGCTCACGCACCCGCACATCCCGCCGGTGCACTACCCGCCGCCCGCCGCGCCGGCGCACGCGTTGCCCGGCGAGCCGCTAGTGCACAAGCGCGACAAGGACGCCATATACGGGCATCCCCTGTTTCCCCTGCTGGCGCTGATCTTCGAGAAGTGCGAGCTGGCAACGTGTACCCCCCGCGACCCCGGCGTAGCCGGCGGTGACGTCTGTTCCTCAGAGTCCTTTAACGAGGACATCGCGGTGTTCAGTAAACAGATACGTCAAGAAAAACCTTATTACATAGCGGACCCCGAGGTAGACTCATTAATGGTGCAAGCAATACAAGTCCTACGGTTTCACCTATTAGAATTAGAAAAAGTGCACGAGCTGTGCGACAACTTCTGCCACCGCTACATCAGCTGCCTGAAGGGCAAGATGCCCATCGACCTGGTGATCGACGAGCGGGAGTCAGCCCGGCCGCCGACACCAACGGGGAGCCGCGGTCGGCGCCTGACAGCAACCACGACGGCGCATCGACCCCCGACGTCAGGCCGCCATCGTCGTCGCTATCATACGGCGGTGCGGTGAACGATGAC GTCCGCTCACCGGGCTCCGGTGGCACCCCCGGTCCCCTCAGCCAGCCCCCGCCGCAGACCCTCGACGCGACAGATCCAGATGCCATGGGCAAATGGTGCGGGTCGCGGCGGGAATGGTCATCCCCTCCCGACGTGGCGCGGCGGGTCTACTCCTCAGTGTTCCTGGGCAGTCCCGGGGAATACCCAGGGGATGCCAGTAACGCGAGTATCGGCTCCGGCGAGGGTACGGGGGAGGAAGACGACGACACGAACGGAAAGAAGAACCAAAAG AAACGGGGAATCTTTCCGAAGGTCGCCACCAACATCCTTAGAGCGTGGCTCTTTCAGCACTTAACGCATCCCTACCCTTCGGAAGACCAGAAGAAACAGTTGGCACAAGACACAGGGTTAACGATACTACAAGTAAATAATTGGTTCATCAACGCGAGACGTAGGATAGTACAGCCAATGATAGACCAGTCGAATAGAGCAGTGTTCTACCCCGCAGTGTTCCCGCACGCGGGCCCCAGCGGCGCCTACAGCCCGGAGGCCACCATGGGCTACATGATGGACGGCCAGCAGATGATGCACAGGCCGCCGGCCGACCCCGCCTTCCACCAGGGCTACGCGCACTACCCCGCCGAGTACTACGGACACCATCTTTAA

Hth1_HCR_P1B1: gAggAgggCAgCAAACggAAGCGACCCGCACCATTTGCCCATGGC

Hth1_HCR_P2B1: CGTCGGGAGGGGATGACCATTCCCGTAgAAgAgTCTTCCTTTACg

Hth2_HCR_P1B1: gAggAgggCAgCAAACggAATTCCTCCCCCGTACCCTCGCCGGAG

Hth2_HCR_P2B1: GTTCTTCTTTCCGTTCGTGTCGTCGTAgAAgAgTCTTCCTTTACg

Hth3_HCR_P1B1: gAggAgggCAgCAAACggAACTGTTTCTTCTGGTCTTCCGAAGGG

Hth3_HCR_P2B1: TATCGTTAACCCTGTGTCTTGTGCCTAgAAgAgTCTTCCTTTACg

Hth4_HCR_P1B1: gAggAgggCAgCAAACggAAACTGCGGGGTAGAACACTGCACGGT

Hth4_HCR_P2B1: GCGCCGCTGGGGCCCGCGTGCGGGATAgAAgAgTCTTCCTTTACg

Hth5_HCR_P1B1: gAggAgggCAgCAAACggAAGTCCATCATGTAGCCCATGGTGGCC

Hth5_HCR_P2B1: CGGCGGCCTGTGCATCATCTGCTGGTAgAAgAgTCTTCCTTTACg

Hth6_HCR_P1B1: gAggAgggCAgCAAACggAATAGTGCGCGTAGCCCTGGTGGAAGG

Hth6_HCR_P2B1: AGATGGTGTCCGTAGTACTCGGCGGTAgAAgAgTCTTCCTTTACg

#### extradenticle

>extradenticle_LOC112056072_B2

Link: https://www.ncbi.nlm.nih.gov/nuccore/XM_052881231.1

ATGGACGATCCGAATAGAATGATGGCGCATAGCGGTGGTCTCATGGGGCCCCAAGGCTACGGGCTTCCTGGCGGGGACGGCGCGCCCGCTGCGGGCGATGGCGAGGCGAGAAAACAGGATATTGGCGAAATTCTGCAGCAAATCATGAACATTACCGACCAG

AGTCTCGACGAGGCGCAAGCTAGAAAACACACGCTGAACTGTCACAGGATGAAGCCTGCTCTTTTTTCCGTGTTGTGCGAAATCAAAGAAAAAACAGTGCTGTCTCTCCGCAACACGCAAGAGGAGGAGCCCCCAGACCCGCAGCTGATGAGGTTGGACAACATGCTGATAGCTGAGGGTGTGGCCGGCCCTGAGAAGGGTGGCGGCGCTGGTGCAGCTGCTTCAGCCTCCGCAGCAGCAGGGGAATGGGATAACGCGATCGAGCATTCCGACTACCGCGCGAAGCTGGCCCAGATCCGCCAGATCTACCACCAGGAGCTGGACAAGTATGAGAACGCCTGCAACGAGTTCACCACCCACGTCATGAACCTGCTGCGGGAGCAGAGCCGCACCAGACCCATCACTCCTAAGGAGATAGAGCGCATGGTGCAGATCATACACAAGAAGTTTAGTTCCATCCAGATGCAGTTGAAACAGTCCACGTGTGAGGCAGTTATGATCCTACGCTCACGTTTCCTGGACGCGCGCCGAAAGCGCCGCAACTTCAGCAAGCAGGCGTCAGAGATCCTGAACGAGTACTTCTACTCGCACCTCTC CAACCCTTACCCCAGCGAAGAGGCTAAGGAGGAGCTGGCGCGCAAGTGCGGCATCACTGTCTCCCAGGTGTCCAATTGGTTCGGCAACAAGCGCATCCGCTACAAGAAGAACATCGGCAAAGCGCAGGAGGAGGCCAACCTGTACGCCGCCAAGAAAGCTGCCGCCGGCGCTTCCCCCTACTCCATGGGCGCGGCGTCGGGCACGGCGACGCCCATGATGTCGCCGGCGCCCACGCAGGACTCCATGGGCTACTCGCTGCCGGCGCCGGGCTACGAGCAGTCGCAGCCGCCCTACGACGCGTCCATGGGCTACGACCCCATGCACCAGGACCTCTCGCCTTAG

Exd1_HCR_P1B3: gTCCCTgCCTCTATATCTTTGCACAACACGGAAAAAAGAGCAGGC

Exd1_HCR_P2B3: AGACAGCACTGTTTTTTCTTTGATTTTCCACTCAACTTTAACCCg

Exd2_HCR_P1B3: gTCCCTgCCTCTATATCTTTTCAGCTGCGGGTCTGGGGGCTCCTC

Exd2_HCR_P2B3: CAGCTATCAGCATGTTGTCCAACCTTTCCACTCAACTTTAACCCg

Exd3_HCR_P1B3: gTCCCTgCCTCTATATCTTTGCTGCACCAGCGCCGCCACCCTTCT

Exd3_HCR_P2B3: TCCCCTGCTGCTGCGGAGGCTGAAGTTCCACTCAACTTTAACCCg

Exd4_HCR_P1B3: gTCCCTgCCTCTATATCTTTCAGCTTCGCGCGGTAGTCGGAATGC

Exd4_HCR_P2B3: CTGGTGGTAGATCTGGCGGATCTGGTTCCACTCAACTTTAACCCg

Exd5_HCR_P1B3: gTCCCTgCCTCTATATCTTTGGGTGGTGAACTCGTTGCAGGCGTT

Exd5_HCR_P2B3: TCTGCTCCCGCAGCAGGTTCATGACTTCCACTCAACTTTAACCCg

Exd6_HCR_P1B3: gTCCCTgCCTCTATATCTTTACCATGCGCTCTATCTCCTTAGGAG

Exd6_HCR_P2B3: GAACTAAACTTCTTGTGTATGATCTTTCCACTCAACTTTAACCCg

Exd7_HCR_P1B3: gTCCCTgCCTCTATATCTTTCATAACTGCCTCACACGTGGACTGT

Exd7_HCR_P2B3: CGCGTCCAGGAAACGTGAGCGTAGGTTCCACTCAACTTTAACCCg

Exd8_HCR_P1B3: gTCCCTgCCTCTATATCTTTGGATCTCTGACGCCTGCTTGCTGAA

Exd8_HCR_P2B3: AGAGGTGCGAGTAGAAGTACTCGTTTTCCACTCAACTTTAACCCg

#### tiptop (tio)

>tiptop_LOC112051061_B1

Link: https://www.ncbi.nlm.nih.gov/nuccore/XM_024089545.2

ATGATCGTCGAGCCAGACCACGTGAGGGAGAGTCCAAGATGTTTGTCACGGGAG TCGTCCGGCGCGCCACGATGTCCCTCCAACGACTCCGTGTACTCGGGTCGGAGCG CGCCGAGTCTGCCTCTGCCGGCTGCGTTGCAAGCGGCGCTCCCGGCAGGTCTCCC CGCGGCGCTCCTGCCCCCCCACTCCGCTGCCGTGGCGGCCTACCTCGGCGCAGCC GCCGCCGCCGCCCAGCAGAGGCTCCTCATGTCCTGCCAAGAAGATATGACAGAC TCAGAACGCGCTGACGCAGTACTAGACTTCAGCACGAAGCGAAGTGAATCCCCC ATCGATGATGAAGACGACGATGCCGTCAATCTGAGCAAAAACGAAAACGGCCCC TTAGATTTATCAGTAGGCACCAGAAAACGAGGACCACAAAATTCTCCATCACCCGTTCCTACGAGAAAAAGTTCACGGACGGCTGATTTTAAACCGATCACTACTCCGT GGACTACGCCGGTTGCGCCACATCTTCCTTATTTTGCAGCAGCTGTTGCAGCTGC TAGCTTATCACCTAAAGGTGGTGTCCCACCAGACTGGAATGGAAAATTAAAACA TGGGACACCGACACCTAACGATGCAACTAAAGCGCTTGAAAAAATGAGCGAGTT AAGTAGGTTAGGTGGAGAGGAGTTATTTAGGTCAGTTCAAAGCGCAGCTTTAGG AGCCGGGCTCACTCCTAACGCAGCCGCTCGCCACTCTGCCTGGCAGTCGCATTGG TTGAATAAAGGCGCCGACCAAACGAAAGACGTTTTGAAATGTGTGTGGTGCAAG AAAAGTTTTAACTCTTTAGCTGATTTAACTGTTCATATGAAAGAAGCTAAGCATT GCGGCGTGAACGTCCCGGTACCACCATCGGGCGGTGCACCGATCCCTCCTTCACT TCTGCCGCCTTCTAGTTCACCTTCGACGCCATCTTACAACTCTTCTTCGTCGAGCG GATCGTCTAAACCTAATCACAACGATCTTAATATGCTAATTAAGGAAAACATGCC CATACCTAGAAAACTAGTCCGCGGCCAAGACGTATGGTTAGGTAAAGGGGCCGAACAAACTAGGCAAATATTAAAATGTATGTGGTGCGCCGAAAGTTTCCGCTCTTTAGCCGAAATGACGAGCCACATGCAGCGCACCCAGCACTACACTAATATTATATCACAAGAACAAATCATTTCTTGGAAGTCTTCAGATGAAGCCAAGGGCTCCAACTCTAGCGCACCCGGTACTAATAACGCGGTTCCTCCGACGACAGGGACGAGTAGTCACGTTAGCGCTGTTTTAACTTGCAAAGTATGCGACCAAGCGTTTAGTTCCTTAAAAGAATTAAGTAATCATATGGTTAAGAACTCTCACTATAAAGAGCACATTATGCGCTCGATAACTGAAAGCGGTGGCAGACGACGACAAACGCGCGAAAAGAGAAAGAAGTCCTTGCCTGTGAGGAAACTTCTCGAGCTCGAGAGAGCCCAACATGAGTTTAAAAATGGGGAGGGCAACGGAGTCCCTATGGGAAAACCTATTAGGGATTTTGGCTCTGGAAGCCGTATCACTTGCGAGAAATGCGGCGACAAAATTGAGACTGCTGTATTCGTAGAACACATCCGACAATGCATAGGCGCGCCCATGTCGAATAGCCAAAGGAATTTTCTAAAAAGTGCTCTACTTTCCAATAATATTATTCCACCCGACGTGCCCGGCCACATCACTCCCACCAGTCGCGACGGACGTAAGAGTATCAATGACGAAATTCCCTCACCAGGGTCGACGCATCACCGTTCACCTTCTTCGGTTAACGATTCTTCTCCTAGTTC TAAAGATCACAACGTCAGCAACGATAAAAGCTCATCACCATCCGTACTCAATGC CATAGAACAATTAATAGAGAAGAGTTTCGATACTCGCTCGCGACATTCGGTCCCC GGTATGCCAGGCGGAGCTTCCCACGCCCCTATTGGTTCTAGTATTTTAAAAAGAT TAGGTATAGACGAGAGCGTAGACTACACGAAACCTCTGGTAGATCCACAAACGA TGAGCATGTTGAGAAATTATCACCACCAGCAAGGCTACGGTCGTCGAGAGCGCA GCGGTAGCGAATCTAGTTCCATGTCCGAGAGGGGCGGCAGTAGGGTGGAGTCTC TCACGCCGGACAGAAAGTTGGATTCGTATCATATGACCCCGCGTACGACTCCCGA CACTCGGGGATCCCAGACCCCCGCTTCGGAGGATCGCCCCGCAGAGGTCAGGAT AAAAAAAGAAATCACCGACGAAGAAGATAGGGAGACTTGCGTAGACTTGAGCA GTCAGCCGGTCAGGGTTAAAACCGAAGTGGACGATGACGAGGAAGCGAGGCCG AACAGTGTGGCGGACGAAGACGTGAAGCCGACCGTCCCCAAGCGCGAGAGCGA GGGTCCGAGCCCCGTGCCTAGCCCTCGCAGCCCGGCCAGCGACCGCTCGGCGCC CACTCCCGGCGCTGACAGAAAGCCGGCTTCCAGCCTAGGCGCGCTCTCCTCCATG TTCGACAACCTCGCTGGGGGAGCCTCCTCCAACGAGCCCAGCTCGTCTCGTCGCG GGGGTAGTCACCCCCTGGCGGCCTTGCAAAAACTATGCGACAAAACAGACACTA ATTCTTCCCGCGCACCGCCCCCCGCGCCGTCCCCCGCCGGGCCGCCGAGTATTTT AACTTTCAGTTGGGCTTGCAACGACGCGGTAGTCACAGATTCGATAATGAAATGC GCTCTTTGCGACACACCGTTTATTTCGAAAGGCGCGTACCGGCACCACCTGTCCA AGATGCACTTCGTGAAGGACGGCGCACTGCCAGAGCCGGTGCCGGTGAAGGCGC CGCCGCCCGCGCCGTCGCCGGGCGCGCACAAGTCCGGCTCCACCGCCGCGTCGC CCCAGGACCCGCGGAGTCCGTCCCAGTCGTTCGACGAGAGCCCGCACTCCAAGT TCCTCAAGTATACGGAGCTGGCGAAGCAGCTGTCCAGCAAGTACGTGTAG

Tio1_HCR_P1B1: gAggAgggCAgCAAACggAACGGCTAAAGAGCGGAAACTTTCGGC

Tio1_HCR_P2B1: GGGTGCGCTGCATGTGGCTCGTCATTAgAAgAgTCTTCCTTTACg

Tio2_HCR_P1B1: gAggAgggCAgCAAACggAACAAGAAATGATTTGTTCTTGTGATA

Tio2_HCR_P2B1: GAGCCCTTGGCTTCATCTGAAGACTTAgAAgAgTCTTCCTTTACg

Tio3_HCR_P1B1: gAggAgggCAgCAAACggAACGTCGGAGGAACCGCGTTATTAGTA

Tio3_HCR_P2B1: AGCGCTAACGTGACTACTCGTCCCTTAgAAgAgTCTTCCTTTACg

Tio4_HCR_P1B1: gAggAgggCAgCAAACggAATTAAGGAACTAAACGCTTGGTCGCA

Tio4_HCR_P2B1: TCTTAACCATATGATTACTTAATTCTAgAAgAgTCTTCCTTTACg

Tio5_HCR_P1B1: gAggAgggCAgCAAACggAACTTTCAGTTATCGAGCGCATAATGT

Tio5_HCR_P2B1: TCGCGCGTTTGTCGTCGTCTGCCACTAgAAgAgTCTTCCTTTACg

Tio6_HCR_P1B1: gAggAgggCAgCAAACggAAGAGCTCGAGAAGTTTCCTCACAGGC

Tio6_HCR_P2B1: ATTTTTAAACTCATGTTGGGCTCTCTAgAAgAgTCTTCCTTTACg

Tio7_HCR_P1B1: gAggAgggCAgCAAACggAAAATCCCTAATAGGTTTTCCCATAGG

Tio7_HCR_P2B1: CGCAAGTGATACGGCTTCCAGAGCCTAgAAgAgTCTTCCTTTACg

Tio8_HCR_P1B1: gAggAgggCAgCAAACggAATGTTCTACGAATACAGCAGTCTCAA

Tio8_HCR_P2B1: ATGGGCGCGCCTATGCATTGTCGGATAgAAgAgTCTTCCTTTACg

Tio9_HCR_P1B1: gAggAgggCAgCAAACggAAGGAAAGTAGAGCACTTTTTAGAAAA

Tio9_HCR_P2B1: GGGCACGTCGGGTGGAATAATATTATAgAAgAgTCTTCCTTTACg

Tio10_HCR_P1B1: gAggAgggCAgCAAACggAATGATACTCTTACGTCCGTCGCGACT

Tio10_HCR_P2B1: TCGACCCTGGTGAGGGAATTTCGTCTAgAAgAgTCTTCCTTTACg

#### dachshund (dac)

>dachshund_LOC112044740_B3

Link: https://www.ncbi.nlm.nih.gov/nuccore/XM_052883449.1

ATGGAGTCCGCCGTGGACTCAGCGTCGACCGCCAGCGAGGTGAGCGGCTCGTCCGGGGGCTCGCCGCGCGTCAAGGCGGCGTCGCCGGCGCGCGGCCTCAGCCCGCCGCAGCTGCTGGCGCCGCGCCTGCCGCTGCCGCCGCCGGGCCTGGGCCTGCTGGGCTCGCTGCAGATGATGCACCACTCGCCGCTCGAGCTCATGGCGGCGGCGCACCACCACCCGCACCGCTACGGGAGCCCGCCGCCCATCTCCACGTCCGACCCCTCGGCGA ACGAGTGCAAGCTGGTGGACTATCGCGGGCAGAAGGTGGCCGCGTTCATCATCCAGGGCGACACGATGCTGTGCCTGCCGCAGGCCTTCGAGCTGTTCCTGAAGCACCTGGTGGGCGGGCTGCACACCGTGTACACCAAGCTGAAGCGGCTGGACATCGTGCCGCTGGTGTGCAACGTGGAGCAGGTCCGCATCCTGCGCGGGCTGGGCGCCATCCAGCCGGGCGTCAACCGCTGCAAGCTGCTCTCGTGCAAGGACTTCGACGTGCTGTACCGCGACTGCACCACGGCAAGGTGCCTGTCAATGAAAGCGCCAGACAGCTCCAGACCGGGTCGACCTCCGAAGCGTGCTTCTGGAGTTGGTCTTTCGCTTGCCGCCACGC AGTTCCCGGGGCACCCCTTCAAGAAGCACCGCCTAGAGAATGGGGACTACTCGC CGTATGAAAATGGACATATGAGCGAGATGGCCCGCATGGATAAGTCCCCGCTCC TCGCGAACGGGTACAACGCACCCCCCACCCACCTGGGGCCCATGGGCTTCATGC ACCAGCACGCCCTCATGTCTCCGGGTATGCCACACCCTGGCGTGCCCAGGCCTGA CGGGTCCATCATTAAGGGGCAGCCTATGCATAACATGGAAGCACTAGCAAGATC TGGTATTTGGGAGAATTGTAGAGCAGCATACGAGGACATCGTAAAACATCTCGA AAGGTTGCGTGATGAACGGGGTGATATTGAACGTGTTATAGCAATGGACAAGGC GCGCGAGGGTTCACATAATGGTTCGTCTCCAGGCCACAGTCCTGTGCTGAACCTT TCGAAGTCAGGCTCCGGTGAGCGCGAGCGGTCCGAGCGGGACCGCGAGCGCGCG GAGCGGGGCGAAGGCTCGGCGAGCGGGCGCAGCTCGGCCGCCTCGCGCCGCACG CCGCAGCCGCCGCGCATCCCCTCCACCGCCGCGCCGGTCTCCCCGCGCTCCCACT CCGACGAGAGCGACGCTGCTCTGTCTGATCAAGACGACCATAACGTCAAAGACG AGGATGACGATCTCAGCGATGGTGAACGAGACCTCGCAACGAATTCCTCGCCAG CGCCCGTCAGCTACCCCCCACAAGGCTCGCCGTCGAACGTTCCCGTGGACCCCAC CGCCGACACCCTGGTCTCCTCCACCGAGACCCTCCTCAGGAACATCCAGGGCCTC CTCAAAGTAGCGGCGGACAACGCACGCCAACAAGAACGGCAAATCAGTTACGAAAAAGCGGAACTAAAAATGGACGTGTTAAGAGAAAGAGAAGTAAAAGACAACC TGGAAAGACAGTTACTGGACGAACAAAAAATGAGAGTAATGTATCAGAAGAGAT TAAAGAAAGAAAGAAAGCAGAGGCAACAAATTCAAGACCAATTAGAATTGGAG TTAAAAAGACGACAGAAGATAGAAGAAGCACTAAAGCAGTCAGGAGCGCCCGG TGAAATTCTGAGAATAGTAACTGAGAATTTAACACCACCGTCCCAAGAAAATCG CGAACGTGAGAACGGTACAGAGAGCAAACCGCCAAGTACAGAGCCCCCCACCTC GTCGCCGCCGTTCCAGAGGGACCCCCCGCGCACGCCCGACAAGCCCCAGTGGAA CTACCCTCCGCCGCCTGTCGACATCATGAGCGGAGGAGCTGCTTTCTGGCAGAAC TACTCTGAATCCTTGGCGCAAGAGTTGGAGATGGAGCGCAAGTCCCGCCAGCAG GCCATGGAGAGAGACGTGAAGAGCCCTCTGTCGGACCGCGCCAGTTACTACAAG AACTCGGTGCTGTTCAGCTCGGCCACTTAG

Dac1_HCR_P1B3: gTCCCTgCCTCTATATCTttCCTCGCTGGCGGTCGACGCTGAGTC

Dac1_HCR_P2B3: GCGGCGAGCCCCCGGACGAGCCGCTTTCCACTCAACTTTAACCCg

Dac2_HCR_P1B3: gTCCCTgCCTCTATATCTTTGCAGGCCCAGGCCCGGCGGCGGCAG

Dac2_HCR_P2B3: AGTGGTGCATCATCTGCAGCGAGCCTTCCACTCAACTTTAACCCg

Dac3_HCR_P1B3: gTCCCTgCCTCTATATCTTTTGCGGGTGGTGGTGCGCCGCCGCCA

Dac3_HCR_P2B3: GAGATGGGCGGCGGGCTCCCGTAGCTTCCACTCAACTTTAACCCg

Dac4_HCR_P1B3: gTCCCTgCCTCTATATCTTTGGCCTGCGGCAGGCACAGCATCGTG

Dac4_HCR_P2B3: CACCAGGTGCTTCAGGAACAGCTCGTTCCACTCAACTTTAACCCg

Dac5_HCR_P1B3: gTCCCTgCCTCTATATCTTTCTGGATGGCGCCCAGCCCGCGCAGG

Dac5_HCR_P2B3: GAGCAGCTTGCAGCGGTTGACGCCCTTCCACTCAACTTTAACCCg

Dac6_HCR_P1B3: gTCCCTgCCTCTATATCTTTAGCACGCTTCGGAGGTCGACCCGGT

Dac6_HCR_P2B3: GGCGGCAAGCGAAAGACCAACTCCATTCCACTCAACTTTAACCCg

#### Distal-less (Dll)

>Distal-less_LOC112055223_B2

Link: https://www.ncbi.nlm.nih.gov/nuccore/XM_052887174.1

ATGCATCGCGCCACGAGAGGCCGGGTAGCGTTCGGAGCAGTCCAGAAATCCCTCAAAATTACACGAATTCAGTCCCCAAACAGCAAACCCACCACGCTGAGTTTCTCAGATCCCTTTGGGCCTCCTCAGTCCACGGACGGGGGGGGCCCGTCGACCCCGCAGCCCGCCATGACCACCCAGGAGCTGGACCACCAACACCACCACCTGGGAGGTTCA CAAACCCCCCACGATATATCCAACTCCACGAATTCAACCCCCACCAACGTTTCCTCGAAATCCGCCTTCATAGAGTTACAGCAACACGGTTACGGGCCTTTCAAAGGTGGTTACCAACATCCCCACCATTTCGGCAGCCCGGGGGGTCAGCAGAACCCCCACGAGGCGTCGGGGTTCCCGAGCCCTAGGTCCTTAGGTTATCCCTTTCCTCCTATGCACCAAAATACGTACGGATATCACATAGGCTCCTACGCTCCTCAATGTGCAAGTCCGCCCAAAGATGAAAAATGTGGTCTATCCGACGACCCTGGGCTGCGGGTGAACGGCAAGGGCAAGAAGATGCGCAAACCGCGCACCATCTACTCCAGCTTGCAGCTGCAG CAGCTCAACAGGCGGTTCCAGAGAACACAGTATCTAGCACTGCCGGAGCGAGCT GAGCTGGCTGCTAGCTTGGGCCTGACGCAGACGCAGAACAGTAGCGCTATTTCG TAADll1_HCR_P1B2: CCTCgTAAATCCTCATCAAAGAACGCTACCCGGCCTCTCGTGGCG

Dll1_HCR_P2B2: AATTTTGAGGGATTTCTGGACTGCTAAATCATCCAgTAAACCgCC

Dll2_HCR_P1B2: CCTCgTAAATCCTCATCAAACGGGCTGCGGGGTCGACGGGCCCCC

Dll2_HCR_P2B2: GGTGGTCCAGCTCCTGGGTGGTCATAAATCATCCAgTAAACCgCC

Dll3_HCR_P1B2: CCTCgTAAATCCTCATCAAATTGCTGTAACTCTATGAAGGCGGAT

Dll3_HCR_P2B2: ACCACCTTTGAAAGGCCCGTAACCGAAATCATCCAgTAAACCgCC

Dll4_HCR_P1B2: CCTCgTAAATCCTCATCAAAGCTGACCCCCCGGGCTGCCGAAATG

Dll4_HCR_P2B2: GGAACCCCGACGCCTCGTGGGGGTTAAATCATCCAgTAAACCgCC

Dll5_HCR_P1B2: CCTCgTAAATCCTCATCAAATTCATCTTTGGGCGGACTTGCACAT

Dll5_HCR_P2B2: CCCAGGGTCGTCGGATAGACCACATAAATCATCCAgTAAACCgCC

Dll6_HCR_P1B2: CCTCgTAAATCCTCATCAAAGGTTTGCGCATCTTCTTGCCCTTGC

Dll6_HCR_P2B2: AGCTGCAAGCTGGAGTAGATGGTGCAAATCATCCAgTAAACCgCC

#### spineless (ss)

>spineless_LOC112043391_B2

Link: https://www.ncbi.nlm.nih.gov/nuccore/XM_024078792.2/

ATGCACAAAGAGAAGGAGGAGCTGCACCGGCGTGACCACGCGCACTACGACCA CATCATGCCGGACGGCGAGATGTTCCTACAGGCACTCAACGGGTTCCTGATGATC CTCACGTGCGAGGGGGAGGTGTTCTTCGCGACACACAGCATCGAGAGCTACCTG GGCTTCCATCAGTCAGACATAGTCCACCAATCAGTGTACGAGTTGGTCCACTCAG AAGATAGAGAAGAGCTTCAGAGGCAGTTACTTTGGAACTCCTTCCTGCCACCAG AAGCAAACAGTCTCTCACTGCAAGATGTTTTGAGACAGGAGAGAGCGCATCTAT TAGAAAGAAGTTTTACAGTCAGATTTAGATGTTTATTAGACAATACATCAGGATT TTTGAGATTAGACATAAGAGGCCGGATCAAAGTATTACATGGACAGAACAAGAA AACAGAAGAGCCACCATTAGCTTTATTTGCAATTTGCGCACCGTTCGGGCCTCCCAGCCTGCTGGAGATACCACAGAAAGAGGTCATGTTCAAGAGCAAACACAAACTGGATTTATCACTAGTCTCGATGGATCAAAGAGGGAAACTTCTCCTCGGTTACACGGACGCCGAGCTCGCAAACATGGGAGGTTACGATTTAGTGCATTACGACGACCTGGCGTATGTAGCCAGCGCTCATCAGGAATTACTGAAGACAGGAGCGTCAGGGATGATCGCATACCGCTTCCAGACGAAGGACGGCCAGTGGCAGTGGCTGCAGACTTCCTCGCGACTCGTCTACAAGAACTCCAAGCCTGACTTCGTGATTAGCACGCATAGGCCTCTAATGGAAGAAGAAGGCAGAGATCTCCTCGGGAAGCGGACGATGGACTTCAAAGTTAGCTATTTAGACACCGGTCTCACCAGCCTTTACTTTTCCGAAACAGAACAGCTTTCAGGCTGCGAGTCGAACGTCTCAACGCCCCCTCGAGCAGCGAGGCGATATAAGACTCAACTGAGAGACTTTCTATCCACTTGTAGGAATAAAAGACGAGTGCCTCCCACTCCCGCGCCACCTCCAGTGGATTACTTAGCTGCTGATGCGGTAGCCGCAGCGTATACTAATCTAAACGGAGCGTACACCGCTCCATACGGGGAATATCCTCCCGCTCCAGCGTTACATTACGCTCCACCACCTTTAGATGATAGATTCCTCGCAGCTGAC AACCTCTTCCATCAATACAAACCATTGTCTTACCATTACACTCCATACGCGCCTA ATGGTTTCTTAGAACCGGCTCCACCTCCTGGATACGAAGTTCCACCGACGTATCA TCGACCTCCGTCAAGAGAATACACTTATGTTGATAGTAGTGGGAGGTACATGAGT CCGGTGACTGGAGAGAGACGGTCGCCAAATGTCGTGCCTGGTCCAAGTCCGGGT GGGTCAAGCGGTTCGACAGAGCAGGATAGGATGGTGCAAACGCAAACGTCGGA GATGCCAAGGCAGACGGTCCTCATGTGGGGTGCCGGTGGAGCAGCTCAAGAAGT CCCCGTGGAGTACTCACCACCGCAGGTTTGGCGATACCATCCCTCCCACTACCAC ACCGCTGAAGCGACTCAATGA

Ss1_HCR_P1B2: CCTCgTAAATCCTCATCAAATGACCTCTTTCTGTGGTATCTCCAG

Ss1_HCR_P2B2: AATCCAGTTTGTGTTTGCTCTTGAAAAATCATCCAgTAAACCgCC

Ss2_HCR_P1B2: CCTCgTAAATCCTCATCAAACCGAGGAGAAGTTTCCCTCTTTGAT

Ss2_HCR_P2B2: ATGTTTGCGAGCTCGGCGTCCGTGTAAATCATCCAgTAAACCgCC

Ss3_HCR_P1B2: CCTCgTAAATCCTCATCAAATACATACGCCAGGTCGTCGTAATGC

Ss3_HCR_P2B2: CTTCAGTAATTCCTGATGAGCGCTGAAATCATCCAgTAAACCgCC

Ss4_HCR_P1B2: CCTCgTAAATCCTCATCAAACCTTCGTCTGGAAGCGGTATGCGAT

Ss4_HCR_P2B2: AAGTCTGCAGCCACTGCCACTGGCCAAATCATCCAgTAAACCgCC

Ss5_HCR_P1B2: CCTCgTAAATCCTCATCAAAATCACGAAGTCAGGCTTGGAGTTCT

Ss5_HCR_P2B2: TCTTCCATTAGAGGCCTATGCGTGCAAATCATCCAgTAAACCgCC

Ss6_HCR_P1B2: CCTCgTAAATCCTCATCAAATTTGAAGTCCATCGTCCGCTTCCCG

Ss6_HCR_P2B2: GGTGAGACCGGTGTCTAAATAGCTAAAATCATCCAgTAAACCgCC

Ss7_HCR_P1B2: CCTCgTAAATCCTCATCAAAACTCGCAGCCTGAAAGCTGTTCTGT

Ss7_HCR_P2B2: CTGCTCGAGGGGGCGTTGAGACGTTAAATCATCCAgTAAACCgCC

Ss8_HCR_P1B2: CCTCgTAAATCCTCATCAAACAAGTGGATAGAAAGTCTCTCAGTT

Ss8_HCR_P2B2: GTGGGAGGCACTCGTCTTTTATTCCAAATCATCCAgTAAACCgCC

Ss9_HCR_P1B2: CCTCgTAAATCCTCATCAAATACCGCATCAGCAGCTAAGTAATCC

Ss9_HCR_P2B2: TCCGTTTAGATTAGTATACGCTGCGAAATCATCCAgTAAACCgCC

Ss10_HCR_P1B2: CCTCgTAAATCCTCATCAAAACGCTGGAGCGGGAGGATATTCCCC

Ss10_HCR_P2B2: CATCTAAAGGTGGTGGAGCGTAATGAAATCATCCAgTAAACCgCC

#### rotund (rn)

>rotund_LOC112056645_B3

Link: https://www.ncbi.nlm.nih.gov/nuccore/XM_024096578.2

ATGATGGAAGGAAGGACGATCGATTACAGGCCAGACGGTGGTGGGCTCGATTAC CATAACTCTCCGTTAATGGCGGAGATACCTGTAGACAATTACTCCCATATCCATA GAAGTATAGAGCATTTGAGGTCGATGGGAGTGGCACCACACCTGGATCCCCATA GACATTTAGCTGCGAATCTGACAGACTTGCGAAGGTACGAACACCCTGATGACA TGCCTGAGATAAAGCCGTCCGTACTTAGATTATCCGAATTCAAAAGTGGTCTACAAGAAATAAGGGTGCACAATGACCAAGAGAATGACATTCAGCGTCAAATGACACCGCACGACGATGGAAAAGGTTACAGCGCACCTTCAACGCCTTTATCAGAAAACGGCGGACAAAGCATTCAAGAGGAAAAGGTGTTCGGCTCCAAAGCGGACCTCCAGCTGCACACGCAGATCCACCTGCGCGAAGCGAAGCCCTACCGCTGCACCCAGTGCCCGAAGGCCTTCGCCAACTCCTCATACCTAGCGCAGCATTCGCGCATCCACCTCGGCATCAAACCGTACCGCTGCGAGATCTGCCAGCGCAAGTTCACCCAACTGTCTCACCTGCAGCAGCACATCCGCACCCACACGGGTGACAAACCCTATCGATGCACCCAGATAGGGTGCACCAAAGCGTTCTCCCAGCTGTCCAACTTGCAGAGCCACAGCCGCTGCCACCAGACCGACAAACCCTTCAAATGCAACTCCTGCTACAAATGCTTCACGCAC GAGAAAGACCTCCTCGAACACATTCCGAAACACAAAGAATCCAAGCATTTGAAG ACCCACATCTGCCAGTACTGCGGCAAGAGTTACACCCAGGAGACGTACTTAAGT AAACACATGAACAAACACGCTGAAAGAGCTGACAAGAGGCCGCCGATCTCCGCG TTAGGACTGAGCGGGCTCAACAGGTCCTTAGGAGCAGCCGCGCCCACTGCGGCC CCTTTCGCGGATCACCCATATTGGCCTAAAGTTAGTCCGGATTCCGCCGCGCACA TGTCTGATGAAGGCGGGTATCATCAGCAGAGGGAGAGCGACGATCACCATGAGC ATCAACTGCAGCAGCAAAGGGCGCTGTTCGCTCAACACGAGAGTCAGGAAGACC GCATCCAGCCGCCGGTGTCGTCGGCTGCGAATTCCGCTTTCACGCCGATAAACTC CATGGCGCCTCATCTCAACGGTCTCTCGCATCACAGCGCTCTGCCCACGCGGCCG TATCTCTACGACCCGTTGCACTTCCAGCAAGGGAAGCAGCAGCCTAACTCCTTCC CCAATCAGCTGATATCGCTCCACCAAATCCGGAACTACGCGCACCAGCCGTCCGC CCTGTTGCCTGCCGAGCACATCCTCCCCCATTCCCTCTCTCACAAAGACAAGCAGTAA

Rn1_HCR_P1B3: gTCCCTgCCTCTATATCTTTTACGGACGGCTTTATCTCAGGCATG

Rn1_HCR_P2B3: ACCACTTTTGAATTCGGATAATCTATTCCACTCAACTTTAACCCg

Rn2_HCR_P1B3: gTCCCTgCCTCTATATCTTTGAATGTCATTCTCTTGGTCATTGTG

Rn2_HCR_P2B3: CATCGTCGTGCGGTGTCATTTGACGTTCCACTCAACTTTAACCCg

Rn3_HCR_P1B3: gTCCCTgCCTCTATATCTTTCCGTTTTCTGATAAAGGCGTTGAAG

Rn3_HCR_P2B3: ACCTTTTCCTCTTGAATGCTTTGTCTTCCACTCAACTTTAACCCg

Rn4_HCR_P1B3: gTCCCTgCCTCTATATCTTTGTGGATCTGCGTGTGCAGCTGGAGG

Rn4_HCR_P2B3: GCAGCGGTAGGGCTTCGCTTCGCGCTTCCACTCAACTTTAACCCg

Rn5_HCR_P1B3: gTCCCTgCCTCTATATCTTTGCGCTAGGTATGAGGAGTTGGCGAA

Rn5_HCR_P2B3: TGATGCCGAGGTGGATGCGCGAATGTTCCACTCAACTTTAACCCg

Rn6_HCR_P1B3:gTCCCTgCCTCTATATCTTTAGTTGGGTGAACTTGCGCTGGCAGA

Rn6_HCR_P2B3: GTGCGGATGTGCTGCTGCAGGTGAGTTCCACTCAACTTTAACCCg

Rn7_HCR_P1B3: gTCCCTgCCTCTATATCTTTGCACCCTATCTGGGTGCATCGATAG

Rn7_HCR_P2B3: GTTGGACAGCTGGGAGAACGCTTTGTTCCACTCAACTTTAACCCg

Rn8_HCR_P1B3: gTCCCTgCCTCTATATCTTTTGAAGGGTTTGTCGGTCTGGTGGCA

Rn8_HCR_P2B3: TGAAGCATTTGTAGCAGGAGTTGCATTCCACTCAACTTTAACCCg

#### bric-a-brac (bab)

**>**bricabrac_LOC112043806_B2

Link: https://www.ncbi.nlm.nih.gov/nuccore/XM_052890735.1

ATGCCGGCCGAGGAAGACCCCGGGGCGATGGACGGCCGCGAAGCCGCCGACGG CTCGCCGCAGCAGTTCTGCCTGCGATGGAACAACTACCAGGCCAACCTGGCCAC CTGCTTCGACCAGCTGCTGCAGACCGAGTCGTTCGTGGACGTGACGCTGGCGTGCGAGGGCCGCAGCCTGAAGGCGCACAAGGTGGTGCTGTCCGCCTGCAGCCCGTACTTCCAGACGCTGTTCACGGAGAACCCGTGCCGGCACCCGATCGTCATCATGCGCGACATCAAGTACTGCGACCTGAAGGCGGTGGTGGACTTCATGTACCGCGGCGAGATCAACGTGTCGCAGGACCAGATCGCGGCGCTGCTGCAGGTGGCGGAAACGCTGAAGGTGCGCGGGCTGACGGACGTGAGCGGCGAGCACGCCGTGCTGCGCGCCGCCAAGCGCAGCGGCACCGCGTCGCCGGACGCGCCGGGCGCGGGGCTGGCGGGCCCGGCCGGCAAGGCGCGCCGCCGCTCGGGCTCCGCGCCGCGCTCGCCGCCCGCCGACCTGCTGCACGCGCCGCCCGACGCGCCGCCGCCCATGCACGACGACGTCGACATCCGCCCCGGCATCGCCGAGATGATCCGCGAGGAGGAACGGGCCAAATTAATTGAAAACTCTCACGCTTGGCTCGGCGCCTCCACATCTTCGATTGCAGATAGCTACCAATATCAGCTGCAGTCCATGTGGCAGAAGTGCTGGAACACCAACCAGAGCATCGTGCACAATCTGAGGTTCCGCGAGCGCGGGCCGCTGAAGTCCTGGCGCCCGGAGACCA TGGCTGAGGCCATATTCTCTGTTCTGAAGGAAGGGCTCTCACTGTCGCAGGCTGCGAGGAAATACGACATTCCGTACCCGACATTTGTGCTGTACGCTAACAGAGTCCAC AACATGCTGGGCCCGAGCGCGGACGGTGGTGCAGATCTGAGACCCAAAGGTCGC GGGCGGCCGCAGCGCATTCTGCTGGGCGTGTGGCCCGACGAGCACATCCGCGGC GTCATCCGCGCCGTCGTGTTCCGCGACTCGCACGCCGCGCACGCCGTGCACGCCT CGCACCACATCAAGGAGGAGGTGCTCCCCGTCTACCCTGCGATCACGGACGGCA TGACCTCGGTGCAGAACGCCTACCCGGTGGCGTGCGGCAACGGGCGCGAGGGCG GCGTGTCGCCCAACGCCGCGGCAGCCGCCGCCGTGGCCGCCGTGGCGCACGGGC TGCGCCGCCAGATGTGCAACATGGTCGTGGCCGCGCAGAGGCCGCCCGACCACC CGGTGCACCTCGGCCTGCCGCCGATGCCCATGCAGTCACCGCGGTTGCCATCGCC GCGCCTGCCGTCGCCGCTCACGGCCGGCTCGGAGTCGCCGCTGTGCTCGCCGCCG CCGCCGGGCCACGCCCTCGAGCCGCCGAAGCCCACCGACCCGTTCCGACCCTTCG CCTCGCCGCGCCCTGACAGCCGCTACAGCGACCGCGACGTCGTCGCCGACATGA CCGACATGGGACCGCCTAAAACTCCGATTTCGATGGTGAAGAACGGCAAGGAGA TGTCGCTGCCGGTGAAGATGGAGCTGCCGCCGGCGGAGGCGCGCGCGGAGAACATCTGA

Bab1_HCR_P1B2: CCTCgTAAATCCTCATCAAAGTTCTCCGTGAACAGCGTCTGGAAG

Bab1_HCR_P2B2: CATGATGACGATCGGGTGCCGGCACAAATCATCCAgTAAACCgCC

Bab2_HCR_P1B2: CCTCgTAAATCCTCATCAAATGAAGTCCACCACCGCCTTCAGGTC

Bab2_HCR_P2B2: GCGACACGTTGATCTCGCCGCGGTAAAATCATCCAgTAAACCgCC

Bab3_HCR_P1B2: CCTCgTAAATCCTCATCAAATTCAGCGTTTCCGCCACCTGCAGCA

Bab3_HCR_P2B2: CCGCTCACGTCCGTCAGCCCGCGCAAAATCATCCAgTAAACCgCC

Bab4_HCR_P1B2: CCTCgTAAATCCTCATCAAACGCGGTGCCGCTGCGCTTGGCGGCG

Bab4_HCR_P2B2: CAGCCCCGCGCCCGGCGCGTCCGGCAAATCATCCAgTAAACCgCC

Bab5_HCR_P1B2: CCTCgTAAATCCTCATCAAAGCGCGGAGCCCGAGCGGCGGCGCGC

Bab5_HCR_P2B2: GCAGCAGGTCGGCGGGCGGCGAGCGAAATCATCCAgTAAACCgCC

Bab6_HCR_P1B2: CCTCgTAAATCCTCATCAAATCGACGTCGTCGTGCATGGGCGGCG

Bab6_HCR_P2B2: ATCATCTCGGCGATGCCGGGGCGGAAAATCATCCAgTAAACCgCC

Bab7_HCR_P1B2: CCTCgTAAATCCTCATCAAAAGCGTGAGAGTTTTCAATTAATTTG

Bab7_HCR_P2B2: AATCGAAGATGTGGAGGCGCCGAGCAAATCATCCAgTAAACCgCC

Bab8_HCR_P1B2: CCTCgTAAATCCTCATCAAAACTTCTGCCACATGGACTGCAGCTG

Bab8_HCR_P2B2: GCACGATGCTCTGGTTGGTGTTCCAAAATCATCCAgTAAACCgCC

#### BarH1 (B-H1)

>BarH1_LOC112051348_B1

Link: https://www.ncbi.nlm.nih.gov/nuccore/XM_024089951.2

ATGACGGTGCAGAGGGACGAGCGGGACCCGCGCGCGCCCCGGACCAGGTTCATG ATCACGGACATCCTCGACGCCGCGCCGCGGGACCTCAGCGCGCATCGGGACTCCGACTCCGACAGATCCGCCACCGACTCGCCCGGTGTTAAAGACGACTCAGACGATGTATCAAGCAAGTCGTGTGGCGGCGACGCCTCAGCTCTGTCGAAGAAGCAGAGGAAAGCGAGGACAGCCTTCACGGACCACCAGCTCCAGACCCTGGAGAAGTCGTTCGAGAGACAGAAGTACCTCAGTGTACAGGACCGCATGGAGCTGGCCGCCAAACTGGGCCTTACGGACACACAAGTCAAGACCTGGTACCAAAATAGACGGACGAAATGGAAGCGTCAAACCGCCGTCGGGCTGGAGCTGCTAGCCGAGGCGGGCAACTACGCAGCGCTGCAGCGGCTGTACGGCGGCTACTGGGCCGGGGTGCCGGCCTACCCCGCGCAGCCCGCGCCCGCCGCCGACCTCTACTACCGGCAGGCCGCCGCCAACGCCGCTGCCGCCGCCTCTGCCTCGGCTAACACGCTGCAGAAACCTTTGCCCTATCGGTTATATCCAGGTGCGCCTCTCGCAGGCGTCCCACCTTTGGGGCTCGGCCTCCCGGGGCCGTCAGCTCACCTCGGCTCTCTGGGGGCTCCAGCTCTCGGGGCCTTGGGATACTAC GCACAGGCGAGACGAACCCCCTCGCCGGACCTAGACCCGGGGAGCCCCGCGCCCCCCCACCGGTCCCCAAGGGAGCCCTCCTTAGAACGGCGCACTGACGACGAAGAC GACGACGAACCCATACATGTGTAG

B-H11_HCR_P1B1: gAggAgggCAgCAAACggAACTTGATACATCGTCTGAGTCGTCTT

B-H11_HCR_P1B2: GCTGAGGCGTCGCCGCCACACGACTTAgAAgAGTCTTCCTTTACg

B-H12_HCR_P1B1: gAggAgggCAgCAAACggAAGTCCGTGAAGGCTGTCCTCGCTTTC

B-H12_HCR_P1B2: CGACTTCTCCAGGGTCTGGAGCTGGTAgAAgAGTCTTCCTTTACg

B-H13_HCR_P1B1: gAggAgggCAgCAAACggAAGCTCCATGCGGTCCTGTACACTGAG

B-H13_HCR_P1B2: TGTCCGTAAGGCCCAGTTTGGCGGCTAgAAgAGTCTTCCTTTACg

B-H14_HCR_P1B1: gAggAgggCAgCAAACggAATTCCATTTCGTCCGTCTATTTTGGT

B-H14_HCR_P1B2: AGCTCCAGCCCGACGGCGGTTTGACTAgAAgAGTCTTCCTTTACg

B-H15_HCR_P1B1:gAggAgggCAgCAAACggAAGTACAGCCGCTGCAGCGCTGCGTAG

B-H15_HCR_P1B2: GGCCGGCACCCCGGCCCAGTAGCCGTAgAAgAGTCTTCCTTTACg

B-H16_HCR_P1B1:gAggAgggCAgCAAACggAAGCCGGTAGTAGAGGTCGGCGGCGGG

B-H16_HCR_P1B2: CGGCGGCAGCGGCGTTGGCGGCGGCTAgAAgAGTCTTCCTTTACg

B-H17_HCR_P1B1: gAggAgggCAgCAAACggAACGATAGGGCAAAGGTTTCTGCAGCG

B-H17_HCR_P1B2: CCTGCGAGAGGCGCACCTGGATATATAgAAgAGTCTTCCTTTACg

B-H18_HCR_P1B1: gAggAgggCAgCAAACggAAGTGAGCTGACGGCCCCGGGAGGCCG

B-H18_HCR_P1B2: GAGAGCTGGAGCCCCCAGAGAGCCGTAgAAgAGTCTTCCTTTACg

#### aristaless (al)

>aristaless_LOC112056678_B3

Link: https://www.ncbi.nlm.nih.gov/nuccore/XM_024097138.2

ATGGGAGTATCAGAACCAAACTGCTCCTCAACTCCCGACCTTCCCCCTCACGACCCGGAGCGACCCGGTTCAGGCAGCGGCATGGATGACGAGGACATTCCCAGAAGGA

AGCAGAGGCGGTATAGGACCACCTTCACTAGCTACCAGCTCGACGAACTGGAGAAGGCTTTTGGGAGGACGCATTATCCTGATGTATTTACCAGGGAGGAGTTAGCTTTAAAAATTGGCCTCACAGAAGCAAGAATACAGGTGTGGTTCCAAAACCGGCGGGCGAAATGGCGTAAGCAGGAGAAGGTGGGGCCCCATGCACATCCGTATAGCGGATACTTAAGCAGCGGGCAACCATTACCGACTACATCTATGCCAGTTCCACCGCATTCGTTCAGTCAGCTCGGATTTGGATTGAGAAAGCCTTTTGACAACGCTCTAGCGTCTTTCAGGTACGCCAACAGTCCCCTGTTTGGAGCACAGTACCTTCCGCCGCTTTCTCGTCCTCCACTTTTCGGGGCTCCACTATACGCTACATCGCCCGCTCATTTCCACTCTCTGTTCGCCAATCTAACCGTACCCGAACTACCTCGAATCTCACCTGAACAATCACGATTGTCACCCGAAGTAACTCGATCGCCAGCACCGTCGATCTCCCCTCCAATATCACCAGGCAGCGAGACTTTACCCCCATCTGAAGATGTTAGGAGTTCCAGCATAGCAGCATTAAGACTAGCTGCTAGAGAACACGAACTCAGATTAGAATTGTTGCGACAG CGAGCGGATTTAATTTGTCAATAG

Al1_HCR_P1B3: gTCCCTgCCTCTATATCTTTGAATGTCCTCGTCATCCATGCCGCT

Al1_HCR_P2B3: TCCTATACCGCCTCTGCTTCCTTCTTTCCACTCAACTTTAACCCg

Al2_HCR_P1B3: gTCCCTgCCTCTATATCTTTAAAGCCTTCTCCAGTTCGTCGAGCT

Al2_HCR_P2B3: AATACATCAGGATAATGCGTCCTCCTTCCACTCAACTTTAACCCg

Al3_HCR_P1B3: gTCCCTgCCTCTATATCTTTTGCTTCTGTGAGGCCAATTTTTAAA

Al3_HCR_P2B3: CCGGTTTTGGAACCACACCTGTATTTTCCACTCAACTTTAACCCg

Al4_HCR_P1B3: gTCCCTgCCTCTATATCTTTGTGCATGGGGCCCCACCTTCTCCTG

Al4_HCR_P2B3: CGCTGCTTAAGTATCCGCTATACGGTTCCACTCAACTTTAACCCg

Al5_HCR_P1B3: gTCCCTgCCTCTATATCTTTGAATGCGGTGGAACTGGCATAGATG

Al5_HCR_P2B3: CTCAATCCAAATCCGAGCTGACTGATTCCACTCAACTTTAACCCg

Al6_HCR_P1B3: gTCCCTgCCTCTATATCTTTGTTGGCGTACCTGAAAGACGCTAGA

Al6_HCR_P2B3: AAGGTACTGTGCTCCAAACAGGGGATTCCACTCAACTTTAACCCg

Al7_HCR_P1B3: gTCCCTgCCTCTATATCTTTCGTATAGTGGAGCCCCGAAAAGTGG

Al7_HCR_P2B3: GAGAGTGGAAATGAGCGGGCGATGTTTCCACTCAACTTTAACCCg

Al8_HCR_P1B3: gTCCCTgCCTCTATATCTTTGGTGAGATTCGAGGTAGTTCGGGTA

Al8_HCR_P2B3: ACTTCGGGTGACAATCGTGATTGTTTTCCACTCAACTTTAACCCg

Al9_HCR_P1B3: gTCCCTgCCTCTATATCTTTTGGTGATATTGGAGGGGAGATCGAC

Al9_HCR_P2B3: TTCAGATGGGGGTAAAGTCTCGCTGTTCCACTCAACTTTAACCCg

Al10_HCR_P1B3: gTCCCTgCCTCTATATCTTTTAGCAGCTAGTCTTAATGCTGCTAT

Al10_HCR_P2B3: ACAATTCTAATCTGAGTTCGTGTTCTTCCACTCAACTTTAACCCg

#### Lim homeodomain 1 (Lim1)

>Lim1_LOC112054918_B2

Link: https://www.ncbi.nlm.nih.gov/nuccore/XM_052883120.1

ATGCCGACGTACGGAGGACGCGGCGAGGCGGGCGGCGCCATGGCGTGCGCCGG CTGCGAGAAGCCCATCCTGGACAAGTTCCTGCTGCACGTGCTGGAGCGCGCGTG GCACGCGGCGTGCGTGCGCTGCGCCGACTGCCGCGCGCCGCTCGCCGACAAGTGCTACTCCAGGGATAACAAACTCTTTTGTAGGAATGATTTCTTTAGACGTTACGGTACAAAATGCAGCGGCTGCGGTCACGGAATATCCCCATCAGATCTAGTTAGGAAAGCACGGGAAAAGGTGTTCCACCTGAATTGCTTCACCTGCTTAGTCTGCAGGAAGCAACTGTCCACTGGTGAAGAATTGTACGTGCTAGACGACAACAAATTTATATGTAAAGAGGATTACTTAGCGGGGAAGGCGCCGACGCATTCTGACTCACTACTCGGCTCAGGTTCAGATGACGAAGAAGAGGAAGAAGCGAGAGCAGCAAACAATTCAAGTAGTCCGGCGCATGCACCGCACCCTGCGCTACATTCTGAGCTATCACATAATGGGGACACCAAACCGCACGAAGATTCGGAAGATCAGGGTTCGCTGGACGGGGACACGGAGACCAGAGACTCTCAAGCGGAGAACAAGTCACCAGATGACGGCAACGGTGGCTCCAAAAGAAGAGGGCCTCGGACGACGATAAAGGCGAAACAGTTGGAGATACTGAAATCGGCTTTTTCACAGACACCCAAACCGACGAGGCACATACGAGAACAGCTGGCAAAAGAAACAGGGCTACCCATGAGGGTGATACAAGTCTGGTTTCAAAACAAGC GGTCAAAGGAACGGCGTCTAAAACAGCTGACGTCGATGGGCAGGGGGCCTTTCT TCGGCTCGTCGCGCAAGATGCGCGGCTTCCCCATGAACCTGTCCCCGGGGGGCCT GGAGGAGGGGCCGCCGGGGTTCCCATACTTCGCGACAGCAGATGGAAAGTTCGA ATTTGGCTATGGCCCGCCGTTTCACCATGACGCGCCGTTCTTCCATCCGCCGCCC GCTATGCCGTTCAACCAGCCAGGTGGTATGGAGACGCTGCAAGGCGGCGAGTTC CAGGACCAGTTCCCTCATCACGAGCACCTGGTGCTGCCGCGGCCTTCTTCCCCCG AGTTCGCGTTCGGCGACGCCCCCCCTCCTCTCCACCCCGAGGGTCTGGTGTGGTAG

Lim11_HCR_P1B2: CCTCgTAAATCCTCATCAAAGTGACCGCAGCCGCTGCATTTTGTA

Lim11_HCR_P2B2: CCTAACTAGATCTGATGGGGATATTAAATCATCCAgTAAACCgCC

Lim12_HCR_P1B2: CCTCgTAAATCCTCATCAAAAGCAGGTGAAGCAATTCAGGTGGAA

Lim12_HCR_P2B2: CAGTGGACAGTTGCTTCCTGCAGACAAATCATCCAgTAAACCgCC

Lim13_HCR_P1B2: CCTCgTAAATCCTCATCAAATTACATATAAATTTGTTGTCGTCTA

Lim13_HCR_P2B2: GGCGCCTTCCCCGCTAAGTAATCCTAAATCATCCAgTAAACCgCC

Lim14_HCR_P1B2: CCTCgTAAATCCTCATCAAATTCGTCATCTGAACCTGAGCCGAGT

Lim14_HCR_P2B2: GTTTGCTGCTCTCGCTTCTTCCTCTAAATCATCCAgTAAACCgCC

Lim15_HCR_P1B2: CCTCgTAAATCCTCATCAAAAATGTAGCGCAGGGTGCGGTGCATG

Lim15_HCR_P2B2: TGGTGTCCCCATTATGTGATAGCTCAAATCATCCAgTAAACCgCC

Lim16_HCR_P1B2: CCTCgTAAATCCTCATCAAATCCCCGTCCAGCGAACCCTGATCTT

Lim16_HCR_P2B2: TCCGCTTGAGAGTCTCTGGTCTCCGAAATCATCCAgTAAACCgCC

Lim17_HCR_P1B2: CCTCgTAAATCCTCATCAAATCTTCTTTTGGAGCCACCGTTGCCG

Lim17_HCR_P2B2: TTTCGCCTTTATCGTCGTCCGAGGCAAATCATCCAgTAAACCgCC

Lim18_HCR_P1B2: CCTCgTAAATCCTCATCAAAGTTTGGGTGTCTGTGAAAAAGCCGA

Lim18_HCR_P2B2: CCAGCTGTTCTCGTATGTGCCTCGTAAATCATCCAgTAAACCgCC

#### apterousA (apA)

>apterousA_LOC112045866_B2

Link: https://www.ncbi.nlm.nih.gov/nuccore/XM_024082231.2

ATGGGAGTGTACGAAGAGAGGGGGGCGATGCACTGGCAACAGAGTGAGCGGTACTTATCATCGGCATACGAAACTGGGGCCGAGTTGTCACCAGTGGCGCCTGTAGCT TCACCAGGGTCCCCCCGTGATTGCACGTCGTGTCGGAAGAGGGAGCCCCCAGACGAGCCCCCACCGCCGCCCACCGAAGACGCTTGCGCTGGTTGCGGGGCGCGGATCACGGACAGATACTACCTACTCGCCTTAGAACGACGCTGGCACACACCCTGCTTGAGGTGCTGCGAATGCAAAATGCCCCTAGACTCCGAACAGAGGTGTTACGCTCGGAACAGTAACATATTCTGCAAGAACGACTACTTCAGGTTGTATGGATCAAAGCGATGCGCGAGATGTAACACCCCTATATCATCGTCGGAGCTGGTGATGAGGGCGCGTGACCTAGTGTTTCATGTGCACTGTTTCTCCTGTGCCCTGTGTAGTACGCGACTCACGAAGGGTGACGAGTACGGTATTAGAAACTCAGCTGTGTATTGCAGGCTACATTTCGAAACAATGCCGGATTATGGTCCTCATATGGCGGTACCAGGTCCTCCGCAAATGTGCCCGGGGCCTTACGTAGGACCACCACCGGGGCCGCACTACCCCGCCTACCCTTCGCCGGAGTTCGGTCGAGTAGAACCTGATGTTCCTAAAGGCCCTTTTTTCAACGGGACTTCAGCGCCACCACCCCGACAAAAAGGTCGTCCGAGAAAAAAGAAACCGAAG GACCAAGATATCATGTCGGCCAACCTTGATCTAAATGCTGAATACTTGGAGATGGGATTCCGCGGCGGTGGCGGCTTGGGCAGCTCGTCGCGCACAAAGCGCATGCGAACCAGCTTCAAGCATCACCAGCTCCGCACCATGAAGTCGTATTTCGCGATCAACCACAATCCTGACGCGAAGGACCTGAAACAGTTGAGCCAGAAGACAGGCCTCCCGAAAAGAGTGTTACAGGTATGGTTCCAGAACGCGCGAGCGAAATGGCGGCGCATGGTGACGAAGCAGGAAAACAAAATGGCGGACAAGTTGTCACCAGACGGCTCGCTGG AGATGGACATGTACCATGGACCGCTTGGGTCCATTCAGTCGCTTCCTCCGCACAGTCCGCCTTACAGCGTAATGGGGGGTCCACCGAGTCCAAGTTCGATGGATTGCCCCTAG

ApA1_HCR_P1B2: CCTCgTAAATCCTCATCAAAGGCGAGTAGGTAGTATCTGTCCGTG

ApA1_HCR_P2B2: GCAGGGTGTGTGCCAGCGTCGTTCTAAATCATCCAgTAAACCgCC

ApA2_HCR_P1B2: CCTCgTAAATCCTCATCAAATCTGTTCGGAGTCTAGGGGCATTTT

ApA2_HCR_P2B2: ATATGTTACTGTTCCGAGCGTAACAAAATCATCCAgTAAACCgCC

ApA3_HCR_P1B2: CCTCgTAAATCCTCATCAAACATCGCTTTGATCCATACAACCTGA

ApA3_HCR_P2B2: GATGATATAGGGGTGTTACATCTCGAAATCATCCAgTAAACCgCC

ApA4_HCR_P1B2: CCTCgTAAATCCTCATCAAACACATGAAACACTAGGTCACGCGCC

ApA4_HCR_P2B2: ACTACACAGGGCACAGGAGAAACAGAAATCATCCAgTAAACCgCC

ApA5_HCR_P1B2: CCTCgTAAATCCTCATCAAACTGAGTTTCTAATACCGTACTCGTC

ApA5_HCR_P2B2: TTTCGAAATGTAGCCTGCAATACACAAATCATCCAgTAAACCgCC

ApA6_HCR_P1B2: CCTCgTAAATCCTCATCAAAGGAGGACCTGGTACCGCCATATGAG

ApA6_HCR_P2B2: CCTACGTAAGGCCCCGGGCACATTTAAATCATCCAgTAAACCgCC

ApA7_HCR_P1B2: CCTCgTAAATCCTCATCAAACTCCGGCGAAGGGTAGGCGGGGTAG

ApA7_HCR_P2B2: AGGAACATCAGGTTCTACTCGACCGAAATCATCCAgTAAACCgCC

ApA8_HCR_P1B2: CCTCgTAAATCCTCATCAAAGTCGGGGTGGTGGCGCTGAAGTCCC

ApA8_HCR_P2B2: GTTTCTTTTTTCTCGGACGACCTTTAAATCATCCAgTAAACCgCC

#### apterousB (apB)

>apterousB_LOC112045833_B3

Link: https://www.ncbi.nlm.nih.gov/nuccore/XM_024082182.2

ATGCTAAAGGAACGAGAAAGCTCGGAGGGGTCCCCTTCAACCCCAGACGAATGT GCTGGATGCGGAGGCAGGATACAGGACAGATATTACCTGCTTGCGGTAGACCGACAATGGCACGGCTCCTGCCTCCGCTGCTGCGAATGCCGTCTGCCTTTGGACAGCGAGTTGACCTGCTTCTCAAGAGACGGAAATATCTATTGCAAGGAAGACTACTATAGGTTGTTTTGTGTGAAAAGGTGTGCTCGATGTGGGAACGGTATCACGGCGAACGAGCTGGTGATGCGAGCCCGCGACATGGTCTTCCATCTCACATGCTTCACCTGCGTTGCATGCGGAACCCTACTTTCGAAAGGCGATGTCTTTGGGATGAAAAACGGCCTAATTTACTGCAGACCTCACTATGACAGTGCGTGTATGGATGATTTTTGTGACGAGGACATCAATAGTGTTTACAGGTGCCAAGAAATGCATAGCGAGGGGGATTCACCAAACCAATACTACAATGGAGGCCCGAGCCAAAAGGGTAGACCAAGAAAGAGAAAAGTGGCCCAAGGTTCCCCAGAAGATATGCAAGTTCAAACGATGAGGATGGCGAGCACAGCTTTAGAAATACTTCACCGCGGGGATCTCTCTTCATCAATGGAGTCGCTGGCTTACGACTCGTCAGTCGCTTCACCGGGTAGTGTTTCGAGCCATAATCAACGCACTAAACGCATGCGCACCAGTTTTAAGCATCACCAGCTTCGCACCATGAAGTCATACTTTGCAATCAATCAGAACCCTGACGCAAAGGATCTTAAACAACTGGCTCAAAAGACTGGTCTTTCAAAAAGGGTATTGCAGGTATGGTTTCAAAACGCAAGAGCAAAATGGCGTCGGAATATGATGAGGCAAGAGTCCAGCCAGCATGGATCTATGATGAACGGCAACCTGGGACATCCAGCAAACGGGGCGTTGGTCGGCGGCGTACCTCCACC GAGTGTGCCAAGCACCATGATCTTATCAGAGTCTCTCCAAGCTTTAGACGATCTG AGGGTCCACACCCCCCACCCGATGACCTATAATGAAATGTACTGA

ApB1_HCR_P1B3: gTCCCTgCCTCTATATCTTTGTCAACTCGCTGTCCAAAGGCAGAC

ApB1_HCR_P2B3: TAGATATTTCCGTCTCTTGAGAAGCTTCCACTCAACTTTAACCCg

ApB2_HCR_P1B3: gTCCCTgCCTCTATATCTTTACACCTTTTCACACAAAACAACCTA

ApB2_HCR_P2B3: CGCCGTGATACCGTTCCCACATCGATTCCACTCAACTTTAACCCg

ApB3_HCR_P1B3: gTCCCTgCCTCTATATCTTTTGAGATGGAAGACCATGTCGCGGGC

ApB3_HCR_P2B3: TTCCGCATGCAACGCAGGTGAAGCATTCCACTCAACTTTAACCCg

ApB4_HCR_P1B3: gTCCCTgCCTCTATATCTTTAGGCCGTTTTTCATCCCAAAGACAT

ApB4_HCR_P2B3: CTGTCATAGTGAGGTCTGCAGTAAATTCCACTCAACTTTAACCCg

ApB5_HCR_P1B3: gTCCCTgCCTCTATATCTTTAACACTATTGATGTCCTCGTCACAA

ApB5_HCR_P2B3: CTCGCTATGCATTTCTTGGCACCTGTTCCACTCAACTTTAACCCg

ApB6_HCR_P1B3: gTCCCTgCCTCTATATCTTTTTTGGCTCGGGCCTCCATTGTAGTA

ApB6_HCR_P2B3: CCACTTTTCTCTTTCTTGGTCTACCTTCCACTCAACTTTAACCCg

ApB7_HCR_P1B3: gTCCCTgCCTCTATATCTTTATCCTCATCGTTTGAACTTGCATAT

ApB7_HCR_P2B3: TGAAGTATTTCTAAAGCTGTGCTCGTTCCACTCAACTTTAACCCg

ApB8_HCR_P1B3: gTCCCTgCCTCTATATCTTTGTCGTAAGCCAGCGACTCCATTGAT

ApB8_HCR_P2B3: AACACTACCCGGTGAAGCGACTGACTTCCACTCAACTTTAACCCg

ApB9_HCR_P1B3: gTCCCTgCCTCTATATCTTTTAAAACTGGTGCGCATGCGTTTAGT

ApB9_HCR_P2B3: ACTTCATGGTGCGAAGCTGGTGATGTTCCACTCAACTTTAACCCg

ApB10_HCR_P1B3: gTCCCTgCCTCTATATCTTTTTAAGATCCTTTGCGTCAGGGTTCT

ApB10_HCR_P2B3: GAAAGACCAGTCTTTTGAGCCAGTTTTCCACTCAACTTTAACCCg

ApB11_HCR_P1B3: gTCCCTgCCTCTATATCTTTTTTTGCTCTTGCGTTTTGAAACCAT

ApB11_HCR_P2B3: CTCTTGCCTCATCATATTCCGACGCTTCCACTCAACTTTAACCCg

ApB12_HCR_P1B3: gTCCCTgCCTCTATATCTTTGATGTCCCAGGTTGCCGTTCATCAT

ApB12_HCR_P2B3: CGCCGCCGACCAACGCCCCGTTTGCTTCCACTCAACTTTAACCCg

#### fringe (fng)

>fringe_LOC112056524_B3

Link: https://www.ncbi.nlm.nih.gov/nuccore/XM_024096965.2

ATGAGTGACAAGGTGCGAAGACTTGGCCAAGATTATCAGCGCTGTGGCGCCGGC TCTGGCATGGGTGGGCGTAGGATAATCAAAGCTGCGGCGCTACTCATAGCTTTGG GATACTGCAGTTTGCTTGTTTATCAAGGCGGAGTGAATTACAACTTCCAAGAAAG TCGGCCAGTGCAAGTGGCGGACTTACCTATTGAGCCCGTGACGAAAACGAGTAT TTTTGATAGTGGTCAGAGTAAGAGCATAACATTGGATGACATTTTCATAAGTGTAAAGACAACTAAGTTTTATCAGTATTCGAGGTTGCCGATAATTTTGAAGACATGGTTTCAATTGGCGAAAGAACAGACATGGTTTTTCACCGACACAGACAACCCACGACACCAAAATCAAACCAACGGTCACATGGTAAACACGAAGTGCTCGGCGTCACACCAGCTAAAACACCTCTGTTGCAAAATGTCCGTGGAGTACGATCATTTTCTCGAGAGCGGTAAAAAGTGGTTTTGTCACTTCGACGATGACAACTACGTCAATATTCCGCGCTTGGTCGCCGTGCTGCAGAAGTACAATCATCAAGAAGACTGGTATCTCGGGAAAACTTCTGTGAACAAACCTGTAGATATGTTCAAGAAGCCTACCAACGAGTTATTATACACTTTTTGGTTCGCAACCGGTGGTGCCGGAATATGTCTAAGCAGGAGTTTAGC TTTAAAAATGTTACCGATTGCAAGTGGTGGCAGATTCATCAGTATCTGTGAGAGT ATTCGATTTCCGGATGATGTCACTTTGGGTTATATAATAGAACATCTAATGGAGA AAAACTTGACCCGATTGCCAGAACTACATTCACATTTGGAGCAAATGAAACTACT GAATCCTGAAACTTTTCGAGACCAAATATCGTTCAGTTACTCAAAAACAGCCAAA GGTTGGAACGTCATCAATGTTCCCGGATTCAACCATAGATATGATCCCACTAGGT TCCTCTCGCTGCATTGTTTCCTCTTTCCACACTTCAAATTTTGTCCGAGATAG

Fng1_HCR_P2B3: TCTTCAAAATTATCggCAACCTCgATTCCACTCAACTTTAACCCG

Fng1_HCR_P1B3: GTCCCTGCCTCTATATCTTTACTgATAAAACTTAgTTgTCTTTAC

Fng2_HCR_P2B3: TgATTTTggTgTCgTgggTTgTCTgTTCCACTCAACTTTAACCCG

Fng2_HCR_P1B3: GTCCCTGCCTCTATATCTTTTCggTgAAAAACCATgTCTgTTCTT

Fng3_HCR_P2B3: CATTTTgCAACAgAggTgTTTTAgCTTCCACTCAACTTTAACCCG

Fng3_HCR_P1B3: GTCCCTGCCTCTATATCTTTgTgTgACgCCgAgCACTTCgTgTTT

Fng4_HCR_P2B3: CgTAgTTgTCATCgTCgAAgTgACATTCCACTCAACTTTAACCCG

Fng4_HCR_P1B3: GTCCCTGCCTCTATATCTTTACCACTTTTTACCgCTCTCgAgAAA

Fng5_HCR_P2B3: gTTTTCCCgAgATACCAgTCTTCTTTTCCACTCAACTTTAACCCG

Fng5_HCR_P1B3: GTCCCTGCCTCTATATCTTTTgATTgTACTTCTgCAgCACggCgA

Fng6_HCR_P2B3: TgCgAACCAAAAAgTgTATAATAACTTCCACTCAACTTTAACCCG

Fng6_HCR_P1B3: GTCCCTGCCTCTATATCTTTgTTggTAggCTTCTTgAACATATCT

#### bowel (bowl)

>brother of odd with entrails limited_LOC112044995_B3 Link: https://www.ncbi.nlm.nih.gov/nuccore/XM_024081020.2

ATGTCGGACAACGGTGGAACAGCTGTATCCCGGGCCTCTACACCTCGGCCAGTTTCCGAAACCCCTGCGAATCCCCCTCTACGCTCTAGACATGACCAGTTTCTCGCAAGTACTGCTGAAAGGGCTCAAAGACCTTCATCTAGTGGCCGAAATCCCAGTGCCAATTTCGACTCACCACCTTTACCATACCATCGTCAGATAACCACTGGCTCTCTTCTCACGGCTTCCCATCGTGAATCATCTGCCTTTGTTCCCGTAGTCCCAACAAGAGCCATGCACCCAGTTATGTACCCTAATGAAATTCACCCTCCCTTATTACCTTCAGAAATCATAGAACGCGAGCGTATTCTAGAAAGAGATAGAAGTGAATCAGCAAAAGGATCTCCAGATAGAGCATCGGCTAACTTTCAAAAAAGGAATTCATTTGACTTAATGGCTATGATGGTAGAAAAGAGAAAGGAGGTAGCATTGCGGGAAGCTGCAGCTGCCATGTTG ATGCCTCATCATAGAACTTCGTCAATTGATGGAATGATATCTGATGGGTCATCACAACCTCCAATTTATGGTCCACCTGGTGCATTCCTCGGTGCACCGGGTCCTTCGCCAACTGCTGCCAATAGTTTCACCTTCCCAGGTGCTGGGCTATTTCCTCCCGGGGCTGGTCCTCATCAAATGCATCCACATCTAGACAGACGACTACTCAGAGCCCCAGGCAGAGCGTCGAGGCCTAAGAAACAGTTCATCTGTAAATTTTGCAACCGACAATTCACGAAGTCCTACAATTTGCTTATCCACGAAAGAACACACACCGACGAGCGTCCCTATTCCTGTGATATTTGTGGGAAGGCATTCAGAAGACAGGATCACTTACGAGATCATAGGTACATACATTCAAAAGAGAAACCCTTTAAGTGCACAGAATGTGGAAAAGGATTCTGCCAGTCGCGGACTTTAGCTGTCCACAAGATTCTCCATATGGAAGAATCTCCCCACAAGTGTCCTGTCTGCAGCCGTAGTTTCAACCAGCGCTCGAATCTCAAAACACACCTTCTAACCCATACTGACCACAAACCTTATGAATGTAATTCATGTGGAA AAGTATTCAGAAGGAACTGTGACTTGAGAAGGCATGCTTTAACACATGCTGTAGGAGACGTTCCAGCAGGTGACGTATTGGATGTAGGTGAAGAAGATATTGGTAGACCTGGTTCACCTAATGAACCCGGTTTCGATGACGATGACCCTGAATTGTCATCTCCAGAACATTCTCCAGTAAGAAGAGCACGTTCTTCATCAATCGAAAGCATTGGAAGAGAAAAAAGTGAGGAAAGACCACGCCCTGTATCTCCGTCCCCAGTTGAACGCACACACTGCCACCATAATGAAGCCAGAGATAGGGCGAGTCCTTATACAATGCGACCTCAACATTTTGACAAGCATCGAATGATGTCGTCAGATATTTATGAAAAAAGACGATACACAACAGACGAAATAATAGAAAAAGAAATGCGCTACGTTGATATACCTGAACATCAAGTTCGTCATCCACAATTACAGATTCGTCGAGATTTACACCAAGTACCACCGCCTGATCCCTCGAAAATCAATCAATTGGCTAGCCCACCAATAGATCCTGGGCCTACAGGGATGTATTTAAGTCCATTTAGAAAAAGAAGTCATCCACCCGATATCGGTGATAATGTACGAACTTGCGCAGCTGTCTATAGAAGTCGATCACCAGAACCCACAAACTTGACAATGCCCAGACATGCGATTGGTAATAGTGATGTCGTAATTGGCATACCACCGACAATGATGGCAATGCCGCCATACCATAAGTTTGGAATGCCATTGCCCCCGCCACATACACAAATTCCGTATGGTGCAGTGCAACATTCTGTTCTTGGGCCTCAACAAACGGTAGCCCAAGATTTAATTGTGAAACACGCTCCACCAAAAGATACAATACCGAGCACTTCAACAAGCGTAACAAATTGTAAAGACACAACTCAAAACATCATGAAAGATAATCTAGATACAGACTCAAATTCGATTGCTAATGGAATCAAAAATATTGTTGGCATACAGCCAAAAATTAACCCCAAACCGGGAACAAGTCAATCAAGTGGTCCTAAAAAAGGCTTTAGCATAGAAGATATAATGAGACGTTAA

Bowl1_HCR_P1B3: GTCCCTGCCTCTATATCTTTATggCTCTTgTTgggACTACgggAA

Bowl1_HCR_P2B3: ATTTCATTAgggTACATAACTgggTTTCCACTCAACTTTAACCCG

Bowl2_HCR_P1B3: GTCCCTGCCTCTATATCTTTACgCTCgCgTTCTATgATTTCTgAA

Bowl2_HCR_P2B3: TgATTCACTTCTATCTCTTTCTAgATTCCACTCAACTTTAACCCG

Bowl3_HCR_P1B3: GTCCCTGCCTCTATATCTTTTTTTTTgAAAgTTAgCCgATgCTCT

Bowl3_HCR_P2B3: TCATAgCCATTAAgTCAAATgAATTTTCCACTCAACTTTAACCCG

Bowl4_HCR_P1B3: GTCCCTGCCTCTATATCTTTggggTTTCggAAACTggCCgAggTg

Bowl4_HCR_P2B3: TgTCTAgAgCgTAgAgggggATTCgTTCCACTCAACTTTAACCCG

Bowl5_HCR_P1B3: GTCCCTGCCTCTATATCTTTAggTCTTTgAgCCCTTTCAgCAgTA

Bowl5_HCR_P2B3: ggCACTgggATTTCggCCACTAgATTTCCACTCAACTTTAACCCG

Bowl6_HCR_P1B3: GTCCCTGCCTCTATATCTTTTggTTATCTgACgATggTATggTAA

Bowl6_HCR_P2B3: gATgggAAgCCgTgAgAAgAgAgCCTTCCACTCAACTTTAACCCG

#### wingless (wg/wnt1)

>wingless_LOC112058530_B1

Link: https://www.ncbi.nlm.nih.gov/nuccore/XM_024099417.2

ATGTCGGGTCCGCCCATAATGAAGTGGTTGTGCTTGTTTGTGCTGTTTCTGTGTAT GAGGTGCGAGGCCAACAAGCCGAGGCGAGGCCGAGGCAGCATGTGGTGGGGCA TAGCAAAGGCAGGCGAACCAAATAACTTATCACCCTTGTCTCCAAGTGTCCTATACATGGACCCGGCTGTTCACGCCACCTTGAGAAGGAAACAGAGAAGGCTAGCGAGGGAGAACCCTGGGGTTCTCGCAGCAATATCCAAGGGAGCCAGCATGGCTGTGGCCGAATGCCAGCATCAGTTCAAATACAGGAGATGGAACTGTTCTACAAGAAATTTTTTGCGAGGGAAGAATCTATTTGGAAAAATTGTTGACAGAGGTTGCCGTGAAACCGCCTTCATCTACGCCATCACAAGCGCGGGGGTGACGCACGCGGTGTCGCGCGCATGCGCCGAAGGTTCCATCGAGTCCTGCACGTGCGACTATTCTCATGTGGACCGTTCGCCGCACCGCGCGCGCGCCGCCGCCGCCGCCAACGTGAGGGTCTGGAAATGGGGCGGGTGCAGCGACAACATCGGCTTCGGCTTCAAGTTCAGCCGAGAGTTCGTTGACACCGGGGAAAGGGGCAAGACGCTTAGGGAGAAGATGAACTTGCACAACAATGAGGCTGGCAGAATGCACGTGCAAACGGAGATGCGCCAGGAGTGCAAGTGCCACGGTATGTCTGGGTCCTGCACGGTGAAGACGTGCTGGATGAGGCTGCCGACGTTCCGGTCTGTAGGCGACGCCCTGAAAGACAGCTTCGACGGGGCGTCGCGGGTCATGATGCCCAATACCGAGGTGGAGGCGCCGTCGCAGAGGAACGACGCCGCACCTCACAGGGTCCCGCGCCGTGACCGCTACAGGTTCCAACTTCGGCCGCACAACCCTGACC ACAAAACACCCGGGGTCAAGGACCTTGTATACTTGGAATCTTCACCAGGTTTCTG CGAAAAGAACCCCAGACTGGGCATCCCGGGTACGCACGGGCGTGCCTGCAACGA CACTAGCATCGGCGTCGACGGTTGCGACCTGATGTGCTGCGGGCGCGGCTACCG GACCGAGACCATGTTCGTAGTGGAACGATGCAACTGCACGTTCCACTGGTGCTGC GAGGTCAAATGCAAATTGTGTCGCACGGAAAAGGTAGTTAACACGTGTTTATAG

Wg1_HCR_P1B1: gAggAgggCAgCAAACggAATGCGAGAACCCCAGGGTTCTCCCTC

Wg1_HCR_P2B1: AGCCATGCTGGCTCCCTTGGATATTTAgAAgAgTCTTCCTTTACg

Wg2_HCR_P1B1: gAggAgggCAgCAAACggAAAGTTCCATCTCCTGTATTTGAACTG

Wg2_HCR_P2B1: TCCCTCGCAAAAAATTTCTTGTAGATAgAAgAgTCTTCCTTTACg

Wg3_HCR_P1B1: gAggAgggCAgCAAACggAAGTTTCACGGCAACCTCTGTCAACAA

Wg3_HCR_P2B1:GCGCTTGTGATGGCGTAGATGAAGGTAgAAgAgTCTTCCTTTACg

Wg4_HCR_P1B1:gAggAgggCAgCAAACggAAGGAACCTTCGGCGCATGCGCGCGAC

Wg4_HCR_P2B1:AGAATAGTCGCACGTGCAGGACTCGTAgAAgAgTCTTCCTTTACg

Wg5_HCR_P1B1:gAggAgggCAgCAAACggAACGGCGGCGGCGGCGCGCGCGCGGTG

Wg5_HCR_P2B1:CGCCCCATTTCCAGACCCTCACGTTTAgAAgAgTCTTCCTTTACg

Wg6_HCR_P1B1:gAggAgggCAgCAAACggAATCTCGGCTGAACTTGAAGCCGAAGC

Wg6_HCR_P2B1:TTGCCCCTTTCCCCGGTGTCAACGATAgAAgAgTCTTCCTTTACg

Wg7_HCR_P1B1:gAggAgggCAgCAAACggAAGCCAGCCTCATTGTTGTGCAAGTTC

Wg7_HCR_P2B1:GCGCATCTCCGTTTGCACGTGCATTTAgAAgAgTCTTCCTTTACg

Wg8_HCR_P1B1:gAggAgggCAgCAAACggAATCACCGTGCAGGACCCAGACATACC

Wg8_HCR_P2B1:ACGTCGGCAGCCTCATCCAGCACGTTAgAAgAgTCTTCCTTTACg

Wg9_HCR_P1B1: gAggAgggCAgCAAACggAAGCCCCGTCGAAGCTGTCTTTCAGGG

Wg9_HCR_P2B1:TCGGTATTGGGCATCATGACCCGCGTAgAAgAgTCTTCCTTTACg

Wg10_HCR_P1B1:gAggAgggCAgCAAACggAACCTGTGAGGTGCGGCGTCGTTCCTC

Wg10_HCR_P2B1: GAACCTGTAGCGGTCACGGCGCGGGTAgAAgAgTCTTCCTTTACg

#### decapentaplegic (dpp)

>decapentaplegic_LOC112044871_B2

Link: https://www.ncbi.nlm.nih.gov/nuccore/XM_052883655.1

ATGCGTGGGGCGTGCGCGTGCGCGGTGGTGTGCGCGTTGGTGGCGCTGTGCGCG GCGCGGCTGGACGAGTCCGCGCGCGCCGCCGCCGAGAAGCAGCTGCTGGCACTG CTGGGCCTGCCGCGCCGGCCGCCGCCGCGCGCCCGCCCGCCGCCCGTGCCGCGCGCGCTGCGCGTGCTGTACGACTCGCGCGCGCTGCCCGCCGCCGCCGCCAACACGGCGCGCTCCTTCCACCACACGCCCACGCCGCTCGACGAGCGCTTCCCCGGCGACCACCGCTTCCGCCTGTTCTTCAACGTGAGCGGCGTACCGGCCGACGAGGTGGCGCGCGGCGCCGACCTCTCGTTCCAACGAGCCGTCGGCACCACCGGCAGACAGAGACTGTTGTTGTACGACGTGGTGCGCCCCGGCCGCCGCGGCCACTCCGAGCCGATCCTGCGGCTGCTGGACTCCGTGCCGCTCCGGCCCGGGGAGGGAATCGTCAACGCCGACGCTCTGGGAGCGGCGCGACGGTGGCTCAAAGAGCCCAAACATAATCACGGACTATTAGTGCGAGTGTTAGAAGAAGACGCCGCGAGTGTGAGCAGGGACGCGAAGTTCCCGCACGTGCGCGTGCGCAGACGCGTCACGGACGAGGAGGAGGAGTGGCGGACGGCGCAGCCGCTGCTCATGCTGTACACGGAGGACGAGCGCGCGCGCGCGTCGCGGGAGACGAGCGAGCGGCTGACGCGCAGCAAGCGCGCGGCGCAGCGGCGGGGGC ACCGCGCGCACCACCGCCGCAAGGAGGCGCGCGAGATCTGCCAGCGCCGCCCGC TGTTCGTCGACTTCGCGGACGTGGGCTGGAGCGACTGGATCGTGGCCCCGCACG GCTACGACGCGTACTACTGCCAGGGCGACTGCCCCTTCCCGCTGCCGGACCACCTCAACGGCACGAACCACGCGATAGTGCAGACTCTGGTCAACTCAGTGAACCCCGC GACGGTGCCCAAAGCGTGCTGCGTGCCGACGCAACTCTCATCTATATCTATGTTA TATATGGACGAAGTGAACAATGTGGTGCTTAAAAACTATCAGGACATGATGGTG GTAGGCTGTGGCTGCCGATGA

Dpp1_HCR_P1B2: CCTCgTAAATCCTCATCAAAGGGCGTGTGGTGGAAGGAGCGCGCC

Dpp1_HCR_P2B2: GCCGGGGAAGCGCTCGTCGAGCGGCAAATCATCCAgTAAACCgCC

Dpp2_HCR_P1B2: CCTCgTAAATCCTCATCAAACCGGTACGCCGCTCACGTTGAAGAA

Dpp2_HCR_P2B2: GGTCGGCGCCGCGCGCCACCTCGTCAAATCATCCAgTAAACCgCC

Dpp3_HCR_P1B2: CCTCgTAAATCCTCATCAAAAGTCTCTGTCTGCCGGTGGTGCCGA

Dpp3_HCR_P2B2: CCGGGGCGCACCACGTCGTACAACAAAATCATCCAgTAAACCgCC

Dpp4_HCR_P1B2: CCTCgTAAATCCTCATCAAAGGAGTCCAGCAGCCGCAGGATCGGC

Dpp4_HCR_P2B2: GATTCCCTCCCCGGGCCGGAGCGGCAAATCATCCAgTAAACCgCC

Dpp5_HCR_P1B2: CCTCgTAAATCCTCATCAAACTTTGAGCCACCGTCGCGCCGCTCC

Dpp5_HCR_P2B2: CTAATAGTCCGTGATTATGTTTGGGAAATCATCCAgTAAACCgCC

Dpp6_HCR_P1B2: CCTCgTAAATCCTCATCAAAGCGTCCCTGCTCACACTCGCGGCGT

Dpp6_HCR_P2B2: CTGCGCACGCGCACGTGCGGGAACTAAATCATCCAgTAAACCgCC

Dpp7_HCR_P1B2: CCTCgTAAATCCTCATCAAACGGCTGCGCCGTCCGCCACTCCTCC

Dpp7_HCR_P2B2: CTCGTCCTCCGTGTACAGCATGAGCAAATCATCCAgTAAACCgCC

Dpp8_HCR_P1B2: CCTCgTAAATCCTCATCAAATGCGCGTCAGCCGCTCGCTCGTCTC

Dpp8_HCR_P2B2: GCCCCCGCCGCTGCGCCGCGCGCTTAAATCATCCAgTAAACCgCC

#### vein (vn)

>vein_LOC112058144_B3

Link: https://www.ncbi.nlm.nih.gov/nuccore/XM_024098831.2

ATGAAGACGTGCGGGCGCGCGGCGTTTTGCTGGGCCCTGCTGTGCCTCGCTGGCA GCCTGGCGGCGCCGCGCCGCCTCTACTGCGAGCGGGCGGACGTCGCCTGGCGCGCCTACCGAGCCCCCGCCGCCTTCGAGGCGCGCGTGCAGTCGCTGGCGCGCGACGCCGCCACCGTGCAGGTGGGCCGCGTGCTGCGGCGCCAGGGCCGCTGGCCGCGCGAGGACGCCGTGCTGCGCCTCAAGCTGCCCGAGCAGCCCCGCGAGTGCGCCGGACGCTTCGAAGTGCCGCTGCAGAACCGAAAGAACTATATCGTGTTCGCGGAGAGGCAGGGCCACGCCGCCGTGGCGCTGGGCCCGCCGCTGCGCAGGACTGCCAAGTTGATCCGGAGGATCCGCGCGGTCTACCGGCCCGGCTACAGCAGCCCAGCGCGGGTGGAGCCGATGCAGTCGGTGCGCGCGACGCGCGGCGGCAAGGTCCGGCTGGAGTGCCGCGCCAGCGCGCGCCCGCCGCCGCGGATAGCCTGGTACAAGGACGGCCGGCCCGTCGCAGACATTGCTTTAAGGAGGTTCCGAGTGCAGAACTATAGGCGGCGCTCGGTATTAGTAATAAGACATGCGCGTCGCGAGGACACCGCGACGTACGAATGTAGGGCGCAGGGTGCGGTGGGGCCGCCGGCCGTCGCCGCCGCCAACGTCAGCGTCCTACCACCGGCCACGATGGCGCCAGATACCCCCCCCGGCGCTCCGTGTCCGATGCCCGATCCTGCCTCTTACTGTCTCAATGGAGGCACTTGTTTATTCTTCGAGCTTGTTCAGGAACAAGCTTGCAAATGCCCGGAAGGCTTCACCGGGCAACGCTGCGAGATCAAG GACGTTTTGAATAGGAACAGCTTGGGAGAACGACTATGGTGA

Vn1_HCR_P1B3: gTCCCTgCCTCTATATCTTTGTGGCGGCGTCGCGCGCCAGCGACT

Vn1_HCR_P2B3: CGCCGCAGCACGCGGCCCACCTGCATTCCACTCAACTTTAACCCg

Vn2_HCR_P1B3: gTCCCTgCCTCTATATCTTTCTTGAGGCGCAGCACGGCGTCCTCG

Vn2_HCR_P2B3: GGCGCACTCGCGGGGCTGCTCGGGCTTCCACTCAACTTTAACCCg

Vn3_HCR_P1B3: gTCCCTgCCTCTATATCTTTCGATATAGTTCTTTCGGTTCTGCAG

Vn3_HCR_P2B3: CGGCGTGGCCCTGCCTCTCCGCGAATTCCACTCAACTTTAACCCg

Vn4_HCR_P1B3: gTCCCTgCCTCTATATCTTTATCAACTTGGCAGTCCTGCGCAGCG

Vn4_HCR_P2B3: GGCCGGTAGACCGCGCGGATCCTCCTTCCACTCAACTTTAACCCg

Vn5_HCR_P1B3: gTCCCTgCCTCTATATCTTTCACCGACTGCATCGGCTCCACCCGC

Vn5_HCR_P2B3: CCGGACCTTGCCGCCGCGCGTCGCGTTCCACTCAACTTTAACCCg

Vn6_HCR_P1B3: gTCCCTgCCTCTATATCTTTAGGCTATCCGCGGCGGCGGGCGCGC

Vn6_HCR_P2B3: CTGCGACGGGCCGGCCGTCCTTGTATTCCACTCAACTTTAACCCg

Vn7_HCR_P1B3: gTCCCTgCCTCTATATCTTTCGCCGCCTATAGTTCTGCACTCGGA

Vn7_HCR_P2B3: CGCGCATGTCTTATTACTAATACCGTTCCACTCAACTTTAACCCg

Vn8_HCR_P1B3: gTCCCTgCCTCTATATCTTTCGCACCCTGCGCCCTACATTCGTAC

Vn8_HCR_P2B3: GGCGGCGGCGACGGCCGGCGGCCCCTTCCACTCAACTTTAACCCg

Vn9_HCR_P1B3: gTCCCTgCCTCTATATCTTTGGGTATCTGGCGCCATCGTGGCCGG

Vn9_HCR_P2B3: CGGGCATCGGACACGGAGCGCCGGGTTCCACTCAACTTTAACCCg

Vn10_HCR_P1B3: gTCCCTgCCTCTATATCTTTAAGAATAAACAAGTGCCTCCATTGA

Vn10_HCR_P2B3: TTGCAAGCTTGTTCCTGAACAAGCTTTCCACTCAACTTTAACCCg

### Appendix D: HCR Overview

**Figure S4:**
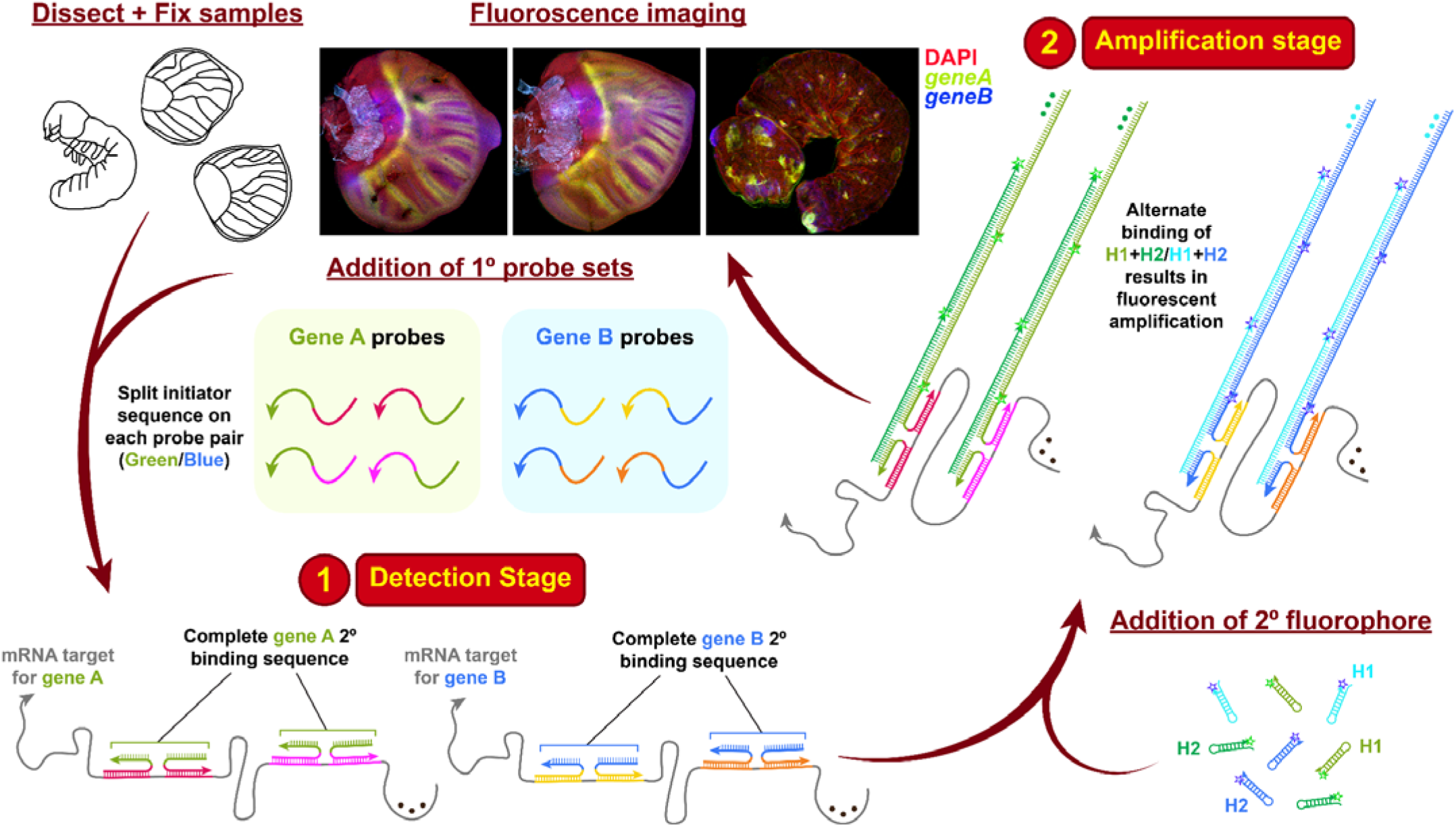
Hybridisation chain reaction v3.0 overview (adapted from Choi *et al*., 2018). The two main principles of this technique are (1) Detection and (2) Amplification, using split initiator 1° probes with different H1+H2 2° fluorescent hairpins for improved background suppression and high resolution multiplexing.

