## Supplementary material for "Parallels in leg and wing proximal-distal patterning in Diptera and Lepidoptera": Lee et al. 2026_Supplementary Information

### **Supplementary materials for “Parallels in leg and wing proximal-distal patterning in Diptera and Lepidoptera”**

#### **Appendix A: List of APG paralogs and gene families.**

Table S1: **List of APGs in *Bicyclus anynana* with relevant paralogs or gene family members.** All paralogs were identified through OrthoDB (v12.2) (<https://www.orthodb.org/>) and/or OrthoFinder v3 (<https://github.com/OrthoFinder/OrthoFinder>). The genes highlighted in yellow were tested in this manuscript.

| **Gene/Gene family name** | **Paralog/**  **member used** | **LOCID** | **Other paralog(s)/**  **member(s)** | **LOCID** |
| --- | --- | --- | --- | --- |
| *apterous* | *apterousA* | LOC112045866 | *apterousB* | LOC112045833 |
| *aristaless* | *aristaless* | LOC112056678 | *aristaless* | LOC112054675 |
| *odd-skipped* family | *bowel* | LOC112044995 | *zinc finger and BTB domain-containing protein 49* | LOC112045928 |
|  |  |  | *zinc finger protein 846* | LOC112054472 |
|  |  |  | *sister of odd and bowel* | LOC112044996 |
|  |  |  | *drumstick* | LOC112045021 |
|  |  |  | *odd-skipped* | LOC112044997 |

#### **Appendix B: Supplementary expression of APGs in legs and wings**

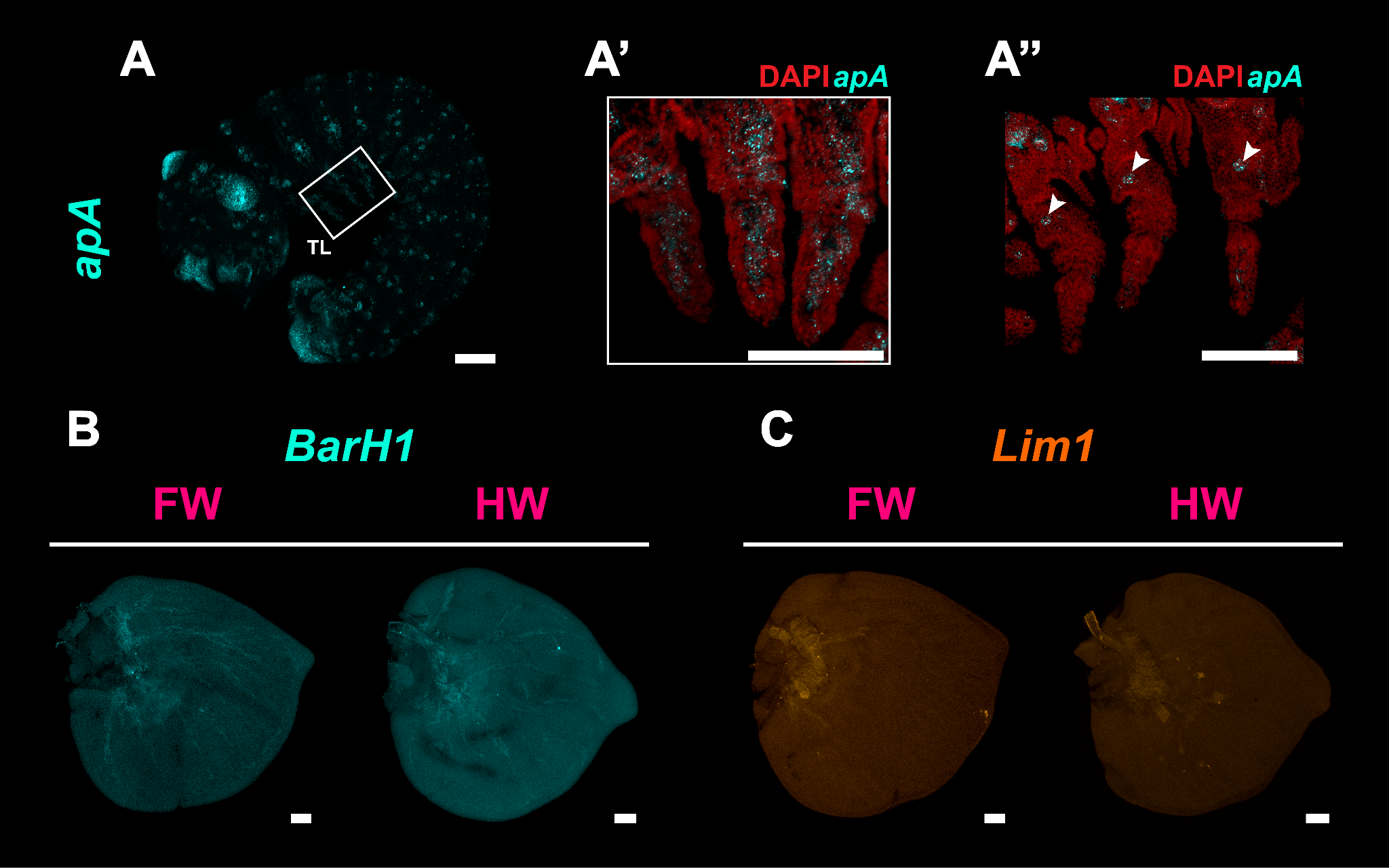

Figure S2: **APGs with no expression in legs and wings.** All wings are mid to late larval, and each subfigure consists of a forewing (FW; left) and hindwing (HW; right). **(A, A’)** *apA* is strongly expressed in mesodermal cell types throughout the centre of the leg. **(A’’)** Epidermal slice of an embryonic leg, where *apA* is not expressed. A small cluster of *apA-*expressing cells is observed at each thoracic segment, located in the ventral body wall and not at the proximal domain of the leg (white arrowheads). **(B)** *B-H1* and **(C)** *Lim1* are not expressed in the larval wings. Scale bars: 100 µm.

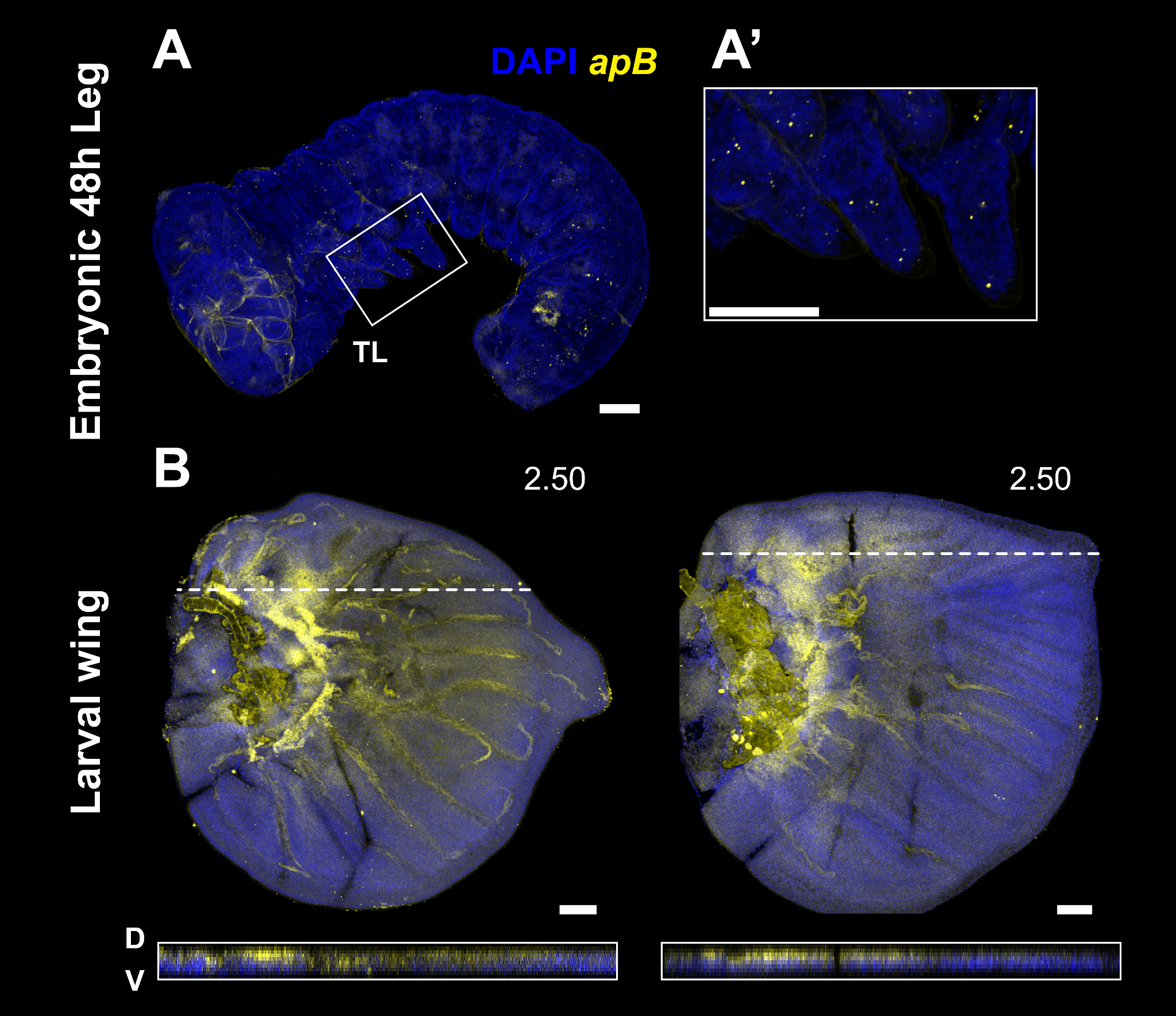

Figure S3: **Expression of *apterous* paralog *apB* in legs and wings. (A)** *apB* was not expressed in the TL, (A’) is a closeup of the legs from (A). **(B)** *apB* expression in the larval fore (left) and hind (right) wing. A DV cross-section of the white dashed line is displayed below each wing. While *apB* spatial expression domain is similar to that of *apA*, its expression is comparably more muted. Scale bar: 100 µm.

***tiptop* (*tio*)**

>tiptop_LOC112051061_B1

Link: <https://www.ncbi.nlm.nih.gov/nuccore/XM_024089545.2>

ATGATCGTCGAGCCAGACCACGTGAGGGAGAGTCCAAGATGTTTGTCACGGGAGTCGTCCGGCGCGCCACGATGTCCCTCCAACGACTCCGTGTACTCGGGTCGGAGCGCGCCGAGTCTGCCTCTGCCGGCTGCGTTGCAAGCGGCGCTCCCGGCAGGTCTCCCCGCGGCGCTCCTGCCCCCCCACTCCGCTGCCGTGGCGGCCTACCTCGGCGCAGCCGCCGCCGCCGCCCAGCAGAGGCTCCTCATGTCCTGCCAAGAAGATATGACAGACTCAGAACGCGCTGACGCAGTACTAGACTTCAGCACGAAGCGAAGTGAATCCCCCATCGATGATGAAGACGACGATGCCGTCAATCTGAGCAAAAACGAAAACGGCCCCTTAGATTTATCAGTAGGCACCAGAAAACGAGGACCACAAAATTCTCCATCACCCGTTCCTACGAGAAAAAGTTCACGGACGGCTGATTTTAAACCGATCACTACTCCGTGGACTACGCCGGTTGCGCCACATCTTCCTTATTTTGCAGCAGCTGTTGCAGCTGCTAGCTTATCACCTAAAGGTGGTGTCCCACCAGACTGGAATGGAAAATTAAAACATGGGACACCGACACCTAACGATGCAACTAAAGCGCTTGAAAAAATGAGCGAGTTAAGTAGGTTAGGTGGAGAGGAGTTATTTAGGTCAGTTCAAAGCGCAGCTTTAGGAGCCGGGCTCACTCCTAACGCAGCCGCTCGCCACTCTGCCTGGCAGTCGCATTGGTTGAATAAAGGCGCCGACCAAACGAAAGACGTTTTGAAATGTGTGTGGTGCAAGAAAAGTTTTAACTCTTTAGCTGATTTAACTGTTCATATGAAAGAAGCTAAGCATTGCGGCGTGAACGTCCCGGTACCACCATCGGGCGGTGCACCGATCCCTCCTTCACTTCTGCCGCCTTCTAGTTCACCTTCGACGCCATCTTACAACTCTTCTTCGTCGAGCGGATCGTCTAAACCTAATCACAACGATCTTAATATGCTAATTAAGGAAAACATGCCCATACCTAGAAAACTAGTCCGCGGCCAAGACGTATGGTTAGGTAAAGGGGCCGAACAAACTAGGCAAATATTAAAATGTATGTGGTGCGCCGAAAGTTTCCGCTCTTTAGCCGAAATGACGAGCCACATGCAGCGCACCCAGCACTACACTAATATTATATCACAAGAACAAATCATTTCTTGGAAGTCTTCAGATGAAGCCAAGGGCTCCAACTCTAGCGCACCCGGTACTAATAACGCGGTTCCTCCGACGACAGGGACGAGTAGTCACGTTAGCGCTGTTTTAACTTGCAAAGTATGCGACCAAGCGTTTAGTTCCTTAAAAGAATTAAGTAATCATATGGTTAAGAACTCTCACTATAAAGAGCACATTATGCGCTCGATAACTGAAAGCGGTGGCAGACGACGACAAACGCGCGAAAAGAGAAAGAAGTCCTTGCCTGTGAGGAAACTTCTCGAGCTCGAGAGAGCCCAACATGAGTTTAAAAATGGGGAGGGCAACGGAGTCCCTATGGGAAAACCTATTAGGGATTTTGGCTCTGGAAGCCGTATCACTTGCGAGAAATGCGGCGACAAAATTGAGACTGCTGTATTCGTAGAACACATCCGACAATGCATAGGCGCGCCCATGTCGAATAGCCAAAGGAATTTTCTAAAAAGTGCTCTACTTTCCAATAATATTATTCCACCCGACGTGCCCGGCCACATCACTCCCACCAGTCGCGACGGACGTAAGAGTATCAATGACGAAATTCCCTCACCAGGGTCGACGCATCACCGTTCACCTTCTTCGGTTAACGATTCTTCTCCTAGTTCTAAAGATCACAACGTCAGCAACGATAAAAGCTCATCACCATCCGTACTCAATGCCATAGAACAATTAATAGAGAAGAGTTTCGATACTCGCTCGCGACATTCGGTCCCCGGTATGCCAGGCGGAGCTTCCCACGCCCCTATTGGTTCTAGTATTTTAAAAAGATTAGGTATAGACGAGAGCGTAGACTACACGAAACCTCTGGTAGATCCACAAACGATGAGCATGTTGAGAAATTATCACCACCAGCAAGGCTACGGTCGTCGAGAGCGCAGCGGTAGCGAATCTAGTTCCATGTCCGAGAGGGGCGGCAGTAGGGTGGAGTCTCTCACGCCGGACAGAAAGTTGGATTCGTATCATATGACCCCGCGTACGACTCCCGACACTCGGGGATCCCAGACCCCCGCTTCGGAGGATCGCCCCGCAGAGGTCAGGATAAAAAAAGAAATCACCGACGAAGAAGATAGGGAGACTTGCGTAGACTTGAGCAGTCAGCCGGTCAGGGTTAAAACCGAAGTGGACGATGACGAGGAAGCGAGGCCGAACAGTGTGGCGGACGAAGACGTGAAGCCGACCGTCCCCAAGCGCGAGAGCGAGGGTCCGAGCCCCGTGCCTAGCCCTCGCAGCCCGGCCAGCGACCGCTCGGCGCCCACTCCCGGCGCTGACAGAAAGCCGGCTTCCAGCCTAGGCGCGCTCTCCTCCATGTTCGACAACCTCGCTGGGGGAGCCTCCTCCAACGAGCCCAGCTCGTCTCGTCGCGGGGGTAGTCACCCCCTGGCGGCCTTGCAAAAACTATGCGACAAAACAGACACTAATTCTTCCCGCGCACCGCCCCCCGCGCCGTCCCCCGCCGGGCCGCCGAGTATTTTAACTTTCAGTTGGGCTTGCAACGACGCGGTAGTCACAGATTCGATAATGAAATGCGCTCTTTGCGACACACCGTTTATTTCGAAAGGCGCGTACCGGCACCACCTGTCCAAGATGCACTTCGTGAAGGACGGCGCACTGCCAGAGCCGGTGCCGGTGAAGGCGCCGCCGCCCGCGCCGTCGCCGGGCGCGCACAAGTCCGGCTCCACCGCCGCGTCGCCCCAGGACCCGCGGAGTCCGTCCCAGTCGTTCGACGAGAGCCCGCACTCCAAGTTCCTCAAGTATACGGAGCTGGCGAAGCAGCTGTCCAGCAAGTACGTGTAG

Tio1_HCR_P1B1: gAggAgggCAgCAAACggAACGGCTAAAGAGCGGAAACTTTCGGC

Tio1_HCR_P2B1: GGGTGCGCTGCATGTGGCTCGTCATTAgAAgAgTCTTCCTTTACg

Tio2_HCR_P1B1: gAggAgggCAgCAAACggAACAAGAAATGATTTGTTCTTGTGATA

Tio2_HCR_P2B1: GAGCCCTTGGCTTCATCTGAAGACTTAgAAgAgTCTTCCTTTACg

Tio3_HCR_P1B1: gAggAgggCAgCAAACggAACGTCGGAGGAACCGCGTTATTAGTA

Tio3_HCR_P2B1: AGCGCTAACGTGACTACTCGTCCCTTAgAAgAgTCTTCCTTTACg

Tio4_HCR_P1B1: gAggAgggCAgCAAACggAATTAAGGAACTAAACGCTTGGTCGCA

Tio4_HCR_P2B1: TCTTAACCATATGATTACTTAATTCTAgAAgAgTCTTCCTTTACg

Tio5_HCR_P1B1: gAggAgggCAgCAAACggAACTTTCAGTTATCGAGCGCATAATGT

Tio5_HCR_P2B1: TCGCGCGTTTGTCGTCGTCTGCCACTAgAAgAgTCTTCCTTTACg

Tio6_HCR_P1B1: gAggAgggCAgCAAACggAAGAGCTCGAGAAGTTTCCTCACAGGC

Tio6_HCR_P2B1: ATTTTTAAACTCATGTTGGGCTCTCTAgAAgAgTCTTCCTTTACg

Tio7_HCR_P1B1: gAggAgggCAgCAAACggAAAATCCCTAATAGGTTTTCCCATAGG

Tio7_HCR_P2B1: CGCAAGTGATACGGCTTCCAGAGCCTAgAAgAgTCTTCCTTTACg

Tio8_HCR_P1B1: gAggAgggCAgCAAACggAATGTTCTACGAATACAGCAGTCTCAA

Tio8_HCR_P2B1: ATGGGCGCGCCTATGCATTGTCGGATAgAAgAgTCTTCCTTTACg

Tio9_HCR_P1B1: gAggAgggCAgCAAACggAAGGAAAGTAGAGCACTTTTTAGAAAA

Tio9_HCR_P2B1: GGGCACGTCGGGTGGAATAATATTATAgAAgAgTCTTCCTTTACg

Tio10_HCR_P1B1: gAggAgggCAgCAAACggAATGATACTCTTACGTCCGTCGCGACT

Tio10_HCR_P2B1: TCGACCCTGGTGAGGGAATTTCGTCTAgAAgAgTCTTCCTTTACg

***dachshund* (*dac*)**

>dachshund_LOC112044740_B3

Link: <https://www.ncbi.nlm.nih.gov/nuccore/XM_052883449.1>

ATGGAGTCCGCCGTGGACTCAGCGTCGACCGCCAGCGAGGTGAGCGGCTCGTCCGGGGGCTCGCCGCGCGTCAAGGCGGCGTCGCCGGCGCGCGGCCTCAGCCCGCCGCAGCTGCTGGCGCCGCGCCTGCCGCTGCCGCCGCCGGGCCTGGGCCTGCTGGGCTCGCTGCAGATGATGCACCACTCGCCGCTCGAGCTCATGGCGGCGGCGCACCACCACCCGCACCGCTACGGGAGCCCGCCGCCCATCTCCACGTCCGACCCCTCGGCGAACGAGTGCAAGCTGGTGGACTATCGCGGGCAGAAGGTGGCCGCGTTCATCATCCAGGGCGACACGATGCTGTGCCTGCCGCAGGCCTTCGAGCTGTTCCTGAAGCACCTGGTGGGCGGGCTGCACACCGTGTACACCAAGCTGAAGCGGCTGGACATCGTGCCGCTGGTGTGCAACGTGGAGCAGGTCCGCATCCTGCGCGGGCTGGGCGCCATCCAGCCGGGCGTCAACCGCTGCAAGCTGCTCTCGTGCAAGGACTTCGACGTGCTGTACCGCGACTGCACCACGGCAAGGTGCCTGTCAATGAAAGCGCCAGACAGCTCCAGACCGGGTCGACCTCCGAAGCGTGCTTCTGGAGTTGGTCTTTCGCTTGCCGCCACGCAGTTCCCGGGGCACCCCTTCAAGAAGCACCGCCTAGAGAATGGGGACTACTCGCCGTATGAAAATGGACATATGAGCGAGATGGCCCGCATGGATAAGTCCCCGCTCCTCGCGAACGGGTACAACGCACCCCCCACCCACCTGGGGCCCATGGGCTTCATGCACCAGCACGCCCTCATGTCTCCGGGTATGCCACACCCTGGCGTGCCCAGGCCTGACGGGTCCATCATTAAGGGGCAGCCTATGCATAACATGGAAGCACTAGCAAGATCTGGTATTTGGGAGAATTGTAGAGCAGCATACGAGGACATCGTAAAACATCTCGAAAGGTTGCGTGATGAACGGGGTGATATTGAACGTGTTATAGCAATGGACAAGGCGCGCGAGGGTTCACATAATGGTTCGTCTCCAGGCCACAGTCCTGTGCTGAACCTTTCGAAGTCAGGCTCCGGTGAGCGCGAGCGGTCCGAGCGGGACCGCGAGCGCGCGGAGCGGGGCGAAGGCTCGGCGAGCGGGCGCAGCTCGGCCGCCTCGCGCCGCACGCCGCAGCCGCCGCGCATCCCCTCCACCGCCGCGCCGGTCTCCCCGCGCTCCCACTCCGACGAGAGCGACGCTGCTCTGTCTGATCAAGACGACCATAACGTCAAAGACGAGGATGACGATCTCAGCGATGGTGAACGAGACCTCGCAACGAATTCCTCGCCAGCGCCCGTCAGCTACCCCCCACAAGGCTCGCCGTCGAACGTTCCCGTGGACCCCACCGCCGACACCCTGGTCTCCTCCACCGAGACCCTCCTCAGGAACATCCAGGGCCTCCTCAAAGTAGCGGCGGACAACGCACGCCAACAAGAACGGCAAATCAGTTACGAAAAAGCGGAACTAAAAATGGACGTGTTAAGAGAAAGAGAAGTAAAAGACAACCTGGAAAGACAGTTACTGGACGAACAAAAAATGAGAGTAATGTATCAGAAGAGATTAAAGAAAGAAAGAAAGCAGAGGCAACAAATTCAAGACCAATTAGAATTGGAGTTAAAAAGACGACAGAAGATAGAAGAAGCACTAAAGCAGTCAGGAGCGCCCGGTGAAATTCTGAGAATAGTAACTGAGAATTTAACACCACCGTCCCAAGAAAATCGCGAACGTGAGAACGGTACAGAGAGCAAACCGCCAAGTACAGAGCCCCCCACCTCGTCGCCGCCGTTCCAGAGGGACCCCCCGCGCACGCCCGACAAGCCCCAGTGGAACTACCCTCCGCCGCCTGTCGACATCATGAGCGGAGGAGCTGCTTTCTGGCAGAACTACTCTGAATCCTTGGCGCAAGAGTTGGAGATGGAGCGCAAGTCCCGCCAGCAGGCCATGGAGAGAGACGTGAAGAGCCCTCTGTCGGACCGCGCCAGTTACTACAAGAACTCGGTGCTGTTCAGCTCGGCCACTTAG

***spineless* (*ss*)**

>spineless_LOC112043391_B2

Link: <https://www.ncbi.nlm.nih.gov/nuccore/XM_024078792.2/>

ATGCACAAAGAGAAGGAGGAGCTGCACCGGCGTGACCACGCGCACTACGACCACATCATGCCGGACGGCGAGATGTTCCTACAGGCACTCAACGGGTTCCTGATGATCCTCACGTGCGAGGGGGAGGTGTTCTTCGCGACACACAGCATCGAGAGCTACCTGGGCTTCCATCAGTCAGACATAGTCCACCAATCAGTGTACGAGTTGGTCCACTCAGAAGATAGAGAAGAGCTTCAGAGGCAGTTACTTTGGAACTCCTTCCTGCCACCAGAAGCAAACAGTCTCTCACTGCAAGATGTTTTGAGACAGGAGAGAGCGCATCTATTAGAAAGAAGTTTTACAGTCAGATTTAGATGTTTATTAGACAATACATCAGGATTTTTGAGATTAGACATAAGAGGCCGGATCAAAGTATTACATGGACAGAACAAGAAAACAGAAGAGCCACCATTAGCTTTATTTGCAATTTGCGCACCGTTCGGGCCTCCCAGCCTGCTGGAGATACCACAGAAAGAGGTCATGTTCAAGAGCAAACACAAACTGGATTTATCACTAGTCTCGATGGATCAAAGAGGGAAACTTCTCCTCGGTTACACGGACGCCGAGCTCGCAAACATGGGAGGTTACGATTTAGTGCATTACGACGACCTGGCGTATGTAGCCAGCGCTCATCAGGAATTACTGAAGACAGGAGCGTCAGGGATGATCGCATACCGCTTCCAGACGAAGGACGGCCAGTGGCAGTGGCTGCAGACTTCCTCGCGACTCGTCTACAAGAACTCCAAGCCTGACTTCGTGATTAGCACGCATAGGCCTCTAATGGAAGAAGAAGGCAGAGATCTCCTCGGGAAGCGGACGATGGACTTCAAAGTTAGCTATTTAGACACCGGTCTCACCAGCCTTTACTTTTCCGAAACAGAACAGCTTTCAGGCTGCGAGTCGAACGTCTCAACGCCCCCTCGAGCAGCGAGGCGATATAAGACTCAACTGAGAGACTTTCTATCCACTTGTAGGAATAAAAGACGAGTGCCTCCCACTCCCGCGCCACCTCCAGTGGATTACTTAGCTGCTGATGCGGTAGCCGCAGCGTATACTAATCTAAACGGAGCGTACACCGCTCCATACGGGGAATATCCTCCCGCTCCAGCGTTACATTACGCTCCACCACCTTTAGATGATAGATTCCTCGCAGCTGACAACCTCTTCCATCAATACAAACCATTGTCTTACCATTACACTCCATACGCGCCTAATGGTTTCTTAGAACCGGCTCCACCTCCTGGATACGAAGTTCCACCGACGTATCATCGACCTCCGTCAAGAGAATACACTTATGTTGATAGTAGTGGGAGGTACATGAGTCCGGTGACTGGAGAGAGACGGTCGCCAAATGTCGTGCCTGGTCCAAGTCCGGGTGGGTCAAGCGGTTCGACAGAGCAGGATAGGATGGTGCAAACGCAAACGTCGGAGATGCCAAGGCAGACGGTCCTCATGTGGGGTGCCGGTGGAGCAGCTCAAGAAGTCCCCGTGGAGTACTCACCACCGCAGGTTTGGCGATACCATCCCTCCCACTACCACACCGCTGAAGCGACTCAATGA

Ss1_HCR_P1B2: CCTCgTAAATCCTCATCAAATGACCTCTTTCTGTGGTATCTCCAG

Ss1_HCR_P2B2: AATCCAGTTTGTGTTTGCTCTTGAAAAATCATCCAgTAAACCgCC

Ss2_HCR_P1B2: CCTCgTAAATCCTCATCAAACCGAGGAGAAGTTTCCCTCTTTGAT

Ss2_HCR_P2B2: ATGTTTGCGAGCTCGGCGTCCGTGTAAATCATCCAgTAAACCgCC

Ss3_HCR_P1B2: CCTCgTAAATCCTCATCAAATACATACGCCAGGTCGTCGTAATGC

Ss3_HCR_P2B2: CTTCAGTAATTCCTGATGAGCGCTGAAATCATCCAgTAAACCgCC

Ss4_HCR_P1B2: CCTCgTAAATCCTCATCAAACCTTCGTCTGGAAGCGGTATGCGAT

Ss4_HCR_P2B2: AAGTCTGCAGCCACTGCCACTGGCCAAATCATCCAgTAAACCgCC

Ss5_HCR_P1B2: CCTCgTAAATCCTCATCAAAATCACGAAGTCAGGCTTGGAGTTCT

Ss5_HCR_P2B2: TCTTCCATTAGAGGCCTATGCGTGCAAATCATCCAgTAAACCgCC

Ss6_HCR_P1B2: CCTCgTAAATCCTCATCAAATTTGAAGTCCATCGTCCGCTTCCCG

Ss6_HCR_P2B2: GGTGAGACCGGTGTCTAAATAGCTAAAATCATCCAgTAAACCgCC

Ss7_HCR_P1B2: CCTCgTAAATCCTCATCAAAACTCGCAGCCTGAAAGCTGTTCTGT

Ss7_HCR_P2B2: CTGCTCGAGGGGGCGTTGAGACGTTAAATCATCCAgTAAACCgCC

Ss8_HCR_P1B2: CCTCgTAAATCCTCATCAAACAAGTGGATAGAAAGTCTCTCAGTT

Ss8_HCR_P2B2: GTGGGAGGCACTCGTCTTTTATTCCAAATCATCCAgTAAACCgCC

Ss9_HCR_P1B2: CCTCgTAAATCCTCATCAAATACCGCATCAGCAGCTAAGTAATCC

Ss9_HCR_P2B2: TCCGTTTAGATTAGTATACGCTGCGAAATCATCCAgTAAACCgCC

Ss10_HCR_P1B2: CCTCgTAAATCCTCATCAAAACGCTGGAGCGGGAGGATATTCCCC

Ss10_HCR_P2B2: CATCTAAAGGTGGTGGAGCGTAATGAAATCATCCAgTAAACCgCC

***rotund* (*rn*)**

>rotund_LOC112056645_B3

Link: <https://www.ncbi.nlm.nih.gov/nuccore/XM_024096578.2>

ATGATGGAAGGAAGGACGATCGATTACAGGCCAGACGGTGGTGGGCTCGATTACCATAACTCTCCGTTAATGGCGGAGATACCTGTAGACAATTACTCCCATATCCATAGAAGTATAGAGCATTTGAGGTCGATGGGAGTGGCACCACACCTGGATCCCCATAGACATTTAGCTGCGAATCTGACAGACTTGCGAAGGTACGAACACCCTGATGACATGCCTGAGATAAAGCCGTCCGTACTTAGATTATCCGAATTCAAAAGTGGTCTACAAGAAATAAGGGTGCACAATGACCAAGAGAATGACATTCAGCGTCAAATGACACCGCACGACGATGGAAAAGGTTACAGCGCACCTTCAACGCCTTTATCAGAAAACGGCGGACAAAGCATTCAAGAGGAAAAGGTGTTCGGCTCCAAAGCGGACCTCCAGCTGCACACGCAGATCCACCTGCGCGAAGCGAAGCCCTACCGCTGCACCCAGTGCCCGAAGGCCTTCGCCAACTCCTCATACCTAGCGCAGCATTCGCGCATCCACCTCGGCATCAAACCGTACCGCTGCGAGATCTGCCAGCGCAAGTTCACCCAACTGTCTCACCTGCAGCAGCACATCCGCACCCACACGGGTGACAAACCCTATCGATGCACCCAGATAGGGTGCACCAAAGCGTTCTCCCAGCTGTCCAACTTGCAGAGCCACAGCCGCTGCCACCAGACCGACAAACCCTTCAAATGCAACTCCTGCTACAAATGCTTCACGCACGAGAAAGACCTCCTCGAACACATTCCGAAACACAAAGAATCCAAGCATTTGAAGACCCACATCTGCCAGTACTGCGGCAAGAGTTACACCCAGGAGACGTACTTAAGTAAACACATGAACAAACACGCTGAAAGAGCTGACAAGAGGCCGCCGATCTCCGCGTTAGGACTGAGCGGGCTCAACAGGTCCTTAGGAGCAGCCGCGCCCACTGCGGCCCCTTTCGCGGATCACCCATATTGGCCTAAAGTTAGTCCGGATTCCGCCGCGCACATGTCTGATGAAGGCGGGTATCATCAGCAGAGGGAGAGCGACGATCACCATGAGCATCAACTGCAGCAGCAAAGGGCGCTGTTCGCTCAACACGAGAGTCAGGAAGACCGCATCCAGCCGCCGGTGTCGTCGGCTGCGAATTCCGCTTTCACGCCGATAAACTCCATGGCGCCTCATCTCAACGGTCTCTCGCATCACAGCGCTCTGCCCACGCGGCCGTATCTCTACGACCCGTTGCACTTCCAGCAAGGGAAGCAGCAGCCTAACTCCTTCCCCAATCAGCTGATATCGCTCCACCAAATCCGGAACTACGCGCACCAGCCGTCCGCCCTGTTGCCTGCCGAGCACATCCTCCCCCATTCCCTCTCTCACAAAGACAAGCAGTAA

Rn1_HCR_P1B3: gTCCCTgCCTCTATATCTTTTACGGACGGCTTTATCTCAGGCATG

Rn1_HCR_P2B3: ACCACTTTTGAATTCGGATAATCTATTCCACTCAACTTTAACCCg

Rn2_HCR_P1B3: gTCCCTgCCTCTATATCTTTGAATGTCATTCTCTTGGTCATTGTG

Rn2_HCR_P2B3: CATCGTCGTGCGGTGTCATTTGACGTTCCACTCAACTTTAACCCg

Rn3_HCR_P1B3: gTCCCTgCCTCTATATCTTTCCGTTTTCTGATAAAGGCGTTGAAG

Rn3_HCR_P2B3: ACCTTTTCCTCTTGAATGCTTTGTCTTCCACTCAACTTTAACCCg

Rn4_HCR_P1B3: gTCCCTgCCTCTATATCTTTGTGGATCTGCGTGTGCAGCTGGAGG

Rn4_HCR_P2B3: GCAGCGGTAGGGCTTCGCTTCGCGCTTCCACTCAACTTTAACCCg

Rn5_HCR_P1B3: gTCCCTgCCTCTATATCTTTGCGCTAGGTATGAGGAGTTGGCGAA

Rn5_HCR_P2B3: TGATGCCGAGGTGGATGCGCGAATGTTCCACTCAACTTTAACCCg

Rn6_HCR_P1B3:gTCCCTgCCTCTATATCTTTAGTTGGGTGAACTTGCGCTGGCAGA

Rn6_HCR_P2B3: GTGCGGATGTGCTGCTGCAGGTGAGTTCCACTCAACTTTAACCCg

Rn7_HCR_P1B3: gTCCCTgCCTCTATATCTTTGCACCCTATCTGGGTGCATCGATAG

Rn7_HCR_P2B3: GTTGGACAGCTGGGAGAACGCTTTGTTCCACTCAACTTTAACCCg

Rn8_HCR_P1B3: gTCCCTgCCTCTATATCTTTTGAAGGGTTTGTCGGTCTGGTGGCA

Rn8_HCR_P2B3: TGAAGCATTTGTAGCAGGAGTTGCATTCCACTCAACTTTAACCCg

***bric-a-brac* (*bab*)**

**>**bricabrac_LOC112043806_B2

Link: <https://www.ncbi.nlm.nih.gov/nuccore/XM_052890735.1>

ATGCCGGCCGAGGAAGACCCCGGGGCGATGGACGGCCGCGAAGCCGCCGACGGCTCGCCGCAGCAGTTCTGCCTGCGATGGAACAACTACCAGGCCAACCTGGCCACCTGCTTCGACCAGCTGCTGCAGACCGAGTCGTTCGTGGACGTGACGCTGGCGTGCGAGGGCCGCAGCCTGAAGGCGCACAAGGTGGTGCTGTCCGCCTGCAGCCCGTACTTCCAGACGCTGTTCACGGAGAACCCGTGCCGGCACCCGATCGTCATCATGCGCGACATCAAGTACTGCGACCTGAAGGCGGTGGTGGACTTCATGTACCGCGGCGAGATCAACGTGTCGCAGGACCAGATCGCGGCGCTGCTGCAGGTGGCGGAAACGCTGAAGGTGCGCGGGCTGACGGACGTGAGCGGCGAGCACGCCGTGCTGCGCGCCGCCAAGCGCAGCGGCACCGCGTCGCCGGACGCGCCGGGCGCGGGGCTGGCGGGCCCGGCCGGCAAGGCGCGCCGCCGCTCGGGCTCCGCGCCGCGCTCGCCGCCCGCCGACCTGCTGCACGCGCCGCCCGACGCGCCGCCGCCCATGCACGACGACGTCGACATCCGCCCCGGCATCGCCGAGATGATCCGCGAGGAGGAACGGGCCAAATTAATTGAAAACTCTCACGCTTGGCTCGGCGCCTCCACATCTTCGATTGCAGATAGCTACCAATATCAGCTGCAGTCCATGTGGCAGAAGTGCTGGAACACCAACCAGAGCATCGTGCACAATCTGAGGTTCCGCGAGCGCGGGCCGCTGAAGTCCTGGCGCCCGGAGACCATGGCTGAGGCCATATTCTCTGTTCTGAAGGAAGGGCTCTCACTGTCGCAGGCTGCGAGGAAATACGACATTCCGTACCCGACATTTGTGCTGTACGCTAACAGAGTCCACAACATGCTGGGCCCGAGCGCGGACGGTGGTGCAGATCTGAGACCCAAAGGTCGCGGGCGGCCGCAGCGCATTCTGCTGGGCGTGTGGCCCGACGAGCACATCCGCGGCGTCATCCGCGCCGTCGTGTTCCGCGACTCGCACGCCGCGCACGCCGTGCACGCCTCGCACCACATCAAGGAGGAGGTGCTCCCCGTCTACCCTGCGATCACGGACGGCATGACCTCGGTGCAGAACGCCTACCCGGTGGCGTGCGGCAACGGGCGCGAGGGCGGCGTGTCGCCCAACGCCGCGGCAGCCGCCGCCGTGGCCGCCGTGGCGCACGGGCTGCGCCGCCAGATGTGCAACATGGTCGTGGCCGCGCAGAGGCCGCCCGACCACCCGGTGCACCTCGGCCTGCCGCCGATGCCCATGCAGTCACCGCGGTTGCCATCGCCGCGCCTGCCGTCGCCGCTCACGGCCGGCTCGGAGTCGCCGCTGTGCTCGCCGCCGCCGCCGGGCCACGCCCTCGAGCCGCCGAAGCCCACCGACCCGTTCCGACCCTTCGCCTCGCCGCGCCCTGACAGCCGCTACAGCGACCGCGACGTCGTCGCCGACATGACCGACATGGGACCGCCTAAAACTCCGATTTCGATGGTGAAGAACGGCAAGGAGATGTCGCTGCCGGTGAAGATGGAGCTGCCGCCGGCGGAGGCGCGCGCGGAGAACATCTGA

***Lim homeodomain 1* (*Lim1*)**

>Lim1_LOC112054918_B2

Link: <https://www.ncbi.nlm.nih.gov/nuccore/XM_052883120.1>

ATGCCGACGTACGGAGGACGCGGCGAGGCGGGCGGCGCCATGGCGTGCGCCGGCTGCGAGAAGCCCATCCTGGACAAGTTCCTGCTGCACGTGCTGGAGCGCGCGTGGCACGCGGCGTGCGTGCGCTGCGCCGACTGCCGCGCGCCGCTCGCCGACAAGTGCTACTCCAGGGATAACAAACTCTTTTGTAGGAATGATTTCTTTAGACGTTACGGTACAAAATGCAGCGGCTGCGGTCACGGAATATCCCCATCAGATCTAGTTAGGAAAGCACGGGAAAAGGTGTTCCACCTGAATTGCTTCACCTGCTTAGTCTGCAGGAAGCAACTGTCCACTGGTGAAGAATTGTACGTGCTAGACGACAACAAATTTATATGTAAAGAGGATTACTTAGCGGGGAAGGCGCCGACGCATTCTGACTCACTACTCGGCTCAGGTTCAGATGACGAAGAAGAGGAAGAAGCGAGAGCAGCAAACAATTCAAGTAGTCCGGCGCATGCACCGCACCCTGCGCTACATTCTGAGCTATCACATAATGGGGACACCAAACCGCACGAAGATTCGGAAGATCAGGGTTCGCTGGACGGGGACACGGAGACCAGAGACTCTCAAGCGGAGAACAAGTCACCAGATGACGGCAACGGTGGCTCCAAAAGAAGAGGGCCTCGGACGACGATAAAGGCGAAACAGTTGGAGATACTGAAATCGGCTTTTTCACAGACACCCAAACCGACGAGGCACATACGAGAACAGCTGGCAAAAGAAACAGGGCTACCCATGAGGGTGATACAAGTCTGGTTTCAAAACAAGCGGTCAAAGGAACGGCGTCTAAAACAGCTGACGTCGATGGGCAGGGGGCCTTTCTTCGGCTCGTCGCGCAAGATGCGCGGCTTCCCCATGAACCTGTCCCCGGGGGGCCTGGAGGAGGGGCCGCCGGGGTTCCCATACTTCGCGACAGCAGATGGAAAGTTCGAATTTGGCTATGGCCCGCCGTTTCACCATGACGCGCCGTTCTTCCATCCGCCGCCCGCTATGCCGTTCAACCAGCCAGGTGGTATGGAGACGCTGCAAGGCGGCGAGTTCCAGGACCAGTTCCCTCATCACGAGCACCTGGTGCTGCCGCGGCCTTCTTCCCCCGAGTTCGCGTTCGGCGACGCCCCCCCTCCTCTCCACCCCGAGGGTCTGGTGTGGTAG

***fringe* (*fng*)**

>fringe_LOC112056524_B3

Link: <https://www.ncbi.nlm.nih.gov/nuccore/XM_024096965.2>

ATGAGTGACAAGGTGCGAAGACTTGGCCAAGATTATCAGCGCTGTGGCGCCGGCTCTGGCATGGGTGGGCGTAGGATAATCAAAGCTGCGGCGCTACTCATAGCTTTGGGATACTGCAGTTTGCTTGTTTATCAAGGCGGAGTGAATTACAACTTCCAAGAAAGTCGGCCAGTGCAAGTGGCGGACTTACCTATTGAGCCCGTGACGAAAACGAGTATTTTTGATAGTGGTCAGAGTAAGAGCATAACATTGGATGACATTTTCATAAGTGTAAAGACAACTAAGTTTTATCAGTATTCGAGGTTGCCGATAATTTTGAAGACATGGTTTCAATTGGCGAAAGAACAGACATGGTTTTTCACCGACACAGACAACCCACGACACCAAAATCAAACCAACGGTCACATGGTAAACACGAAGTGCTCGGCGTCACACCAGCTAAAACACCTCTGTTGCAAAATGTCCGTGGAGTACGATCATTTTCTCGAGAGCGGTAAAAAGTGGTTTTGTCACTTCGACGATGACAACTACGTCAATATTCCGCGCTTGGTCGCCGTGCTGCAGAAGTACAATCATCAAGAAGACTGGTATCTCGGGAAAACTTCTGTGAACAAACCTGTAGATATGTTCAAGAAGCCTACCAACGAGTTATTATACACTTTTTGGTTCGCAACCGGTGGTGCCGGAATATGTCTAAGCAGGAGTTTAGCTTTAAAAATGTTACCGATTGCAAGTGGTGGCAGATTCATCAGTATCTGTGAGAGTATTCGATTTCCGGATGATGTCACTTTGGGTTATATAATAGAACATCTAATGGAGAAAAACTTGACCCGATTGCCAGAACTACATTCACATTTGGAGCAAATGAAACTACTGAATCCTGAAACTTTTCGAGACCAAATATCGTTCAGTTACTCAAAAACAGCCAAAGGTTGGAACGTCATCAATGTTCCCGGATTCAACCATAGATATGATCCCACTAGGTTCCTCTCGCTGCATTGTTTCCTCTTTCCACACTTCAAATTTTGTCCGAGATAG

***decapentaplegic* (*dpp*)**

>decapentaplegic_LOC112044871_B2

Link: <https://www.ncbi.nlm.nih.gov/nuccore/XM_052883655.1>

ATGCGTGGGGCGTGCGCGTGCGCGGTGGTGTGCGCGTTGGTGGCGCTGTGCGCGGCGCGGCTGGACGAGTCCGCGCGCGCCGCCGCCGAGAAGCAGCTGCTGGCACTGCTGGGCCTGCCGCGCCGGCCGCCGCCGCGCGCCCGCCCGCCGCCCGTGCCGCGCGCGCTGCGCGTGCTGTACGACTCGCGCGCGCTGCCCGCCGCCGCCGCCAACACGGCGCGCTCCTTCCACCACACGCCCACGCCGCTCGACGAGCGCTTCCCCGGCGACCACCGCTTCCGCCTGTTCTTCAACGTGAGCGGCGTACCGGCCGACGAGGTGGCGCGCGGCGCCGACCTCTCGTTCCAACGAGCCGTCGGCACCACCGGCAGACAGAGACTGTTGTTGTACGACGTGGTGCGCCCCGGCCGCCGCGGCCACTCCGAGCCGATCCTGCGGCTGCTGGACTCCGTGCCGCTCCGGCCCGGGGAGGGAATCGTCAACGCCGACGCTCTGGGAGCGGCGCGACGGTGGCTCAAAGAGCCCAAACATAATCACGGACTATTAGTGCGAGTGTTAGAAGAAGACGCCGCGAGTGTGAGCAGGGACGCGAAGTTCCCGCACGTGCGCGTGCGCAGACGCGTCACGGACGAGGAGGAGGAGTGGCGGACGGCGCAGCCGCTGCTCATGCTGTACACGGAGGACGAGCGCGCGCGCGCGTCGCGGGAGACGAGCGAGCGGCTGACGCGCAGCAAGCGCGCGGCGCAGCGGCGGGGGCACCGCGCGCACCACCGCCGCAAGGAGGCGCGCGAGATCTGCCAGCGCCGCCCGCTGTTCGTCGACTTCGCGGACGTGGGCTGGAGCGACTGGATCGTGGCCCCGCACGGCTACGACGCGTACTACTGCCAGGGCGACTGCCCCTTCCCGCTGCCGGACCACCTCAACGGCACGAACCACGCGATAGTGCAGACTCTGGTCAACTCAGTGAACCCCGCGACGGTGCCCAAAGCGTGCTGCGTGCCGACGCAACTCTCATCTATATCTATGTTATATATGGACGAAGTGAACAATGTGGTGCTTAAAAACTATCAGGACATGATGGTGGTAGGCTGTGGCTGCCGATGA

#### **Appendix D: HCR Overview**

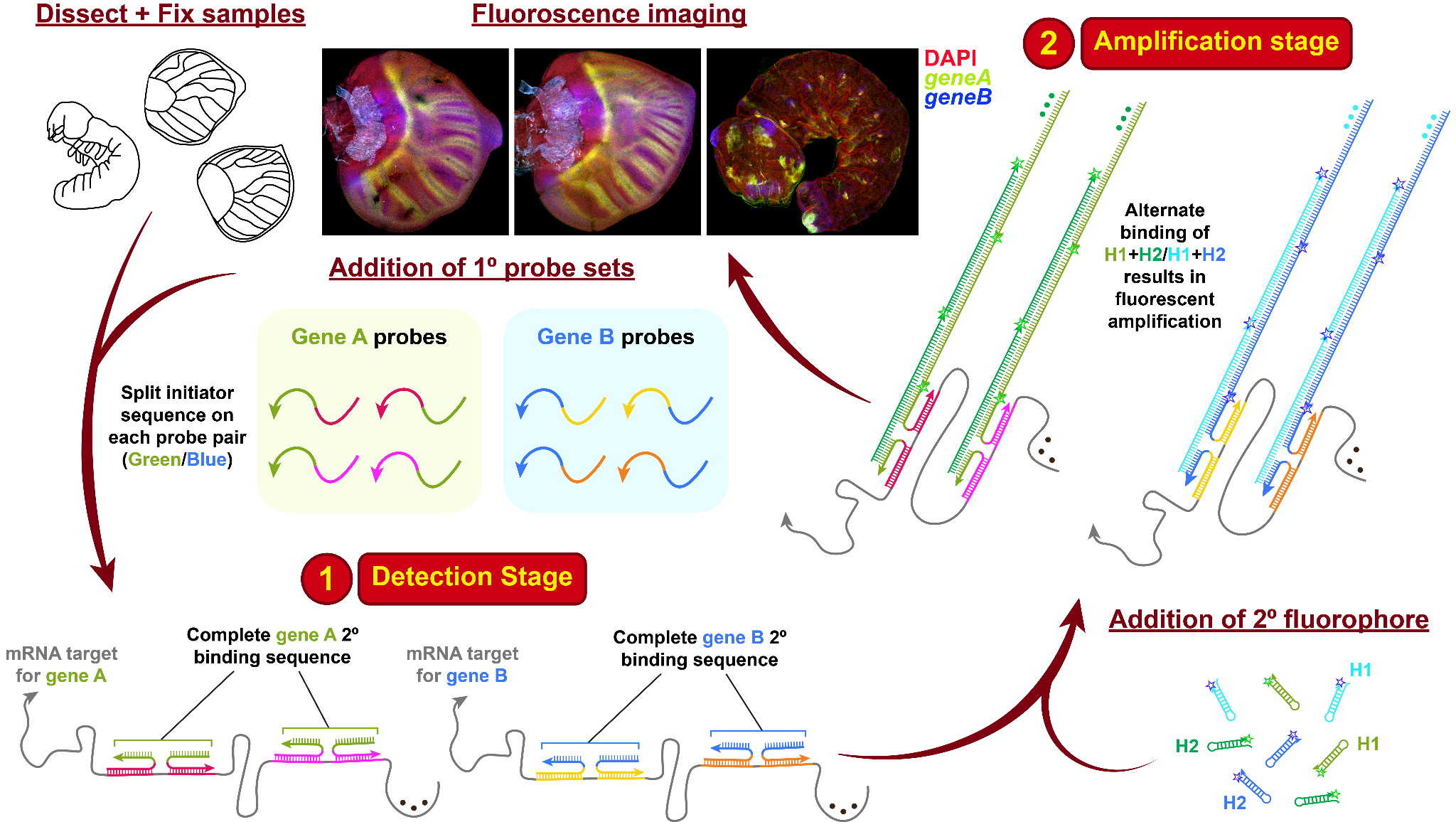

Figure S4: **Hybridisation chain reaction v3.0 overview (adapted from Choi *et al*., 2018).** The two main principles of this technique are (1) Detection and (2) Amplification, using split initiator 1° probes with different H1+H2 2° fluorescent hairpins for improved background suppression and high resolution multiplexing.
